# Spin-Coupled Electron Densities of Iron-Sulfur Cluster Imaged by *In Situ* Serial Laue Diffraction

**DOI:** 10.1101/2023.01.09.523341

**Authors:** Zhong Ren, Fan Zhang, Weijia Kang, Cong Wang, Heewhan Shin, Xiaoli Zeng, Semini Gunawardana, Kalinga Bowatte, Norbert Krauß, Tilman Lamparter, Xiaojing Yang

**Affiliations:** Department of Chemistry, University of Illinois at Chicago, Chicago, IL 60607, USA; Department of Ophthalmology and Vision Sciences, University of Illinois at Chicago, Chicago, IL 60607, USA; Renz Research, Inc., Westmont, IL 60559, USA; Botanical Institute, Karlsruhe Institute of Technology, Karlsruhe, Germany

**Keywords:** DNA photolyase, dynamic crystallography, metalloprotein, quantum crystallography

## Abstract

Iron-sulfur clusters are inorganic cofactors found in many proteins involved in fundamental biological processes including DNA processing. The prokaryotic DNA repair enzyme PhrB, a member of the protein family of cryptochromes and photolyases, carries a four-iron-four-sulfur cluster [4Fe4S] in addition to the catalytic cofactor flavin adenine dinucleotide (FAD) and a second pigment 6,7-dimethyl-8-ribityllumazine (DMRL). The light-induced redox reactions of this multi-cofactor protein complex were recently shown as two interdependent photoreductions of FAD and DMRL mediated by the [4Fe4S] cluster functioning as an electron cache to hold a fine balance of electrons. Here, we apply the more traditional temperature-scan cryo-trapping technique in protein crystallography and the newly developed technology of *in situ* serial Laue diffraction at room temperature. These diffraction methods in dynamic crystallography enable us to capture strong signals of electron density changes in the [4Fe4S] cluster that depict quantized electronic movements. The mixed valence layers of the [4Fe4S] cluster due to spin coupling and their dynamic responses to light illumination are observed directly in our difference maps between its redox states. These direct observations of the quantum effects in a protein bound iron-sulfur cluster have thus opened a window into the mechanistic understanding of metal clusters in biological systems.

## Introduction

Iron-sulfur clusters are redox cofactors ubiquitously found in biological systems. Ligated to a protein through the sulfur atoms of multiple cysteines, iron-sulfur clusters may contain two, three, four, or eight metal centers with the cubane type of four-iron-four-sulfur cluster [4Fe4S] being the most popular one (Beinert, 2000; Fontecave, 2006; Johnson et al., 2005). Owing to their structural and electronic versatility, these clusters are ubiquitous in life to carry out electron transfer and storage functions with or without a coupled proton transfer, thus mediate a variety of important physiological processes from nitrogen fixation, photosynthesis, cellular respiration, to DNA processing. More recently, the essential role of the iron-sulfur cluster is emerging among eukaryotic DNA processing enzymes, such as primases (Liu and Huang, 2015; O’Brien et al., 2017), replicative DNA polymerases (Baranovskiy et al., 2018; Jozwiakowski et al., 2019; Netz et al., 2012), and more broadly, many enzymes involved in nucleic acid metabolism and regulation (Khodour et al., 2019).

A new class of bacterial photolyases responsible for photoreactivation of damaged DNA by ultraviolet (UV) light irradiation was found containing a [4Fe4S] cluster in the C-terminal domain in addition to the usual catalytic cofactor flavin adenine dinucleotide (FAD) and a second chromophore 6,7-dimethyl-8-ribityllumazine (DMRL; Fig. S1) (Dikbas et al., 2019; Geisselbrecht et al., 2012; Marizcurrena et al., 2020; Oberpichler et al., 2011; Zhang et al., 2013). This is not surprising since flavin and iron-sulfur clusters are the usual redox partners (Vanoni, 2021). One member of this class PhrB from *Agrobacterium fabrum* specifically repairs the pyrimidine-pyrimidone 6-4 photoproducts of UV irradiation with a binding affinity to its substrate nearly 2000-fold higher than that to undamaged DNA (Graf et al., 2015; Zhang et al., 2013). PhrB shares the conserved protein architecture of cryptochrome/photolyase superfamily (Fig. S1) (Kavakli et al., 2019). Two electrons are required to reduce the fully oxidized FAD to activate photolyases, which process may have occurred under the reducing condition *in vivo*, instead of a light-dependent reduction (Sancar, 2003). Another electron is transferred to the DNA lesion site upon photoexcitation of the fully reduced hydroquinone FADH^-^ for the repair reaction (Zhang et al., 2017).

We recently proposed a photoreaction scheme in which the three cofactors of PhrB are interdependent. We suggested that the ribolumazine DMRL in PhrB could play a photoprotective role since it was found actively reverting the flavin to its fully oxidized, catalytically incompetent form FAD while itself is photoreduced under a prolonged light illumination (Ren et al., 2022). This process is again prominently displayed here in Fig. S2. We also hypothesized that the [4Fe4S] cluster plays a role of electron cache that absorbs excess electrons and supplies them back when needed, since two engaged photoreductions of FAD and DMRL favoring blue and violet light absorbing respectively could result in some electron imbalance under varying light intensity and wavelength composition. This reaction scheme explains the observed divergence and fluctuation in the absorption signature of the [4Fe4S] cluster (Figs. S2, S3, and S4).

In this work, we conduct crystallographic cryo-trapping experiments at low temperature and serial Laue diffraction from thousands of PhrB crystals at room temperature. We directly observe the electron density changes in the [4Fe4S] cluster of PhrB due to the redox responses of FAD and DMRL to light illumination. The electron density differences with respect to the resting valence state of [4Fe4S]^2+^ in PhrB (Bauer et al., 2014) are experimentally captured in this study (Fig. 1). These light-induced changes in the metal cluster cannot be explained by a displacement or deformation of the cluster, commonly known as a conformational change that requires atomic motions. Instead, the captured changes in electron densities exhibit a variety of complex features only obtained by quantum chemistry calculations in the past (Beinert, 2000; Noodleman and Case, 1992; Voityuk, 2010). These observed electron density features agree well with the theoretical prediction. Therefore, we conclude that these features are due to movements of electrons, that is, redox changes, and fundamentally caused by the interaction between nearby metal centers known as coupling of electron spin, the fourth quantum number that parameterizes the quantized intrinsic angular momentum of electrons.

**Figure 1.**
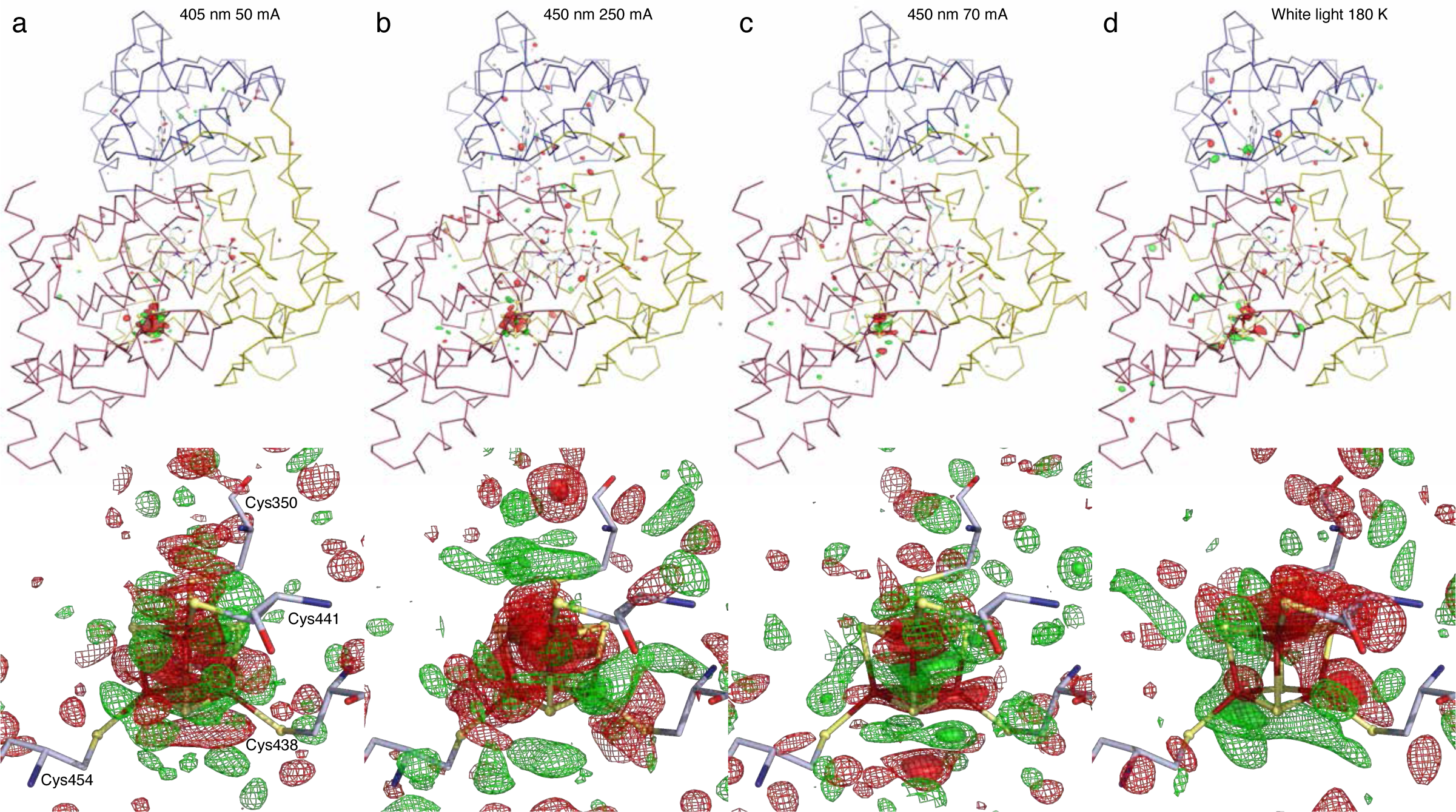
Dissimilar examples of raw difference Fourier maps. Light-minus-dark difference maps are calculated from datasets collected at room temperature (a-c) and cryo temperature (d). Strong difference signals up to ±12*σ* are concentrated at the [4Fe4S] cluster only, where *σ* is the standard deviation of the difference electron density over the entire protein molecule. These difference maps indicate that the overall signal-to-noise ratio is significant. Positive and negative electron densities above the noise level in green and red demonstrate electron density gain and loss, respectively. Illumination conditions are labeled. See Fig. S4 for corresponding power densities. (a) The large map over the entire protein molecule is contoured at ±5*σ* in green and red, respectively. The zoom-in view at the [4Fe4S] cluster is contoured at ±3*σ* in mesh and ±8*σ* in surface. (b) The large map is contoured at ±4*σ*. The zoom-in view is contoured at ±2.5*σ* in mesh and ±4.5*σ* in surface. (c) The large map is contoured at ±4*σ*. The zoom-in view is contoured at ±2.5*σ* in mesh and ±4*σ* in surface. (d) The cry temperature map is contoured at ±4*σ*. The zoom-in view is contoured at ±2*σ* in mesh and ±3.5*σ* in surface.

Briefly, a ferric ion Fe^3+^ and a ferrous ion Fe^2+^ carry five and six 3*d* electrons, respectively. According to Hund’s rule, the five 3*d* orbitals of a ferric Fe^3+^ are exactly half filled with electrons of the same spin. A ferrous Fe^2+^ would have only one of the 3*d* orbitals doubled with an additional electron of the opposite spin. When multiple metal centers cluster together, the weak metal-metal interactions tend to form antiferromagnetic alignment among the metals, known as spin coupling. As a result, the majority spin of two irons in a [4Fe4S] cluster is antiparallel to that of the other two according to the two-layer model (Beinert, 1997; Noodleman et al., 2002, 1995). When ferric and ferrous ions are mixed in the same cluster, electrons are delocalized in these layers, which effectively results in a mixed valence layer of two Fe^2.5+^ (Fig. 2c). The layered arrangements of some redox states in [4Fe4S] cluster due to spin coupling are directly observed in our maps of electron density changes upon light illumination. Although redox reactions of [4Fe4S] clusters involve only valence changes of the iron atoms, according to quantum chemical theory, the actual charge distribution changes significantly around the sulfur atoms, including the Sγ atoms of the Cys anchors, known as electron relaxation of the passive orbitals (Noodleman et al., 1995; Noodleman and Case, 1992). Our electron density maps clearly exhibit the effect of electron relaxation contrary to a potential deception that an electron gain or loss centers only on the iron atom where a valence change occurs. We are unaware of any previous report on a direct imaging of such quantum effect on the electron density of metal centers in protein. Observations of such layered electronic structure and its dynamic behavior could offer a direct insight into the functional role of these metal clusters in proteins, such as their magnetic dipole moment. In general, direct imaging of quantum effects in protein bound metal clusters could enable an entirely novel approach to investigate elusive functional mechanisms of metals in biological systems (Holm et al., 1996).

**Figure 2.**
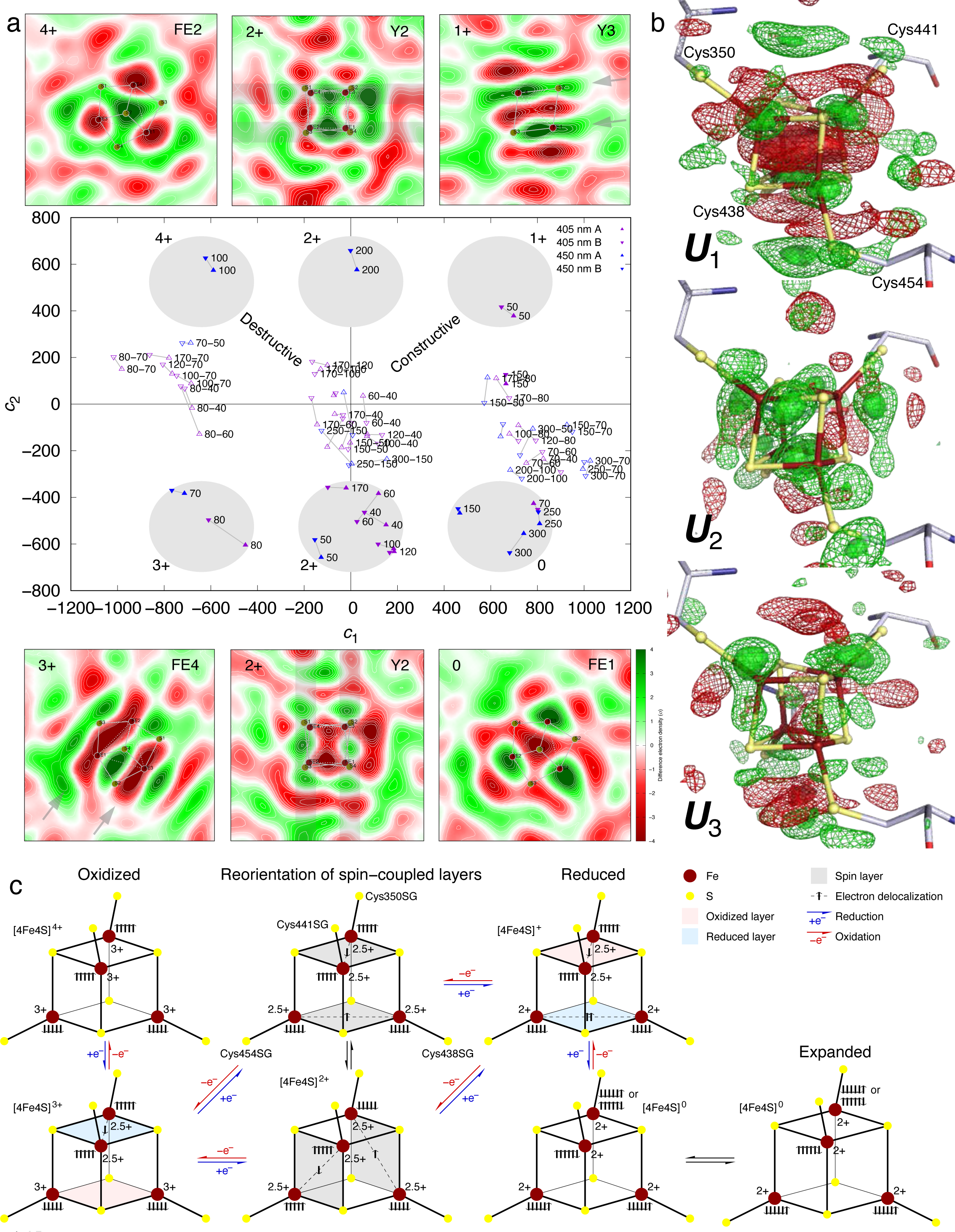
Overview of spin-coupled redox responses at [4Fe4S] cluster. Light datasets are collected after a 2-second illumination of violet or blue light emitted from laser diodes at various power densities (Methods). Dark datasets are collected without illumination. Each light-minus-dark and light-minus-light difference map is represented as a linear combination of several major components *c*_1_***U***_1_ + *c*_2_***U***_2_ + *c*_3_***U***_3_ + … derived from an SVD analysis (Methods). Each component map ***U****_k_* (*k* = 1, 2, 3, …) captures the common features among the raw difference maps (Fig. 1). (a) The center panel shows a scatter plot of the coefficients *c*_1_ and *c*_2_ for the top two components of the difference maps at room temperature. See Fig. S12 for the orthographical views of the coefficients *c*_1_, *c*_2_, and *c*_3_ for the first three components. Light – dark and light – light difference maps are marked with solid and open triangles, respectively. Maps from two chains A and B related by the noncrystallographic symmetry are in up- and down-pointing triangles and linked with a gray line. The electric current setting of the laser diode is labeled wherever possible. See Fig. S4 for the corresponding power density of illumination. The light – dark maps in solid triangles are divided into six groups shaded by gray ellipses, while the light – light maps in open triangles are in three groups near the axis of *c*_2_ = 0. This study presents evidence to support that each group of the light – dark maps depicts a valence state of the [4Fe-4S] cluster from 0 to 4+ as marked by the gray ellipses. Above and below the center panel, six square cross sections of difference maps highlight the most striking features of the corresponding valence state (see main text). See Figs. S1 and S13 for the orientation of each cross section. (b) First three map components ***U***_1_, ***U***_2_, and ***U***_3_. Each map is contoured at ±2.5*σ* in green and red mesh and ±4.5*σ* in green and red surfaces. +Z direction approximately aligned with helix X toward its C-terminus is oriented upright (Fig. S1). C, N, O, S, and Fe atoms are in white, blue, red, yellow, and rusty colors, respectively. See Figs. S13-S16 for detailed plots of various cross sections through these components and the seventh component. In the first component ***U***_1_, positive densities up to 7*σ* are on four irons (Fig. S13). Weaker positive peaks around 3*σ* are associated with each of the eight sulfurs, including four from the Cys residues. Another shell of positive densities forms an incomplete cubic shape of 7.1 Å from side to side. Each face of the cube carries a stronger density up to 4-5*σ* and the edges and corners of the cube are broken. The positive cube is visible from the cross sections X2, Y2, and Z2 in Fig. S13. In the second component ***U***_2_, 12 positive peaks form a 3×2×2 array in X, Y, and Z directions, respectively. The three 2×2 layers are normal to X axis with the middle layer going through the center of [4Fe4S] (Fig. S14). Two of the center peaks are fused together (X2 and Z3 in Fig. S14), and two of the corner peaks are weakened to almost none (Y1, Y3, and Z1 in Fig. S14). Weak negative densities up to -2.5*σ* in four layers flank and interleave three positive layers (Y1-Y3 and Z1-Z3 in Fig. S14). As a result, all positive and negative peaks miss the atomic positions of the irons and sulfurs. The 3×2 arrays normal to Z axis are coplanar with two atomic layers in [4Fe4S] (Y1-Y3 in Fig. S14). This is also seen as arrays of six positive densities in the top mid and bottom mid Y2 cross sections shaded in gray (a). The other 3×2 arrays normal to Y axis are slightly misaligned with the atomic layers (Z1-Z3 in Fig. S14). The third component ***U***_3_ is similar to ***U***_2_. However, a 2×2×3 array of positive peaks is arranged in X, Y, and Z directions, respectively. Two strong positive peaks at 6.5*σ* are located between two irons and two sulfurs of Cys350 and 441. Both skew toward the irons (YX in Fig. S15). The equivalent peaks associated with Cys438 and 454 are weaker at 4*σ* (XY in Fig. S15). The seventh component ***U***_7_ displays some additional layering structure (Fig. S16). (c) Schematic summary of spin-coupled redox responses of the [4Fe4S] cluster in PhrB. 3*d* electrons of the irons are marked as up or down arrows to differentiate their spin momenta. These drawings are arranged in the same way as the six shaded groups in (a) except an additional, bottom rightmost drawing depicting the expanded cluster. The second column depicts an equilibrium between two different orientations of valence layers in the dark resting state. This does not exclude the possibility for valence layers oriented in the third way also mixed in the equilibrium in dark. However, we capture no clear signal of electron transfer involving the third orientation in the top 10 component maps.

## Results

### Dynamic crystallography experiments

To capture light-induced electron density changes in PhrB, we adopt two different technologies in protein crystallography at cryo and room temperatures. However, the common spirit in dynamic crystallography is the same, that is, a direct contrast of many experimental conditions in light and dark and a joint analysis of multiple datasets using singular value decomposition (SVD) to dissect the common features among datasets under variable conditions (Ren et al., 2013). Our dynamic crystallography observations at both cryo and room temperatures uncover unique structural insights into the redox role of the [4Fe4S] cluster in PhrB. More importantly, the approaches presented here provide a set of experimental and analytical tools to study the complicated redox responses in metalloproteins.

At room temperature, our observations of the movements of individual electrons are obtained from *in situ* Laue diffraction of thousands of crystals grown on crystallization chips that require no crystal harvesting and manipulation thus offer very stable redox condition. We have recently developed an automated serial platform inSituX for *in situ* Laue diffraction at room temperature that facilitates such observations (Biju et al., 2022; Ren et al., 2020). Here the phrase “*in situ* diffraction” is strictly defined as shooting crystals in their original place where they grew without ever breaking the seal of the crystallization chamber, that is, a procedure devoid of sample mounting, loading, transfer, drying, air exposure, or any manipulation, except taking micrographs of the crystals under infrared (IR) light. PhrB crystals are grown on crystallization devices (Fig. S5) that are made of monocrystalline quartz (Ren, 2017; Ren et al., 2018). These on-chip crystals are imaged under IR to avoid photoactivation and recognized by an image analysis software developed as part of inSituX. They are then automatically centered in a polychromatic X-ray beam for Laue diffraction (Ren et al., 1999) according to a preprogramed sequence (Fig. S6). A light illumination on the centered crystal precedes the X-ray exposure for light datasets (Fig. S3a). Serial Laue diffraction experiments (Methods; Figs. S7-S9) are conducted in three separate runs at BioCARS beamline 14-ID-B of Advanced Photon Source (Graber et al., 2011). A dark dataset is collected in each run to ensure the best reference dataset. Light datasets are collected after a two-second illumination of the crystals with wide ranges of power density at a wavelength of 405 nm in violet and 450 nm in blue (Table S1; Figs. S3a and S4; Methods). A total of 29,296 Laue images are collected *in situ* from 69 crystallization devices. Each image is derived from a fresh crystal previously unexposed to X-rays. Diffraction data reduced from 14,002 images are merged into three dark and 16 light datasets under different illumination conditions (Table S1). Difference Fourier maps are calculated among datasets from the same run, including 32 light-minus-dark and 66 light-minus-light difference maps from both protein chains in the asymmetric unit, according to previously established protocols. These 98 difference Fourier maps are subjected to an SVD analysis (Methods) (Ren, 2019; Ren et al., 2013). Strong difference signals concentrated at the iron-sulfur cluster differ markedly from those more familiar features caused by atomic displacements that often display positive-negative pairs of electron densities (see Figs. 6b and S10 for examples of atomic displacement). These prominent signals on the [4Fe4S] cluster capture the light-induced transitions among various valence layers of [4Fe4S] (Fig. 1abc). Only slight signals are found along helix P, to which the iron-sulfur cluster is ligated (Fig. S1). However, no difference densities are found above the noise level in and around FAD or DMRL cofactors (Fig. 1) although single-crystal spectroscopy shows significant spectral change associated with the photoreductions of these redox-active chromophores (Fig. S2).

Our temperature-scan experiments are conducted at elevated cryo temperatures between 100 and 200 K while the crystal is illuminated under white light (Figs. S3b and S4) (Bandara et al., 2017; Shin et al., 2019; Yang et al., 2011; Zeng et al., 2015). We collect a total of nine datasets after white light illumination. Each of these light datasets corresponds to a dark dataset collected from a different segment of the same crystal (Table S2). Multiple dark datasets are also collected from the same crystal segment at 100 K after X-ray irradiation of variable dosages. Therefore, the difference maps produced from these datasets serve as controls to reveal the effect of X-ray reduction. 18 light-minus-dark difference maps and eight high X-ray dose – low dose difference maps from both protein chains in the asymmetric unit are produced and subjected to an SVD analysis. These low temperature datasets also give rise to strong difference signals centered at the [4Fe4S] cluster. The layered electron densities resemble those obtained at room temperature, yet they directly contrast with the map features arising from X-ray irradiation in the control experiments. Additional signals that are not found in room temperature Laue data are also present on the [4Fe4S] cluster and a few surrounding helices that show their concerted motions (Fig. S10).

### Quantized signals in difference maps

Crystallographic observations are sensitive to conformational changes in proteins and chromophores that involve atomic displacements. However, the extensive spectral changes we reported recently can be interpreted as redox reactions that involve relocation of electrons but not necessarily any displacement of atoms (Ren et al., 2022). The only indication for possible protein motion is absent in crystals. Direct observations of the movements of a few individual electrons in electron density maps of proteins or even smaller molecules are theoretically possible but were reported rarely (Ren, 2019; Velazquez-Garcia et al., 2022; Zuo et al., 1999).

98 difference maps from Laue diffraction at room temperature are masked within a 4-Å radius of the [4Fe4S] cluster and its four Cys anchors and then subjected to an SVD analysis that decomposes the observed maps into several component maps ***U***_1_, ***U***_2_, etc. that contain the common features among the 98 observed maps (Fig. 2b). Each observed map consists of a different amount of the component maps, which is presented as a set of coefficients *c*_1_, *c*_2_, etc. The coefficients corresponding to the first two most significant components form six discrete groups with no obvious correlation to the wavelength or power density of the light illumination. Light – dark difference maps (solid triangles in Fig. 2a) are clustered around *c*_1_ = 0 and ±640; *c*_2_ = ±525. Light – light difference maps (open triangles in Fig. 2a) also form three discrete groups around *c*_2_ = 0 that are separated from the light – dark groups. That is to say, difference maps with a dark reference take a form of {0,±1}640***U***_1_ ± 525***U***_2_, known as reconstituted maps that approximate the raw difference maps in a least-squares sense (Methods), where ***U***_1_ and ***U***_2_ are two most significant component maps (Figs. 2b, S13, S14). This paper presents the component maps, and many observed raw difference maps, along with the reconstituted maps that best represent the discrete groups in three-dimensional contours and two-dimensional cross sections. Table S3 summarizes the figures and figure parts that present various maps and serves as a directory for easy lookup.

The surprising discreteness revealed by the distribution of the SVD coefficients strongly suggests that our observed difference electron density maps depict relocations of a few individual electrons, that is, quantized orbital transitions. In other words, although the diffraction data are collected from thousands of crystals (Table S1), the light-induced changes in each protein molecule among a large ensemble of molecules involve either none or only a very small number of the same types of individual orbital transitions. These transitions may also include a reorientation of the valence layers without changing the total charge of the cluster (see below). The light – light difference maps are also quantized in *c*_1_ direction, that is, the difference between two light responses is also discrete as a discrepancy that involves only one or very few electrons (Fig. 2a). But they differ from the light – dark difference maps in *c*_2_ dimension.

As controls, we also produce three dark-minus-dark maps from the room temperature Laue datasets in the same way (Fig. S11). It seems that a low level of X-ray induced redox changes could exist at the [4Fe4S] cluster. Nevertheless, compared to the light-minus-dark maps (Fig. 1), we conclude that the quantized transitions of a few individual electrons at the [4Fe4S] cluster are mostly caused by two entangled photoreductions of the FAD and DMRL cofactors rather than the X-ray induced reduction of the [4Fe4S] cluster. It is entirely possible that some weak signals of X-ray reduction at the [4Fe4S] cluster also involve a small number of electrons, therefore, indistinguishable from the changes induced by visible light (see below for cryo data).

### Electron relaxation

The first component ***U***_1_ of the difference maps is nearly twice as significant as the second, judged by their singular values (Fig. S12d). Positive densities up to 7*σ*are on four irons (Figs. 2b and S13), where *σ* is the standard deviation of the component map. These are shown as the deep green spots on the iron atoms in the cross sections of ***U***_1_ map (Fig. S13). A large negative peak is located at the body center of [4Fe4S]. Four negative pockets fused with the central negative peak wrap around the positive peaks on the irons. The red negative pockets and the central peak are obvious in ***U***_1_ cross sections. Each of the eight sulfurs, including four Sγ atoms from the Cys residues, is associated with a positive peak around 3*σ*. These positive peaks are not exactly centered on the sulfurs but skewed outward away from the center of [4Fe4S]. The positive peaks associated with the sulfur atoms in the [4Fe4S] cluster are clearly visible in the body diagonal cross sections S1-S4 and those associated with the Cys Sγ atoms can be seen in six corner diagonal cross sections such as XY, YZ, etc. (see Figs. S1 and S13 for definition of coordinate system and cross sections). ***U***_1_ is nearly symmetric among X, Y, and Z axes, known as full tetrahedral symmetry T_d_, up to the available resolution in our crystallographic data. However, features along Z axis are stronger, which makes Z axis unique compared to X and Y axes, thus deviates slightly from the ideal tetrahedral symmetry (Fig. S13). Here we interpret ***U***_1_ as the overall effect of an addition of electron(s) into the system of [4Fe4S], that is, a reduction step. Therefore, the negation -***U***_1_ depicts an oxidation step. The weaker positive peaks associated with all eight sulfurs largely agree with the theory on electron relaxation of the passive orbitals (Noodleman et al., 1995; Noodleman and Case, 1992). That is to say, a valence change nominally at one of the irons would cause a redistribution of the electrons on all four irons and on the eight sulfurs including those from the Cys anchors. The top component of the net change of electron density distribution experimentally captured here largely resembles the theoretical calculation of difference electron density maps concerning electron relaxation (Beinert, 2000; Noodleman and Case, 1992; Voityuk, 2010).

We further clarify that the most significant component map ±***U***_1_ depicts the features of electron density redistribution as one or two electrons are added to or subtracted from the [4Fe4S]^2+^ cluster. However, no information is available on the amplitude of difference electron densities since none of our difference maps is calibrated to an absolute scale. Therefore, no map carries a physical unit, such as e^-^/Å^3^. As a consequence, an addition of one electron to the [4Fe4S] cluster results in the same coefficient *c*_1_ in +640***U***_1_ as an addition of two electrons (right column in Fig. 2a), because a difference map of a two-electron reduction does not display twice the electron density changes compared to a map of a one-electron reduction when the absolute scale is absent. However, one- and two-electron reductions or oxidations will lead to dissimilar molecular symmetries that are depicted by the second component ***U***_2_ (Fig. 2a; see below).

### Spin-coupled valence layers

The first component ***U***_1_ of the difference maps could carry a positive or negative coefficient depicting a reduction or oxidation step, respectively, if it is present in an observed difference map (Fig. 2a), which is consistent with our spectroscopic observations that a redox step of the [4Fe4S] cluster could go either way depending on the balance between the photoreductions of FAD and DMRL (Ren et al., 2022). It is interesting that ***U***_1_ usually combines with other components required by the light – dark maps in four corner groups, either constructively or destructively (Fig. 2a). The tetrahedral symmetry of the component map ***U***_1_ is broken by these combinations since the second component ***U***_2_ is reduced to a lower symmetry of C_2_ with the only 2-fold axis along Z (Z1-Z3 in Fig. S14). The most striking features in ***U***_2_ are 12 positive peaks up to 8*σ* arranged in a 3×2×2 array in X, Y, and Z directions, respectively (Figs. 2b and S14).

These 12 positive peaks are most obvious as a 3×2 array in ***U***_2_ cross sections of Y1, Y3, and Z1, Z3 (Fig. S14). The top layer connected to Cys350 and 441 is not symmetrical to the bottom layer connected to Cys438 and 454. This asymmetry in ***U***_2_ differentiates the top from the bottom layer, where the top and bottom layers Z1 and Z3 are perpendicular to Z axis and helix X (Fig. S1). This second component is added to or subtracted from the first one in nearly the same strength so that the first component is modified one way or the other. As a result, the direction along Z axis is further differentiated from X and Y axes when ***U***_1_ and ***U***_2_ are constructively combined together, that is, ±640***U***_1_ ± 525***U***_2_ depicting electron density changes from [4Fe4S]^2+^ to [4Fe4S]^+^ and to [4Fe4S]^3+^, respectively (constructive diagonal in Fig. 2ac). The raw difference maps with good signal-to-noise ratios show striking layers of difference densities normal to Z axis (Fig. 3ab). Gain and loss of a single electron from [4Fe4S]^2+^ are visualized in two layers of the iron-sulfur cluster shown as two dark green and dark red layers (marked by arrows in Y3 and FE4 in Fig. 2a). This observation agrees well with the two-layer model (Beinert, 1997; Noodleman et al., 2002, 1995; Wang et al., 2003). Here we abstract these two cases in the top right drawing for [4Fe4S]^+^ and bottom left drawing for [4Fe4S]^3+^ in Fig. 2c.

The third component ***U***_3_, nearly as strong as the second (Fig. S12d), further modifies the difference maps so that the ferric and ferrous layers are differentiated from the mixed valence layer. ***U***_3_ features two strong positive peaks at 6.5*σ* between two irons and two sulfurs of Cys350 and 441 in the top layer (Figs. 2b and S15). This can be seen as two dark green peaks in YX cross section of ***U***_3_ map (Fig. S15). ***U***_3_ also resembles ***U***_2_ with an array of 12 positive peaks but arranged as a 2×2×3 array in X, Y, and Z directions, respectively, which is most obvious as a 2×3 array of positive peaks in the ***U***_3_ cross sections X1, X3, and Y1, Y3 (Fig. S15). The intensities of these 12 peaks vary a lot (XY and YX in Fig. S15) and this array in ***U***_3_ appears larger than the array of 12 positive peaks in ***U***_2_ (see below). The coefficient of the third component does not appear to be quantized (Fig. S12ac). Linear combinations with ***U***_3_ constructively, that is, ±640***U***_1_ ± 525***U***_2_ ± *c*_3_***U***_3_ (*c*_3_ > 0), result in separate peaks for gain or loss of electron densities on four irons. However, the light – dark maps seem to favor a destructive combination (Figs. 3abc, S17, and S19), that is, ±640***U***_1_ ± 525***U***_2_ ∓ *c*_3_***U***_3_ (*c*_3_ > 0). A destructive combination shows that any gain or loss of electron density in the bottom layer centers on the cubic face (gray arrows in Z3 of Fig. 3cd), which seems to suggest that the delocalized electron in the bottom layer is lost if oxidized to [4Fe4S]^3+^ (Fig. 2c bottom left) and that a gained electron is also delocalized in the bottom layer if reduced to [4Fe4S]^+^ (Fig. 2c top right). On the other hand, electron density changes in the top layer are divided between two irons. In addition, the positive or negative densities associated with each iron in the top layer are further split into two peaks (gray arrows in Z1 of Fig. 3cd). As a result, the destructive combinations of ***U***_3_ mark the bottom layer as the ferric layer in [4Fe4S]^3+^ or the ferrous layer in [4Fe4S]^+^ that is differentiated from the top layer as the mixed valence layer after a redox step (Fig. 3ab). Despite the strongly layered features in the difference maps (Figs. 3 and S17-S20), it is not clear whether and how the protein environment contributes to the selection of the valence layers, either temporarily or permanently. If the valence layers are randomly distributed among X, Y, and Z directions, it would be difficult to observe the layered electronic structure. Therefore, we presume that the protein environment plays a role.

**Figure 3.**
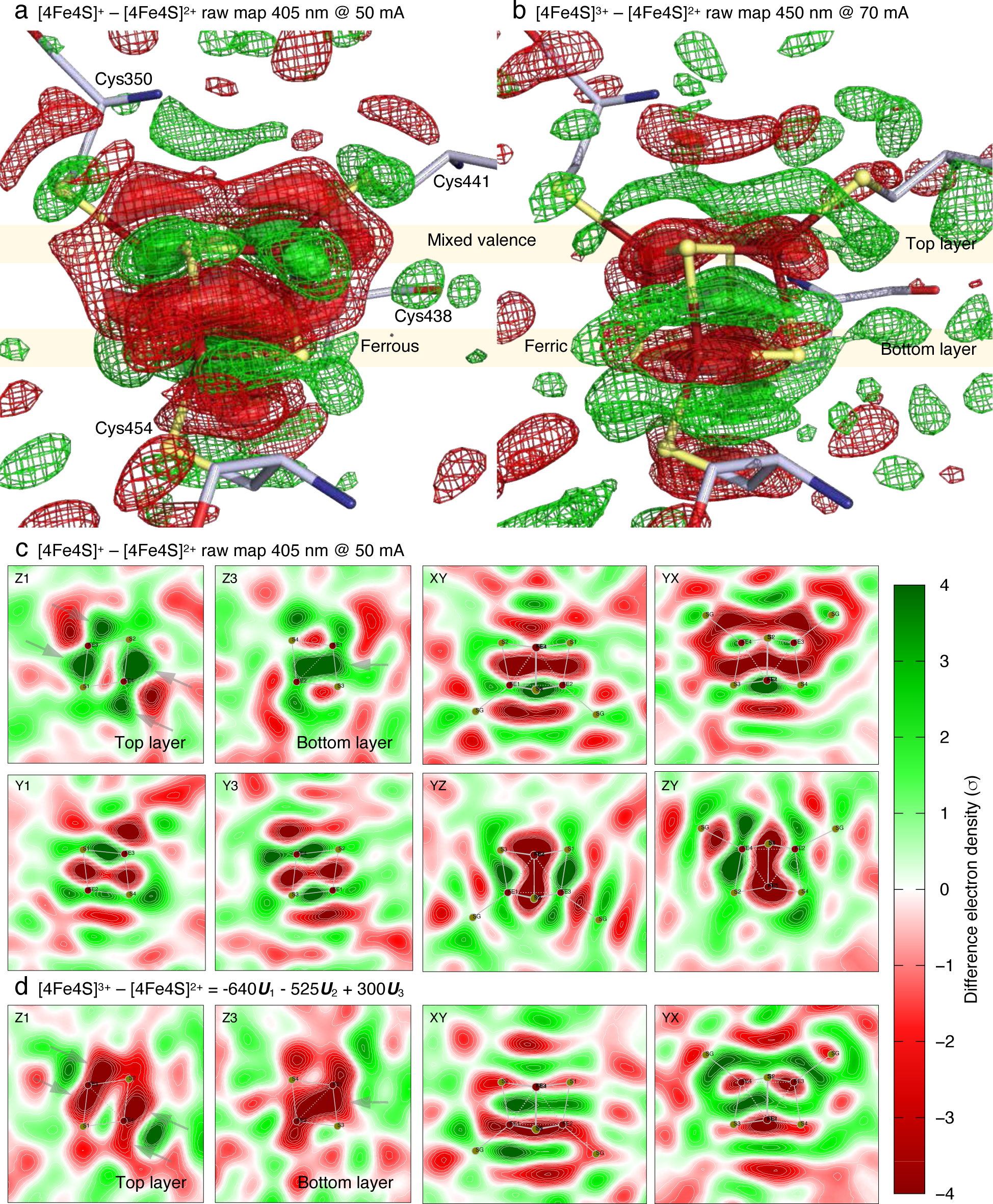
Direct observations of valence layers. (a) Difference map derived from violet light illumination of 405 nm at 50 mA (Fig. S17). This raw difference map prior to SVD is contoured at ±2*σ* in green and red mesh and ±4*σ* in green and red surfaces. (b) Difference map derived from blue-light illumination of 450 nm at 70 mA (Fig. S19). This raw difference map prior to SVD is contoured the same way as (a). See Fig. S4 for the corresponding power density of illumination. (c) Cross sections of (a). See Fig. S1 for definitions of the cross sections. Despite different signal-to-noise ratios, (a) and (b) are approximately the negation of each other. They are direct observations of [4Fe4S]^+^ – [4Fe4S]^2+^ and [4Fe4S]^3+^ – [4Fe4S]^2+^ differences, respectively. Strong features are in layers directly caused by spin coupling. In the top layer Z1, the electron gain or loss is split between two irons ligated to Cys350 and 441, while the electron gain or loss is centered on the cubic face in the bottom layer Z3. See Figs. S18 and S20 for detailed plots of the cross sections of the reconstituted difference maps that reproduce these observations. (d) Reconstituted difference map -640***U***_1_ - 525***U***_2_ + 300***U***_3_. This reconstituted map by linear combination reproduces the difference map in (b) except the noise (Fig. S20). The difference densities split between two irons in the top layer (Z1) but form a center peak in the bottom layer (Z3).

### Symmetric redox states

While the constructive combinations of ***U***_1_ and ***U***_2_ produce the layered structure in difference maps, destructive combinations ±640***U***_1_ ∓ 525***U***_2_ result in well separated peaks associated with irons and sulfurs without any obvious layers. ***U***_3_ again modifies the combinations and substantially restores the tetrahedral symmetry. We infer that the redox states have reached [4Fe4S]^0^ and [4Fe4S]^4+^ from the restored T_d_ symmetry absent of obvious layers. In the highly oxidized, all-ferric cluster [4Fe4S]^4+^, electrons of the minor spin are lost. All 3*d* orbitals of four ferric ions are exactly half filled (Fig. 2c top left). Therefore, valence layers disappear, and the observed difference map follows the tetrahedral symmetry (Fig. 2a top left). Electron density losses on four irons are distributed evenly compared to [4Fe4S]^2+^ in dark (Fig. 4a). This is only observed once from the illumination of 450 nm blue light at 100 mA (Fig. S21) that requires a destructive combination of -640***U***_1_ + 525***U***_2_ with a negative term of ***U***_3_ (Figs. 4a and S22). A crystallographic structure of such all-ferric cluster [4Fe4S]^4+^ determined previously showed that two Fe-Fe distances are 0.05 Å shorter than the other four (Moula et al., 2018). However, the tetrahedral symmetry holds here with the resolution available to this work.

**Figure 4.**
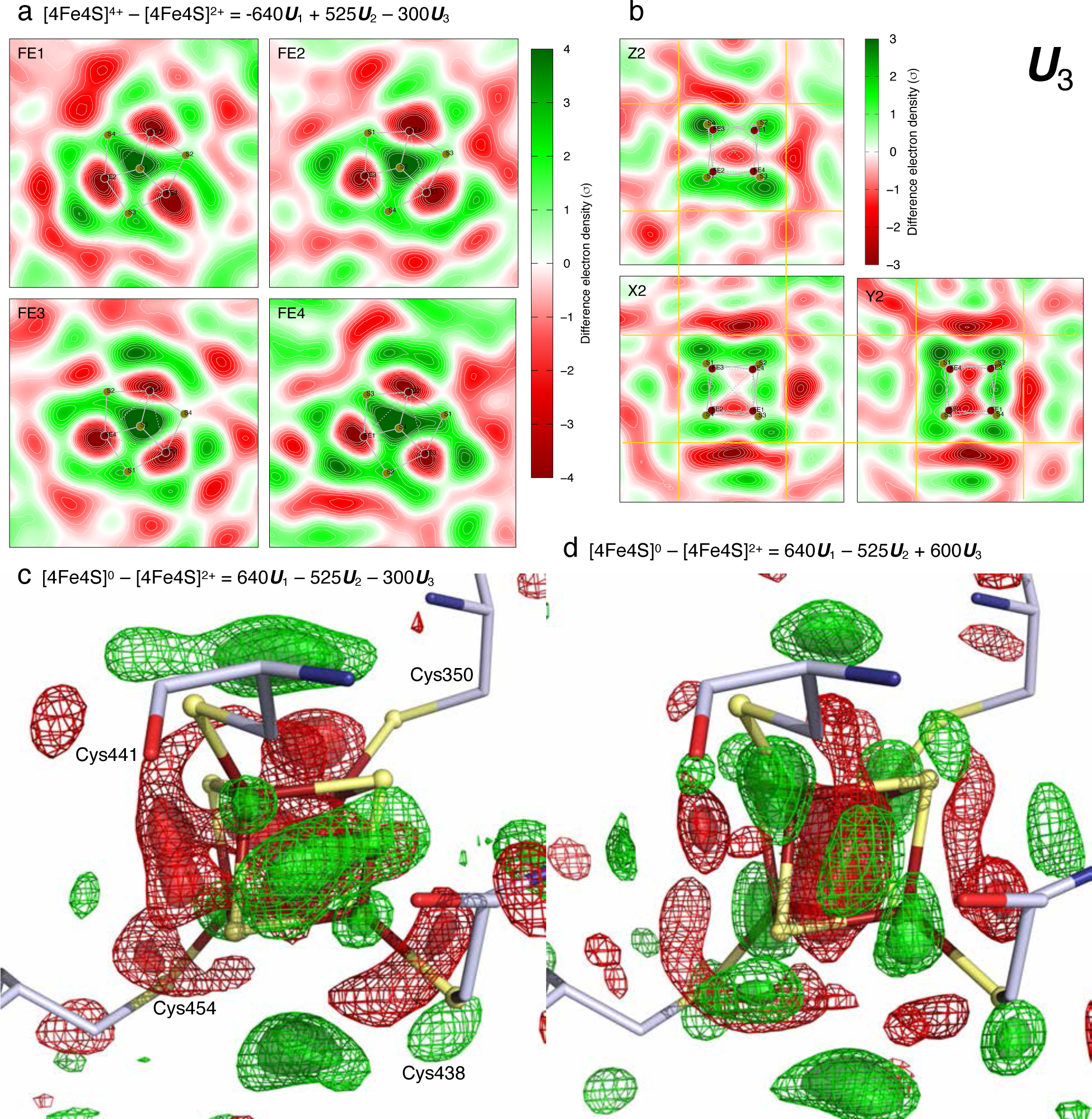
Tetrahedral symmetry restored in [4Fe4S]^0^ and [4Fe4S]^4+^. (a) Three-fold axes of all-ferric cluster [4Fe4S]^4+^. The reconstituted difference map -640***U***_1_ + 525***U***_2_ - 300***U***_3_ restores the tetrahedral symmetry T_d_ displayed in ***U***_1_ but not in the other components (compare Fig. S22 with Figs. S13-S15). A raw difference map of all-ferric cluster [4Fe4S]^4+^ is captured by illumination of blue light of 450 nm at 100 mA (Fig. S21). (b) Thick integrations through ***U***_3_. Three orthographical cross sections X2, Y2, and Z2 are displayed again with thick integrations ±2.3 Å around these central cross sections. The slabs integrated are shown between the yellow lines. These thick integrations demonstrate that positive densities of the third component distribute slightly outside the [4Fe4S] cluster (compare to Fig. S13). (c) and (d) Two extremes of all-ferrous cluster [4Fe4S]^0^. These reconstituted difference maps are contoured at ±2.5*σ* with mesh and ±3.5*σ* with surfaces. The positive and negative contours are in green and red, respectively. The reconstituted difference map 640***U***_1_ - 525***U***_2_ - 300***U***_3_ restores the tetrahedral symmetry and shows positive densities on four irons (c). The cross sections of the reconstituted difference map are plotted in Fig. S24. A raw difference map of the observed all-ferrous cluster [4Fe4S]^0^ without expansion is obtained by an illumination of 450 nm at 250 mA (Fig. S23). The reconstituted difference map 640***U***_1_ - 525***U***_2_ + 600***U***_3_ shows positive densities on four irons that skew away from the center (d). The cross sections of the reconstituted difference map are plotted in Fig. S26. A raw difference map of the expanded all-ferrous cluster [4Fe4S]^0^ is obtained by an illumination of 450 nm at 150 mA (Fig. S25).

Several other datasets also require the destructive combination 640***U***_1_ – 525***U***_2_ of the first two components that restores the tetrahedral symmetry and significantly weakens the layered features. These datasets likely capture the all-ferrous [4Fe4S]^0^ by illuminations of both violet and blue light (Figs. S23 and S25). These observations seem to contradict the two-layer model, which predicts that four electrons of the minor spin should be distributed in two opposite faces. All-ferrous cluster [4Fe4S]^0^ has been previously detected in proteins. The most studied one is a nitrogenase that reduces nitrogen to ammonia. The [4Fe4S]^0^ in nitrogenase could carry a total spin of 0 or 4, in which the high spin state consists of one ferrous site different from the other three (Yoo et al., 1999). Two Fe-Fe distances were found longer than the other four by X-ray absorption spectroscopy that studies the fine structure of Fe K-edge (Musgrave et al., 1998), which is beyond the resolution available to this work. Nevertheless, these observations seem to suggest that the two-layer model no longer applies to the all-ferrous cluster. Our datasets show significant variations in the third component (Fig. S12ac). Due to the larger array of 12 positive peaks, the positive densities in ***U***_3_ distribute consistently outside the [4Fe4S] cluster (Fig. 4b). Therefore, an addition of ***U***_3_ appears to expand the size of the neutral, all-ferrous cluster [4Fe4S]^0^ compared to a subtraction of this component (Figs. 4cd, S24, and S26). A difference map captured by blue light illumination at 250 mA does not show the cluster expansion (Fig. S23), while another at 150 mA shows the most expansion (Figs. S12abc and S25). We hypothesize that the neutral cluster [4Fe4S]^0^ has gained sufficient electrons so that the repulsion forces among them could start to influence the geometry of the cluster (Figs. 2ac bottom right). This is consistent with the previous result of two longer Fe-Fe distances (Musgrave et al., 1998). However, it is not clear from our maps which two Fe-Fe distances are longer due to the limited resolution from protein crystals. Or these longer Fe-Fe distances could be distributed randomly so that the all-ferrous cluster in our maps appears larger than normal.

An orthogonal technique demonstrating the existence of all-ferric and all-ferrous clusters in PhrB under illumination could be difficult. So far, we find no specific redox or illumination condition that would certainly lead to two-electron oxidation or reduction from [4Fe4S]^2+^. In addition, even if the [4Fe4S] cluster in PhrB reaches all-ferric or all-ferrous state, the lifetime could be as short as milliseconds according to the fluctuation frequency of a major spectral component derived from time-resolved absorption spectra (Ren et al., 2022). These highly oxidized or reduced clusters may not populate significantly in a steady state. Therefore, the signals of the all-ferric and all-ferrous clusters could be swamped by the signals from the other redox states. However, these short-lived clusters could be captured by Laue diffraction with exposure times in microseconds (Methods).

### Reorientation of valence layers

In addition to the observed changes in redox states that require a combination that involves the first component, we observe nearly half of the datasets under the illuminations of both violet and blue light that are largely absent of ***U***_1_ (Fig. 2a). Like the others, these datasets consist of either a positive or a negative term of ***U***_2_ and a variable amount of ***U***_3_ without a clear discreteness. Little difference densities, positive or negative, are found on the irons or sulfurs, which leads us to conclude that these light-induced difference maps depict changes within [4Fe4S]^2+^ that does not involve a net gain or loss of electrons. Alternatively, zero net gain after multiple redox steps is a more likely possibility. The fact that nearly half of the datasets fall in these groups demonstrates that the [4Fe4S] cluster under light illumination likely remain identical to the dark, 2+ valence state. We interpret these difference maps as a result that one electron is transferred from each valence layer shaded in red to another shaded in blue in Fig. 5ab, directly or indirectly, that is, a change in the orientation of the valence layers (Fig. 2c second column). The arrays of 12 peaks are the characteristic features of ***U***_2_ and ***U***_3_, the main sources of signal indicating layered electronic structure (Figs. 2b, S14, and S15). Linear combinations of these two components but without ***U***_1_ result in two 12-peak arrays coexist and transition from one to another. For example, Fig. 5c demonstrates that an array of 3×2×2 negative peaks in X, Y, and Z directions, respectively, gradually transition to the array of 2×2×3 positive peaks as the coefficient *c*_3_ varies from -300 to 600 (Figs. 5c and S12). The captured wide range of *c*_3_ coefficient indicates that the reorientation of valence layers is not quantized but could occur in various extent. The observed difference map by blue light illumination at 200 mA shows a 3×2×2 positive peaks in X, Y, and Z directions, respectively (Fig. S27), while another observed difference map by violet light illumination at 40 mA displays 2×2×3 positive peaks (Fig. S29). Reconstituted maps to fit these observations essentially show the same (Figs. S28 and S30). Two mixed valence layers shaded horizontally in Y2 cross section transition to other two mixed valence layers shaded vertically in the mid column of Fig. 2a.

**Figure 5.**
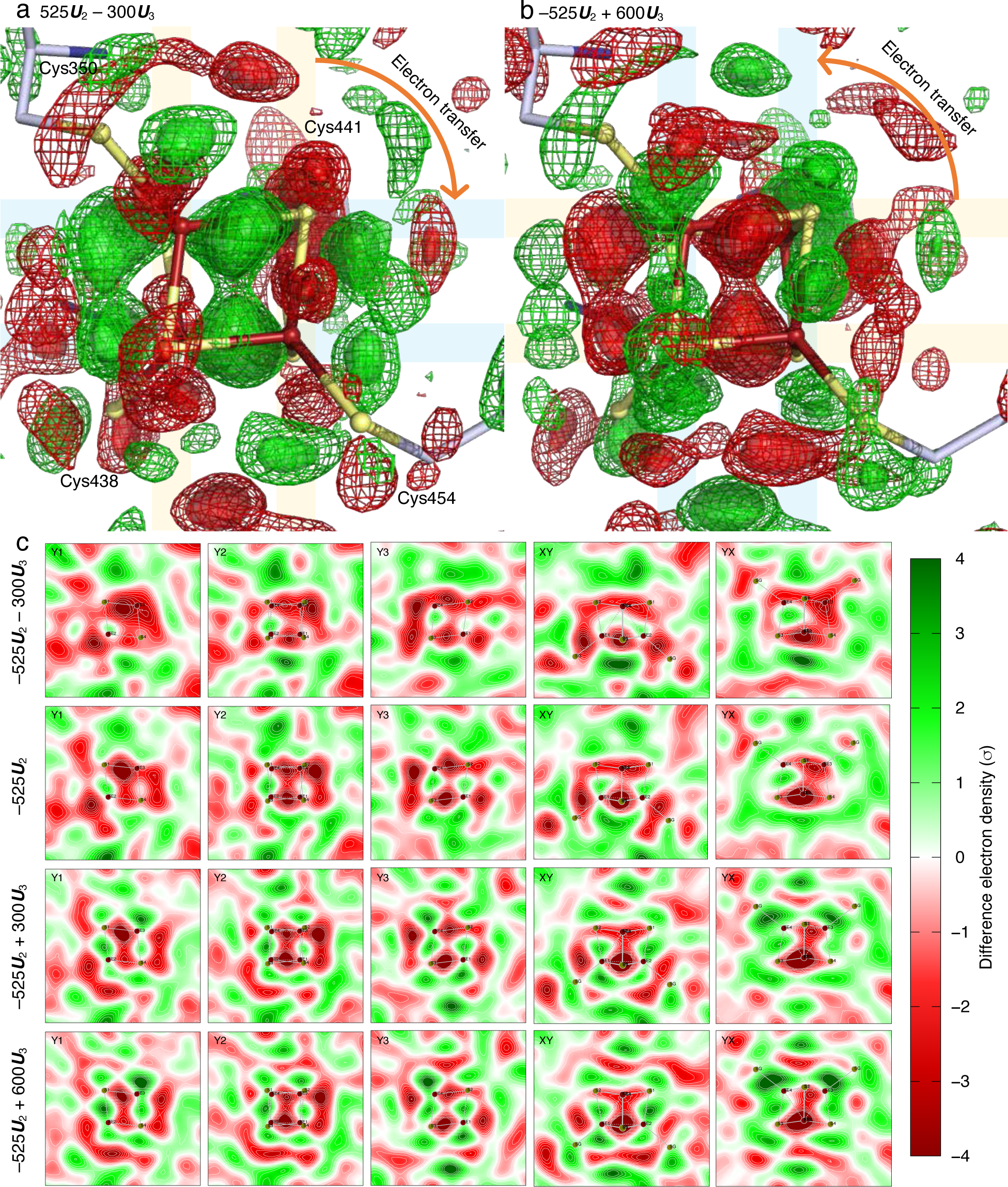
Light induced reorientation of valence layers in [4Fe4S]^2+^. (a) and (b) These reconstituted difference maps are contoured at ±2*σ* with mesh and ±3*σ* with surfaces. The positive and negative contours are in green and red, respectively. They represent two extremes of the observed difference maps that do not involve the first component (Figs. S28 and S30). 525***U***_2_ – 300***U***_3_ in (a) reproduces the observed difference map by blue-light illumination at 200 mA (Fig. S27). -525***U***_2_ + 600***U***_3_ in (b) reproduces that by violet-light illumination at 40 mA (Fig. S29) and 60 mA. An electron in a valence layer shaded in red is transferred directly or indirectly to another layer shaded in blue. Thus, the red layers are oxidized, and the blue layers are reduced without a net valence change. (c) Linear combinations with variable amount of ***U***_3_. An array of 3×2×2 negative peaks (top row) in the directions X, Y, and Z axes, respectively, is transitioning to an array of 2×2×3 positive peaks (bottom row) as more and more ***U***_3_ is added into the linear combination.

These observations imply an equilibrium between valence layers normal to multiple axes in the dark resting state. No unique orientation of the valence layers stands out in dark. Therefore, our observed light – dark difference maps are with respect to a reference of a good tetrahedral symmetry despite the inherent asymmetry of the [4Fe4S]^2+^ cluster in dark. Upon light illumination, electrons are observed exchanging between the valence layers normal to X axis (Fig. 2a bottom mid panel) and those normal to Z axis (Fig. 2a top mid panel) as a net result therefore shifting the dark equilibrium (Fig. 2c). The electron transfers between the valence layers are observed going either way to and from two orientations but not captured involving the third orientation (second columns in Fig. 2ac). One or even two electrons in the valence layers at one orientation could transfer first to somewhere else, such as the flavin or the ribolumazine cofactors as we presented previously (Ren et al., 2022), resulting in [4Fe4S]^3+^ or even [4Fe4S]^4+^ and then transfer back to restore the [4Fe4S]^2+^ cluster. The returned electrons may occupy valence layers at the same or a different orientation. Alternatively, the reorientation of valence layers could be the net result from gaining electrons first before losing them, that is, going through [4Fe4S]^+^ or even [4Fe4S]^0^.

### Electron density changes observed by cryo-trapping

An independent SVD analysis is conducted on a total of 26 difference maps obtained at cryo temperatures (Fig. 6). The first component clearly shows a displacement of the [4Fe4S] cluster and the surrounding helices (Figs. 6b, S10, and S31). This type of signal is not observed at room temperature, at least not among the top components from SVD. We suspect that these conformational changes are the consequence of the light induced redox reaction in PhrB because this component is not required for any map derived from the control datasets that test the effect of X-ray damage (Fig. 6d). These conformational changes do not seem to hold for long at room temperature. They are beyond the scope of the quantum effect of spin coupling discussed in this paper and will be further explored and presented elsewhere.

**Figure 6.**
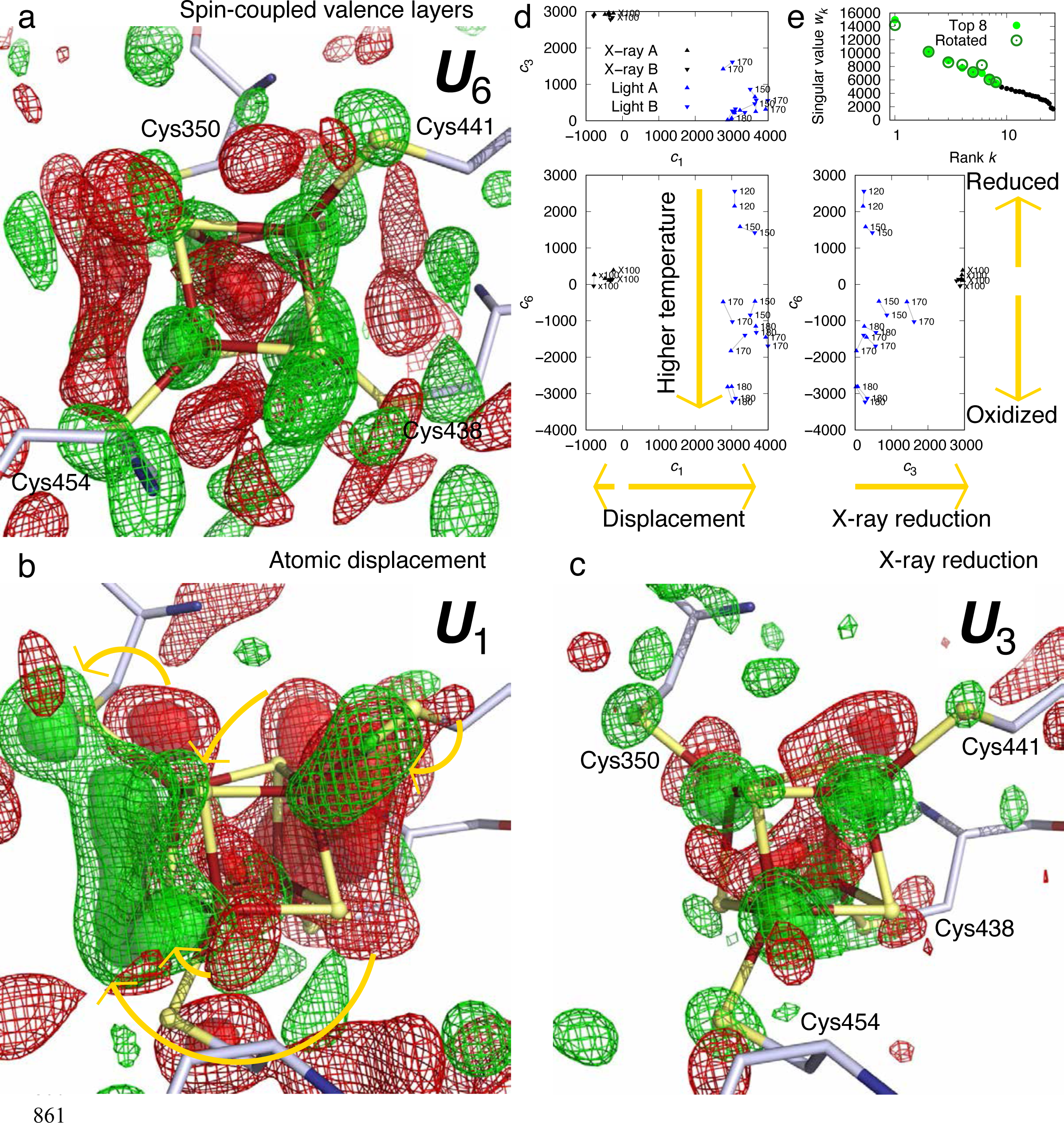
SVD analysis of difference Fourier maps in cryo-trapping experiments. (a) Sixth component ***U***_6_ of difference maps at cryo temperatures. This component map is contoured at ±2.5*σ* with mesh and ±5*σ* with surfaces. The positive and negative contours are in green and red, respectively. This component is equivalent to the first component ***U***_1_ of the difference maps at room temperature (Figs. 2b and S13). See Fig. S33 for detailed plots of the cross sections through this component. (b) First component ***U***_1_ of difference maps at cryo temperatures. The positive and negative contours are in green and red, respectively, at ±3*σ* with mesh and ±5*σ* with surfaces. Arrows mark the paired negative and positive densities indicating the observed atomic motions in the first component ***U***_1_. See Figs. S31 for detailed plots of the cross sections through this component. (c) Third component ***U***_3_ of difference maps at cryo temperatures. Contoured the same way as (b). See Figs. S32 for detailed plots of the cross sections through this component. The other major components describe the differences between two protein chains related by noncrystallographic symmetry. (d) Correlation between coefficients for the first, third, and sixth components. Difference maps from two protein chains related by the noncrystallography symmetry are represented by up- and down-pointing triangles, respectively, and connected by a gray line. The illumination temperature in K is marked wherever possible. A positive value of *c*_6_ indicates a reduction of [4Fe4S]^2+^, while a negative value means an oxidation. Difference maps derived from the control experiments without a light illumination but with variable X-ray doses are in black. These maps carry a small negative term of ***U***_1_ but do not feature ***U***_6_. Therefore, ***U***_6_ depicts the light-induced redox responses, and ***U***_3_ displays the X-ray induced reduction to the [4Fe4S] cluster. (e) Singular values derived from SVD before and after Ren rotation (Methods).

The third component ***U***_3_ is mostly featured in the difference maps of the X-ray dose effect because only a map derived from high X-ray dose – low dose requires a significant coefficient of *c*_3_. Therefore, this component depicts the image of X-ray reduction (Fig. 6d). Four positive peaks of 7-9*σ* are found on the irons. Weaker positive peaks up to 5*σ* are on the Sγ atoms of the Cys residues. Only slight positive peaks up to 3*σ* are near the sulfurs in [4Fe4S] cluster (Figs. 6c and S32). This component largely follows the tetrahedral symmetry. It seems that X-ray reduction does not produce a layered electronic structure in a specific direction. However, electron relaxation of passive orbitals still applies in X-ray induced reduction of iron-sulfur clusters. The light - dark maps carry small amounts of ***U***_3_ indicating that our strategy to acquire dark and light datasets from different segments of the same crystal keeps the radiation damage in check (Fig. 6d).

The sixth component ***U***_6_ of the cryo-trapping maps is qualitatively as same as ***U***_1_ from the serial Laue diffraction at room temperature, and again a good experimental observation of electron relaxation and layered electronic structure due to spin coupling (compare Figs. S13 and S33). The positive peaks on the irons range from 6 to 9*σ*. A central negative peak at -7*σ* is similar to that of ***U***_1_ of the difference maps at room temperature. The positive peaks 2.5-5*σ*associated with the eight sulfur atoms are skewed outward away from the center (Fig. S33). What the sixth component of the cryo data differs from ***U***_1_ of the serial Laue data is the orientation of the layers normal to Y axis (Figs. 6a and S33). This again shows that the valence layers of the irons could change orientation. None of the difference maps of the X-ray reduction consists of this component as their coefficients *c*_6_ ≍ 0 (Fig. 6d). That is to say, the X-ray damage datasets are good controls to show that the change of valence layers are light induced, not X-ray induced. In contrast, the coefficient of the sixth component could be negative or positive for the light – dark maps (Fig. 6d), which is again consistent with our spectroscopic data and serial Laue data, that is, the [4Fe4S] cluster functions as a source and a storage of electrons as needed depending on the redox balance of FAD and DMRL (Ren et al., 2022). There is a tendency that a difference map derived at a more elevated cryo temperature would require a more negative component of ***U***_6_. That is, the [4Fe4S] cluster is more likely to be oxidized at higher temperature. However, the cryo-trapping data present no further details compared to the serial Laue data at the room temperature.

## Discussion

It has been widely believed that a crystallographic observation of electron density distribution of a single or several individual electrons would require ultrahigh spatial resolution such as those demonstrated in charge density analyses of small molecule crystallography. An individual event of electron transfer in protein, the essence of biochemistry, is usually out of reach largely due to the insufficient spatial resolution achievable by protein crystallography, and in case of transient events, the insufficient temporal resolution as well. Therefore, the goal of quantum crystallography, defined by Massa et al. as for extracting molecular quantum properties from X-ray diffraction (Massa et al., 1995), is seemingly infeasible for biological macromolecules despite the great desire (Polkosnik et al., 2019). However, we argued recently that the difficulty to visualize electron density changes from protein crystals due to a quantized electronic event originates from the inability to experimentally isolate such event rather than the inability to capture sharp images at atomic and subatomic resolution (Ren, 2019). Our reasoning was based on the fact that an experimental electron density map frequently captures multiple coexisting changes. For example, we obtained one of the SVD components of difference Fourier maps derived from the ultrafast photolysis of carbonmonoxy myoglobin that resembles a superposition of several 3*d* orbitals of the heme Fe(II). We interpretated the positive densities of this component as the net gain of electron distribution while the heme Fe(II) returns to its high-spin state by occupying all of its 3*d* orbitals while its sixth ligand is lost by photolysis. The crossover of the heme Fe(II) from low- to high-spin takes place concurrently with the other repeatedly observed events such as the departure of the CO ligand, the heme buckling, and the out-of-plane displacement of the iron. The electron density redistribution due to the spin crossover is relatively minor compared to the other relocation of many tens or even hundreds of electrons, therefore, completely immersed and not easily observable (Ren, 2019). This work offers another example to show that electron density redistribution due to relocations of a few electrons could stand out as the top three map components in absence of atomic displacement (Figs. 1 and 2). However, when atomic displacements are present, the most significant map component describes these displacements that easily carry many tens of electrons (Figs. 6b, S10, and S31). The remaining signals due to light-induced redox responses of the iron-sulfur cluster are captured by a relatively minor, sixth component (Figs. 6a and S33). Even X-ray reduction signals are ranked higher as the third component (Figs. 6c and S32).

Therefore, an individual electronic event in protein is usually overwhelmed by other changes. This cause for difficulty in detection of individual electronic events fundamentally differs from the previous reasoning – the lack of spatial resolution. We argue that ultrahigh spatial resolution is not required to analytically depict an electron redistribution caused by an electron transfer event because none of the probability distribution functions of electrons features a mathematical discontinuity that could only be properly depicted by high frequency terms in the Fourier summation, that is, ultrahigh spatial resolution.

The decomposition of difference Fourier maps and the subsequent deconvolution to achieve a physical or chemical interpretation by Ren rotation (Ren, 2022, 2021, 2019, 2016) are therefore crucial to a better understanding of several concurrent events. Here in this work, again we dissect many observed difference Fourier maps into multiple components. Each component plays a role in the interpretation of a total of seven redox states of the [4Fe4S] cluster (Fig. 2c). This work demonstrates that a fundamental quantum mechanical property, here the spin coupling among the clustered metal centers incorporated in proteins, could be extracted and evaluated from electron density distributions, the source of X-ray diffraction. The methods employed and extended here could open new opportunities to visualize essential processes in biochemistry at the level of individual electronic events.

## Acknowledgements

Use of Advanced Photon Source, an Office of Science User Facility operated for US Department of Energy by Argonne National Laboratory, was supported by contract DE-AC02-06CH11357. Use of BioCARS was supported by the National Institute of General Medical Sciences of the National Institutes of Health under grant number R24GM111072. The content is solely the responsibility of the authors and does not necessarily represent the official views of the National Institutes of Health. Use of LS-CAT Sector 21 was supported by Michigan Economic Development Corporation and Michigan Technology Tri-Corridor grant 085P1000817. This work is supported by grants from the University of Illinois at Chicago, National Institutes of Health (R01EY024363), and National Science Foundation (MCB 2017274) to XY. This work is also supported by grants from Deutsche Forschungsgemeinschaft to TL (LA 799/7-3) and NK (KR 2034/1). The China Scholarship Council and Karlsruhe House of Young Scientists are acknowledged to support the Ph.D. and postdoc studies of FZ.

## Data availability

Laue diffraction images, structure factor sets, and difference Fourier maps are available from the corresponding authors upon request.

## Code availability

The following software packages are used in this work: Coot (Emsley and Cowtan, 2004; www2.mrc-lmb.cam.ac.uk/Personal/pemsley/coot), dynamiX™ (Renz Research, Inc.), HKL2000 (Otwinowski and Minor, 1997; hkl-xray.com), Precognition/Epinorm™ (Renz Research, Inc.), PyMOL (pymol.org), Python (python.org), SciPy (scipy.org), and serialX (ZR).

## Methods

### Expression, purification, and crystallization

Wildtype PhrB protein is expressed and purified as previously described (Oberpichler et al., 2011). Crystallization of wildtype PhrB using the method of hanging drop vapor diffusion follows the published protocol (Zhang et al., 2013). These crystals are used for spectroscopic measurements (Fig. S2). PhrB crystallization on chip uses the microbatch method (Biju et al., 2022; Ren et al., 2018). The crystallization chamber is about 150 μm in thickness. On each device, 16 μL of crystallization solution contains 3.5-4.5 mg/mL protein, 3.4% PEG 400, and 33 mM MES buffer [2-(N-morpholino)ethanesulfonic acid] at pH 5.6. In addition, seeding of the crystallization solution stimulates the growth of more and chunkier crystals. Seeds are produced by grinding of previous crystals. New PhrB crystals of typical size of 300 × 30 × 30 μm^3^ appear on chips within a few days (Fig. S5). Altogether, 69 devices consume < 5 mg of protein (Table S1). All devices are stored in dark, and some devices are monitored for crystal growth with a near infrared (IR) light source using a digital camera.

### Absorbance measurement of PhrB single crystals

Absorption spectra of PhrB crystals are measured by an Ocean Optics spectrometer QE Pro with a deuterium-halogen light source DH-2000-BAL (Ocean Insight). This combination of instruments covers a spectral range from mid-UV to near IR (200-1000 nm). PhrB crystals are pumped by violet and blue light illuminations at 405 and 450 nm, respectively. Two laser diodes L405P150 and L450P1600MM (Thorlabs, Inc.) are collimated by an aspherical lens. Power densities of the pump light at various current settings (Fig. S4) are measured by an optical power meter PM100USB (Thorlabs, Inc.). The timing control of these light sources and the spectrometer (Fig. S3a) is programmed on a Raspberry Pi microcomputer (Raspberry Pi Foundation) by taking advantages of its versatile general-purpose input-output (GPIO).

### Serial Laue diffraction at room temperature

Laue diffraction experiments (Ren et al., 1999) are conducted in December 2018, June and November 2019 at the BioCARS beamline 14-ID-B of Advanced Photon Source (APS). PhrB crystals grown on chip are imaged with near IR light and recognized automatically by an image analysis software (Fig. S5). Qualified crystals are preprogrammed in a sequence (Fig. S6). The exposure time of each crystal or each segment of a crystal is 5-8 μs, which consists of 33-50 consecutive single bunches of the synchrotron evenly spaced out in time. Each single bunch lasts about 100 ps (Graber et al., 2011). X-ray exposures in the light datasets occur after a two-second illumination of violet or blue light at 405 and 450 nm, respectively. The continuous wave laser light is focused and trimmed to a spot of ∼300 μm in diameter. A wide range of power density is controlled by setting the electric current through the laser diodes (Fig. S4). Diffraction images are recorded by a Rayonix MX 340 X-ray detector (Rayonix, LLC). Experimental details using our newly developed inSituX serial crystallography platform were published recently (Biju et al., 2022; Ren et al., 2018, 2020). Laue diffraction images are processed with the software package Precognition/Epinorm™ (Ren, 2006, 2008) with an additional software development called serialX to suit applications in serial crystallography (Table S1 and Figs. S7-S9).

### Temperature scan cryo-trapping

The protocol of temperature scan cryo-trapping is schematically presented in Fig. S3b. A dark dataset is collected from one segment of each crystal before any illumination. One or more light datasets are collected from other fresh segments of the same crystal after illuminations at elevated cryo temperatures (Table S2). We also conduct control experiments to evaluate the effect of X-ray radiation damage. Multiple datasets are collected from the same crystal segment in dark at 100 K. The earlier datasets represent a low dose data point, while the later datasets represent a high dose one. Difference maps of high X-ray dose – low dose reveal the effect the X-ray alone (Table S2). The monochromatic X-ray diffraction datasets at 100 K are collected at the LS-CAT beamline 21-ID-D of APS. All diffraction images are collected on an Eiger 9M detector (Dectris Ltd.) and processed using HKL2000 (Otwinowski and Minor, 1997).

### Difference Fourier maps

A difference Fourier map is synthesized from a Fourier coefficient set of *F*_light_-*F*_reference_ with the best available phase set, often from the ground state structure. Before Fourier synthesis, *F*_light_ and *F*_reference_ must be properly scaled to the same level so that the distribution of difference values is centered at zero and not skewed either way. A weighting scheme proven effective assumes that a greater amplitude of a difference Fourier coefficient *F*_light_-*F*_reference_ is more likely caused by noise than by signal (Ren et al., 2001, 2013; Šrajer et al., 2001; Ursby and Bourgeois, 1997). Both the dark and light datasets can serve as a reference in difference maps. However, both the dark and light datasets must be collected in the same experiment. A cross reference from a different experimental setting usually causes large systematic errors in the difference map that would swamp the desired signals. Each difference map is masked 4 Å around the [4Fe4S] cluster and the four anchor Cys residues.

### Singular value decomposition of (difference) electron density maps

An electron density map, particularly a difference map as emphasized here, consists of density values on an array of grid points within a mask of interest. All *M* grid points in a three-dimensional map can be serialized into a one-dimensional sequence of density values according to a specific protocol. It is not important what the protocol is as long as a consistent protocol is used to serialize all maps of the same grid setting and size, and a reverse protocol is available to erect a three-dimensional map from a sequence of *M* densities. Therefore, a set of *N* serialized maps, also known as vectors in linear algebra, can fill the columns of a data matrix **A** with no specific order, so that the width of **A** is *N* columns, and the length is *M* rows. Often, *M* >> *N*, thus **A** is an elongated matrix. If a consistent protocol of serialization is used, the corresponding voxel in all *N* maps occupies a single row of matrix **A**. This strict correspondence in a row of matrix **A** is important. Changes of the density values in a row from one structure to another are due to either signals, systematic errors, or noises. Although the order of columns in matrix **A** is unimportant, needless to say, the metadata associated with each column must remain in good bookkeeping.

SVD of the data matrix **A** results **A = UWV**^T^, also known as matrix factorization. Matrix **U** has the same shape as **A**, that is, *M* rows and *N* columns. The *N* columns contain decomposed basis components ***U****_k_*, known as left singular vectors of *M* items each, where *k* = 1, 2, …, *N*. Therefore, each component ***U****_k_* can be erected using the reverse protocol to form a three-dimensional map. This decomposed elemental map can be presented in the same way as the original maps, for example, rendered in molecular graphics software such as Coot and PyMol. It is worth noting that these decomposed elemental maps or map components ***U****_k_* are independent of any metadata. That is to say, these components remain constant when the metadata vary. Since each left singular vector ***U****_k_* has a unit length due to the orthonormal property of SVD (see below), that is, |***U****_k_*| = 1, the root mean squares (rms) of the items in a left singular vector is 1/√*M* that measures the quadratic mean of the items.

The second matrix **W** is a square matrix that contains all zeros except for *N* positive values on its major diagonal, known as singular values *w_k_*. The magnitude of *w_k_* is considered as a weight or significance of its corresponding component ***U****_k_*. The third matrix **V** is also a square matrix of *N* × *N*. Each column of **V** or row of its transpose %^!^, known as a right singular vector ***V****_k_*, contains the relative compositions of ***U****_k_* in each of the *N* original maps. Therefore, each right singular vector ***V****_k_* can be considered as a function of the metadata. Right singular vectors also have the same unit length, that is, |***V****_k_*| = 1. Effectively, SVD separates the constant components independent of the metadata from the compositions that depend on the metadata.

A singular triplet denotes 1) a decomposed component ***U****_k_*, 2) its singular value *w_k_*, and 3) the composition function ***V****_k_*. Singular triplets are often sorted in a descending order of their singular values *w_k_*. Only a small number of *n* significant singular triplets identified by the greatest singular values *w*_1_ through *w_n_* can be used in a linear combination to reconstitute a set of composite maps that closely resemble the original ones in matrix **A**, where *n* < *N*. For example, the original map in the *i*th column of matrix **A** under a certain experimental condition can be closely represented by the *i*th composite map *w*_1_*v*_1*i*_***U***_1_ + *w*_2_*v*_2*i*_***U***_2_ + … + *w_n_v_ni_**U**_n_*, where (*v*_1*i*_, *v*_2*i*_, …) is from the *i*th row of matrix **V**. The coefficient set for the linear combination is redefined here as *c_ki_* = *w_k_v_ki_*/√*M*. The rms of the values in a map component, or the average magnitude measured by the quadratic mean, acts as a constant scale factor that resets the modified coefficients *c_ki_* back to the original scale of the core data, such as e^-^/Å^3^ for electron density maps if these units are used in the original matrix **A**. Practically, an electron density value usually carries an arbitrary unit without a calibration, which makes this scale factor unnecessary. In the linear combination *c*_1*i*_***U***_1_ + *c*_2*i*_***U***_2_ + … + *c_ni_**U**_n_*, each component ***U****_k_* is independent of the metadata while how much of each component is required for the approximation, that is, *c_ki_*, depends on the metadata.

Excluding the components after ***U****_n_* in this approximation is based on an assumption that the singular values after *w_n_* are very small relative to those from *w*_1_ through *w_n_*. As a result, the structural information evenly distributed in all *N* original maps is effectively concentrated into a far fewer number of *n* significant components, known as information concentration or dimension reduction. On the other hand, the trailing components in matrix **U** contain inconsistent fluctuations and random noises. Excluding these components effectively rejects noises (Schmidt et al., 2003). The least-squares property of SVD guarantees that the rejected trailing components sums up to the least squares of the discrepancies between the original core data and the approximation using the accepted components.

However, no clear boundary is guaranteed between signals, systematic errors, and noises. Systematic errors could be more significant than the desired signals. Therefore, excluding some components from 1 through *n* is also possible. An example of such systematic differences is the discrepancy between two chains related by noncrystallographic symmetry in the crystal form of PhrB. If systematic errors are correctly identified, the reconstituted map without these significant components would no longer carry the systematic errors.

### The orthonormal property of SVD

The solution set of SVD must guarantee that the columns in **U** and **V**, the left and right singular vectors ***U****_k_* and ***V****_k_*, are orthonormal, that is, ***U****_h_*•***U****_k_* = ***V****_h_*•***V****_k_* = 0 (ortho) and ***U****_k_*•***U****_k_*= ***V****_k_*•***V****_k_* = 1 (normal), where *h* ≠ *k* but both are from 1 to *N*. The orthonormal property also holds for the row vectors. As a result, each component ***U****_k_* is independent of the other components. In other words, a component cannot be represented by a linear combination of any other components. However, two physical or chemical parameters in the metadata, such as temperature and reducing agent, may cause different changes to a structure. These changes are not necessarily orthogonal. They could exhibit some correlation. Therefore, the decomposed components ***U****_k_* not necessarily represent any physically or chemically meaningful changes (see below).

Due to the orthonormal property of SVD, an *N*-dimensional Euclidean space is established, and the first *n* dimensions define its most significant subspace. Each coefficient set ***c****_i_* = (*c*_1*i*_, *c*_2*i*_, …, *c_ni_*) of the *i*th composite map is located in this *n*-dimensional subspace. All coefficient sets for *i* = 1, 2, …, *N* in different linear combinations to approximate the *N* original maps in a least-squares sense can be represented by *N* points or vectors ***c***_1_, ***c***_2_, …, ***c****_N_* in the Euclidean subspace. This *n*-dimensional subspace is essentially the conformational space as surveyed by the jointly analyzed core data. The conformational space is presented as scatter plots with each captured structure represented as a dot located at a position determined by the coefficient set ***c****_i_* of the *i*th observed map. When the subspace has greater dimensionality than two, multiple two-dimensional orthographical projections of the subspace are presented. These scatter plots are highly informative to reveal the relationship between the (difference) electron density maps and their metadata. If two coefficient sets ***c****_i_* ≍ ***c****_j_*, they are located close to each other in the conformational space. Therefore, these two structures *i* and *j* share two similar conformations, for example, the six clusters in Fig. 2a. Two structures located far apart from each other in the conformational space are distinct in their conformations, and distinct in the compositions of the map components.

### Rotation in SVD space

Dimension reduction is indeed effective in meta-analysis of protein structures when many datasets are evaluated at the same time. However, the default solution set of SVD carries complicated physical and chemical meanings that are not immediately obvious. The interpretation of a basis component ***U****_k_*, that is, “what-does-it-mean”, requires a clear demonstration of the relationship between the core data and their metadata. The outcome of SVD does not guarantee any physical meaning in a basis component. Therefore, SVD alone provides no direct answer to “what-does-it-mean”, thus its usefulness is very limited to merely a mathematical construction and some noise reduction. However, the factorized set of matrices **U**, **W**, and **V** from SVD is not a unique solution. That is to say, they are not the only solution to factorize matrix **A**. Therefore, it is very important to find one or more alternative solution sets that are physically meaningful to elucidate a structural interpretation. The concept of a rotation after SVD was introduced by Henry & Hofrichter (Henry and Hofrichter, 1992). But they suggested a protocol that fails to preserve the orthonormal and least-squares properties of SVD. The rotation protocol suggested by Ren incorporates the metadata into the analysis and combines with SVD of the core data. This rotation achieves a numerical deconvolution of multiple physical and chemical factors after a pure mathematical decomposition, and therefore, provides a route to answer the question of “what-does-it-mean” (Ren, 2019). This rotation shall not be confused with a rotation in the three-dimensional real space, in which a molecular structure resides.

A rotation in the *n*-dimensional Euclidean subspace is necessary to change the perspective before a clear relationship emerges to elucidate scientific findings. It is shown below that two linear combinations are identical before and after a rotation applied to both the basis components and their coefficients in a two-dimensional subspace of *h* and *k*. That is,

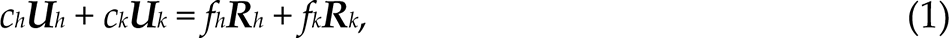

where *c_h_* and *c_k_* are the coefficients of the basis components ***U****_h_* and ***U****_k_* before the rotation; and *f_h_* and *f_k_* are the coefficients of the rotated basis components ***R****_h_* and ***R****_k_*, respectively. The same Givens rotation of an angle *θ* is applied to both the components and their coefficients:

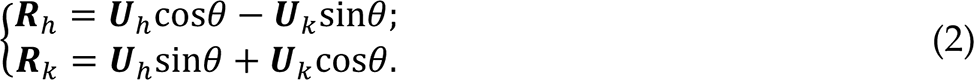

Obviously, the rotated components ***R****_h_* and ***R****_k_* remain mutually orthonormal and orthonormal to other components. And

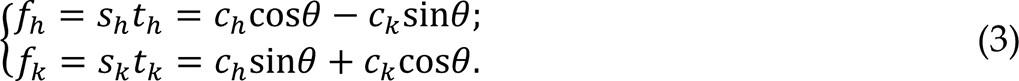

Here 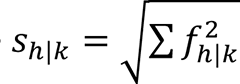 are the singular values that replace *w_h_* and *w_k_*, respectively, after the rotation. They may increase or decrease compared to the original singular values so that the descending order of the singular values no longer holds. ***T****_h_*_|*k*_ = (*t_h_*_|*k*1_, *t_h_*_|*k*2_, …, *t_h_*_|*kN*_) = (*f_h_*_|*k*1_, *f_h_*_|*k*2_, …, *f_h_*_|*kN*_)/*s_h_*_|*k*_ are the right singular vectors that replace ***V****_h_* and ***V****_k_*, respectively. ***T****_h_* and ***T****_k_* remain mutually orthonormal after the rotation and orthonormal to other right singular vectors that are not involved in the rotation.

Eq. 1 holds because the dot product of two vectors does not change after both vectors rotate the same angle. To prove Eq. 1 in more detail, Eqs. 2 and 3 are combined and expanded. All cross terms of sine and cosine are self-canceled:

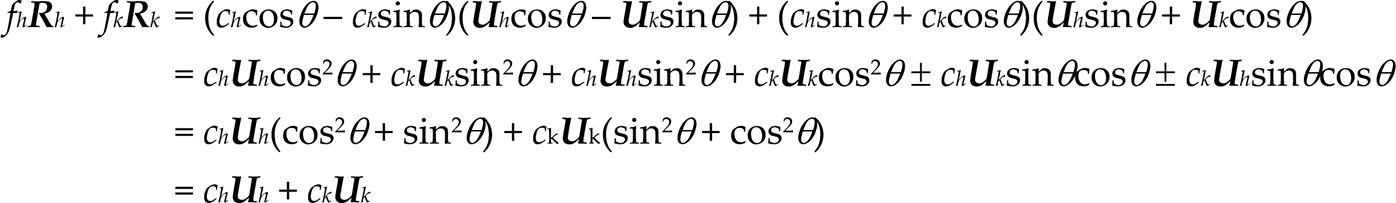

where ± signs indicate the presence of both the positive and negative terms simultaneously, therefore canceling each other.

A rotation in two-dimensional subspace of *h* and *k* has no effect in other dimensions, as the orthonormal property of SVD guarantees. Multiple steps of rotations can be carried out in many two-dimensional subspaces consecutively to achieve a multi-dimensional rotation. A new solution set derived from a rotation retains the orthonormal property of SVD. The rotation in the Euclidean subspace established by SVD does not change the comparison among the core data of protein structures. Rather it converts one solution set **A** = **UWV**^T^ to other alternative solutions **A** = **RST**^T^ so that an appropriate perspective can be found to elucidate the relationship between the core data and metadata clearly and concisely.

For example, if one physical parameter could be reoriented along a single dimension *k* but not involving other dimensions by a rotation, it would be convincing to show that the left singular vector ***U****_k_* of this dimension illustrates the structural impact by this physical parameter. Before this rotation, the same physical parameter may appear to cause structural variations along several dimensions, which leads to a difficult interpretation. Would a proper rotation establish a one-on-one correspondence from all physical or chemical parameters to all the dimensions? It depends on whether each parameter induces an orthogonal structural change, that is, whether structural responses to different parameters are independent or correlated among one another. If structural changes are indeed orthogonal, it should be possible to find a proper rotation to cleanly separate them in different dimensions. Otherwise, two different rotations are necessary to isolate two correlated responses, but one at a time.

For another example, if the observed core datasets form two clusters in the conformational space, a rotation would be desirable to separate these clusters along a single dimension *k* but to align these clusters along other dimensions. Therefore, the component ***U****_k_* is clearly due to the structural transition from one cluster to the other. Without a proper rotation, the difference between these clusters could be complicated with multiple dimensions involved. A deterministic solution depends on whether a clear correlation exists between the core data and metadata. A proper rotation may require a user decision. A wrong choice of rotation may select a viewpoint that hinders a concise conclusion. However, it would not alter the shape of the reaction trajectory, nor create or eliminate an intrinsic structural feature. A wrong choice of rotation cannot eliminate the fact that a large gap exists between two clusters of observed core datasets except that these clusters are not obvious from that viewpoint. A different rotation may reorient the perspective along another direction. But the structural conclusion would be equivalent. See example of before and after a rotation in (Ren, 2016).

This rotation procedure finally connects the core crystallographic datasets to the metadata of experimental conditions and accomplishes the deconvolution of physical or chemical factors that are not always orthogonal to one another after a mathematical decomposition. SVD analysis presented in this paper employs rotations extensively except that no distinction is made in the symbols of components and coefficients before and after a rotation except in this section. This method is widely applicable in large-scale structural comparisons. Furthermore, Ren rotation after SVD is not limited to crystallography and may impact other fields wherever SVD is used. For example, SVD is frequently applied to spectroscopic data, images, and genetic sequence data.

## Supplementary Tables

**Table S1.** Laue data statistics

| Illumination<br>(nm/mA/s) | Dev. | Images merged/<br>collected = index<br>rate (%) | Spots/unique reflections =<br>redundancy | Harmonic<br>/spatial<br>overlaps | <sup>1</sup>Resolution/<br><sup>2</sup>completeness<br>(Å/%) | <sup>2</sup><F>/$\sigma$(F) | R<sub>merge</sub>/<br>R<sub>1/2</sub> (%) | CC<sub>1/2</sub> |
| --- | --- | --- | --- | --- | --- | --- | --- | --- |
| Dark 1 | 9 | 370/1,490 = 24.8 | 992,217/82,932 = 12.0 | 14,278/813 | 2.0 (2.1)/94.0 (69.1) | 9.2 (7.0) | 28.7/23.0 | 0.7472 |
| 450/100/2 | 9 | 844/1,835 = 46.0 | 2,402,722/83,995 = 28.6 | 14,637/1,287 | 2.0 (2.1)/98.4 (89.6) | 13.1 (7.0) | 28.0/13.8 | 0.9551 |
| 450/200/2 | 9 | 481/1,490 = 32.3 | 1,343,900/83,577 = 16.1 | 14,291/880 | 2.0 (2.1)/95.5 (75.5) | 10.4 (6.8) | 27.6/18.5 | 0.8866 |
| Dark 2 | 1 | 249/584 = 42.6 | 987,953/109,059 = 9.1 | 18,344/1,133 | 1.8 (1.9)/92.7 (65.3) | 9.2 (7.1) | 25.8/23.4 | 0.8432 |
| 405/50/2 | 5 | 1,288/2,425 = 53.1 | 5,211,578/115,795 = 45.0 | 19,007/1,918 | 1.8 (1.9)/97.8 (82.6) | 18.9 (10.9) | 25.9/13.7 | 0.9199 |
| 405/150/2 | 6 | 717/1,427 = 50.2 | 2,861,018/115,143 = 24.8 | 19,635/1,826 | 1.8 (1.9)/96.8 (77.5) | 14.2 (9.2) | 26.1/15.0 | 0.9278 |
| Dark 3 | 4 | 710/1,670 = 42.5 | 3,261,477/133,504 = 24.4 | 24,468/1,896 | 1.7 (1.8)/88.1 (43.4) | 12.7 (7.2) | 29.3/13.8 | 0.9473 |
| 405/40/2 | 2 | 940/1,851 = 50.8 | 4,308,260/134,003 = 32.2 | 23,274/1,723 | 1.7 (1.8)/91.5 (55.9) | 14.6 (7.9) | 28.2/12.7 | 0.9504 |
| 405/60/2 | 2 | 918/1,730 = 53.1 | 4,201,704/134,431 = 31.3 | 24,602/2,030 | 1.7 (1.8)/91.2 (55.5) | 14.8 (7.7) | 28.1/14.9 | 0.9273 |
| 405/70/2 | 1 | 582/1,235 = 47.1 | 2,578,183/131,302 = 19.6 | 22,280/1,335 | 1.7 (1.8)/87.6 (43.6) | 13.1 (9.0) | 27.3/15.8 | 0.8622 |
| 405/80/2 | 2 | 755/1,390 = 54.3 | 3,524,649/133,458 = 26.4 | 24,181/1,945 | 1.7 (1.8)/88.1 (44.7) | 12.5 (6.7) | 29.0/15.6 | 0.9450 |
| 405/100/2 | 2 | 697/1,686 = 41.3 | 3,187,178/132,978 = 24.0 | 23,541/1,786 | 1.7 (1.8)/88.2 (46.0) | 12.3 (7.4) | 30.0/26.9 | 0.7098 |
| 405/120/2 | 2 | 791/1,619 = 48.9 | 3,662,195/132,806 = 27.6 | 23,360/1,621 | 1.7 (1.8)/88.9 (48.1) | 13.3 (7.6) | 29.1/19.4 | 0.9066 |
| 405/170/2 | 3 | 1,114/1,997 = 55.8 | 5,154,250/134,651 = 38.3 | 21,333/1,379 | 1.7 (1.8)/92.3 (57.2) | 15.1 (7.9) | 29.5/12.0 | 0.9519 |
| 450/50/2 | 3 | 653/1,509 = 43.3 | 3,041,423/132,443 = 23.0 | 26,118/2,258 | 1.7 (1.8)/85.5 (39.9) | 11.8 (6.4) | 30.0/24.9 | 0.7708 |
| 450/70/2 | 3 | 837/1,347 = 62.1 | 3,967,265/133,636 = 36.1 | 22,805/1,677 | 1.7 (1.8)/90.5 (52.1) | 13.3 (6.7) | 28.4/14.6 | 0.9527 |
| 450/150/2 | 2 | 961/1,732 = 55.5 | 4,306,422/134,371 = 32.0 | 23,427/1,761 | 1.7 (1.8)/92.1 (60.5) | 15.7 (9.0) | 27.7/13.4 | 0.8825 |
| 450/250/2 | 1 | 361/720 = 50.1 | 1,613,038/126,429 = 12.8 | 20,784/988 | 1.7 (1.8)/79.2 (27.8) | 10.1 (7.6) | 28.4/19.7 | 0.8556 |
| 450/300/2 | 3 | 734/1,559 = 47.1 | 3,262,099/133,779 = 24.2 | 25,942/2,120 | 1.7 (1.8)/88.8 (47.0) | 13.5 (7.9) | 28.6/19.9 | 0.7883 |
| Sum/average | 69 | 14,002/29,296 = 47.8 | 59,867,531/2,318,292 = 25.8 | 406,307/30,376 | - | 13.0 (7.7) | 28.2/17.4 | 0.8805 |

<sup>1</sup>High-resolution shell is between the values inside and outside the parentheses.<sup>2</sup>The value in the high-resolution shell is in parentheses.

**Table S2.**
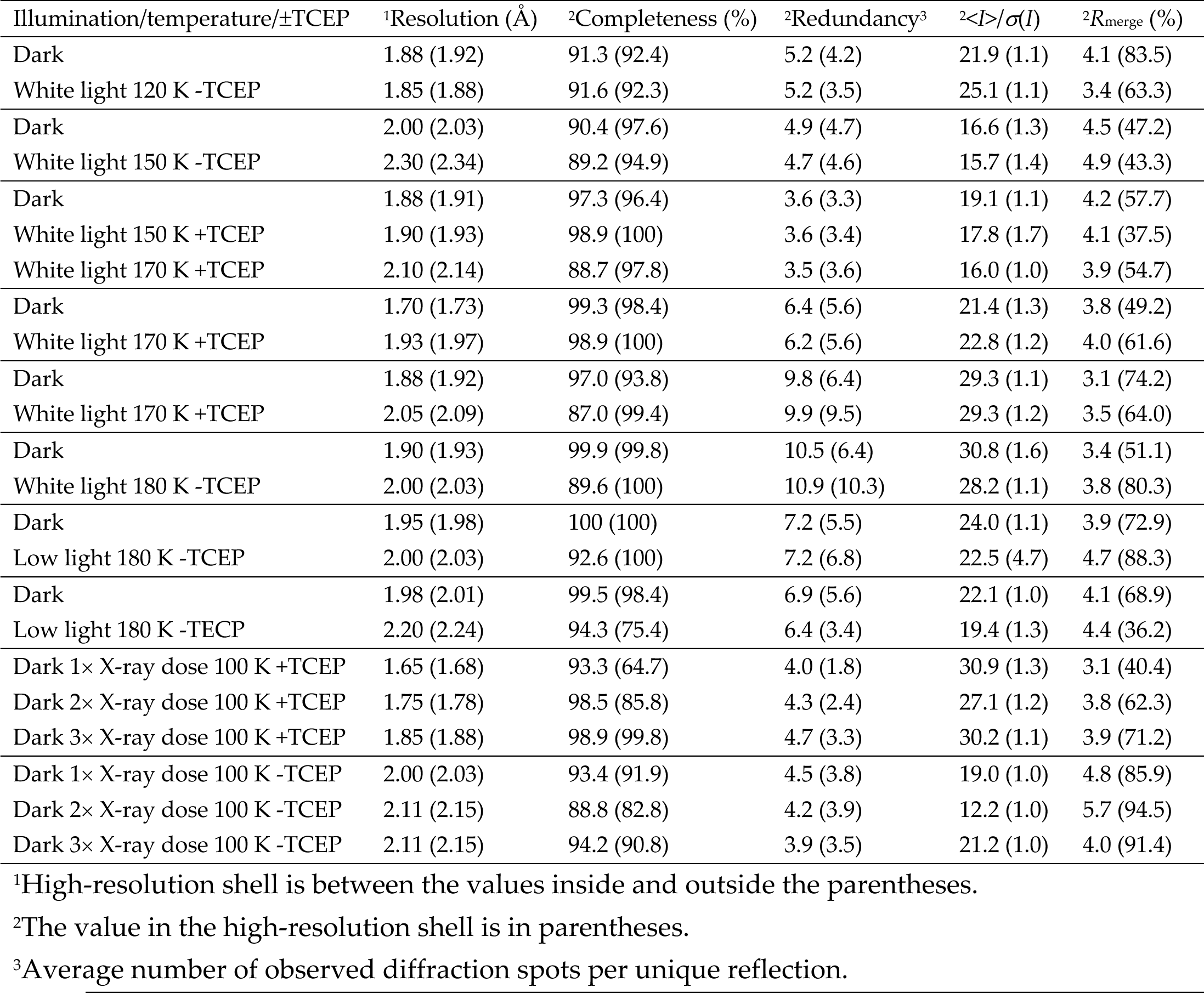
Monochromatic data statistics

| Illumination/temperature/±TCEP | <sup>1</sup>Resolution (Å) | <sup>2</sup>Completeness (%) | <sup>2</sup>Redundancy<sup>3</sup> | <sup>2</sup>$\langle I \rangle / \sigma(I)$ | <sup>2</sup>$R_{\text{merge}}$ (%) |
| --- | --- | --- | --- | --- | --- |
| Dark | 1.88 (1.92) | 91.3 (92.4) | 5.2 (4.2) | 21.9 (1.1) | 4.1 (83.5) |
| White light 120 K -TCEP | 1.85 (1.88) | 91.6 (92.3) | 5.2 (3.5) | 25.1 (1.1) | 3.4 (63.3) |
| Dark | 2.00 (2.03) | 90.4 (97.6) | 4.9 (4.7) | 16.6 (1.3) | 4.5 (47.2) |
| White light 150 K -TCEP | 2.30 (2.34) | 89.2 (94.9) | 4.7 (4.6) | 15.7 (1.4) | 4.9 (43.3) |
| Dark | 1.88 (1.91) | 97.3 (96.4) | 3.6 (3.3) | 19.1 (1.1) | 4.2 (57.7) |
| White light 150 K +TCEP | 1.90 (1.93) | 98.9 (100) | 3.6 (3.4) | 17.8 (1.7) | 4.1 (37.5) |
| White light 170 K +TCEP | 2.10 (2.14) | 88.7 (97.8) | 3.5 (3.6) | 16.0 (1.0) | 3.9 (54.7) |
| Dark | 1.70 (1.73) | 99.3 (98.4) | 6.4 (5.6) | 21.4 (1.3) | 3.8 (49.2) |
| White light 170 K +TCEP | 1.93 (1.97) | 98.9 (100) | 6.2 (5.6) | 22.8 (1.2) | 4.0 (61.6) |
| Dark | 1.88 (1.92) | 97.0 (93.8) | 9.8 (6.4) | 29.3 (1.1) | 3.1 (74.2) |
| White light 170 K +TCEP | 2.05 (2.09) | 87.0 (99.4) | 9.9 (9.5) | 29.3 (1.2) | 3.5 (64.0) |
| Dark | 1.90 (1.93) | 99.9 (99.8) | 10.5 (6.4) | 30.8 (1.6) | 3.4 (51.1) |
| White light 180 K -TCEP | 2.00 (2.03) | 89.6 (100) | 10.9 (10.3) | 28.2 (1.1) | 3.8 (80.3) |
| Dark | 1.95 (1.98) | 100 (100) | 7.2 (5.5) | 24.0 (1.1) | 3.9 (72.9) |
| Low light 180 K -TCEP | 2.00 (2.03) | 92.6 (100) | 7.2 (6.8) | 22.5 (4.7) | 4.7 (88.3) |
| Dark | 1.98 (2.01) | 99.5 (98.4) | 6.9 (5.6) | 22.1 (1.0) | 4.1 (68.9) |
| Low light 180 K -TECP | 2.20 (2.24) | 94.3 (75.4) | 6.4 (3.4) | 19.4 (1.3) | 4.4 (36.2) |
| Dark 1× X-ray dose 100 K +TCEP | 1.65 (1.68) | 93.3 (64.7) | 4.0 (1.8) | 30.9 (1.3) | 3.1 (40.4) |
| Dark 2× X-ray dose 100 K +TCEP | 1.75 (1.78) | 98.5 (85.8) | 4.3 (2.4) | 27.1 (1.2) | 3.8 (62.3) |
| Dark 3× X-ray dose 100 K +TCEP | 1.85 (1.88) | 98.9 (99.8) | 4.7 (3.3) | 30.2 (1.1) | 3.9 (71.2) |
| Dark 1× X-ray dose 100 K -TCEP | 2.00 (2.03) | 93.4 (91.9) | 4.5 (3.8) | 19.0 (1.0) | 4.8 (85.9) |
| Dark 2× X-ray dose 100 K -TCEP | 2.11 (2.15) | 88.8 (82.8) | 4.2 (3.9) | 12.2 (1.0) | 5.7 (94.5) |
| Dark 3× X-ray dose 100 K -TCEP | 2.11 (2.15) | 94.2 (90.8) | 3.9 (3.5) | 21.2 (1.0) | 4.0 (91.4) |

<sup>1</sup>High-resolution shell is between the values inside and outside the parentheses.
<sup>2</sup>The value in the high-resolution shell is in parentheses.
<sup>3</sup>Average number of observed diffraction spots per unique reflection.

**Table S3.**
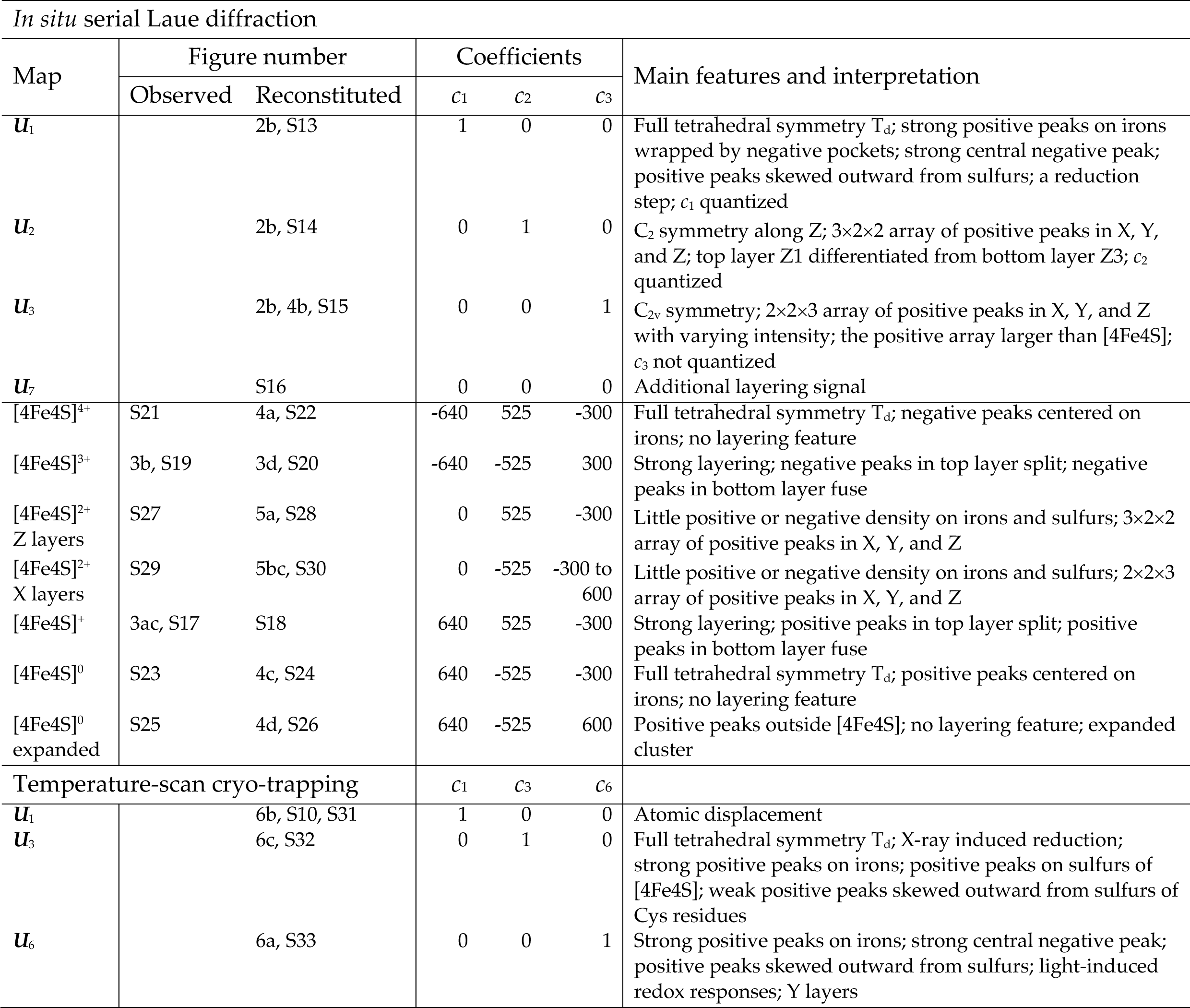
Component maps, observed, and reconstituted difference maps

## Supplementary Figures and Legends

**Figure S1.**
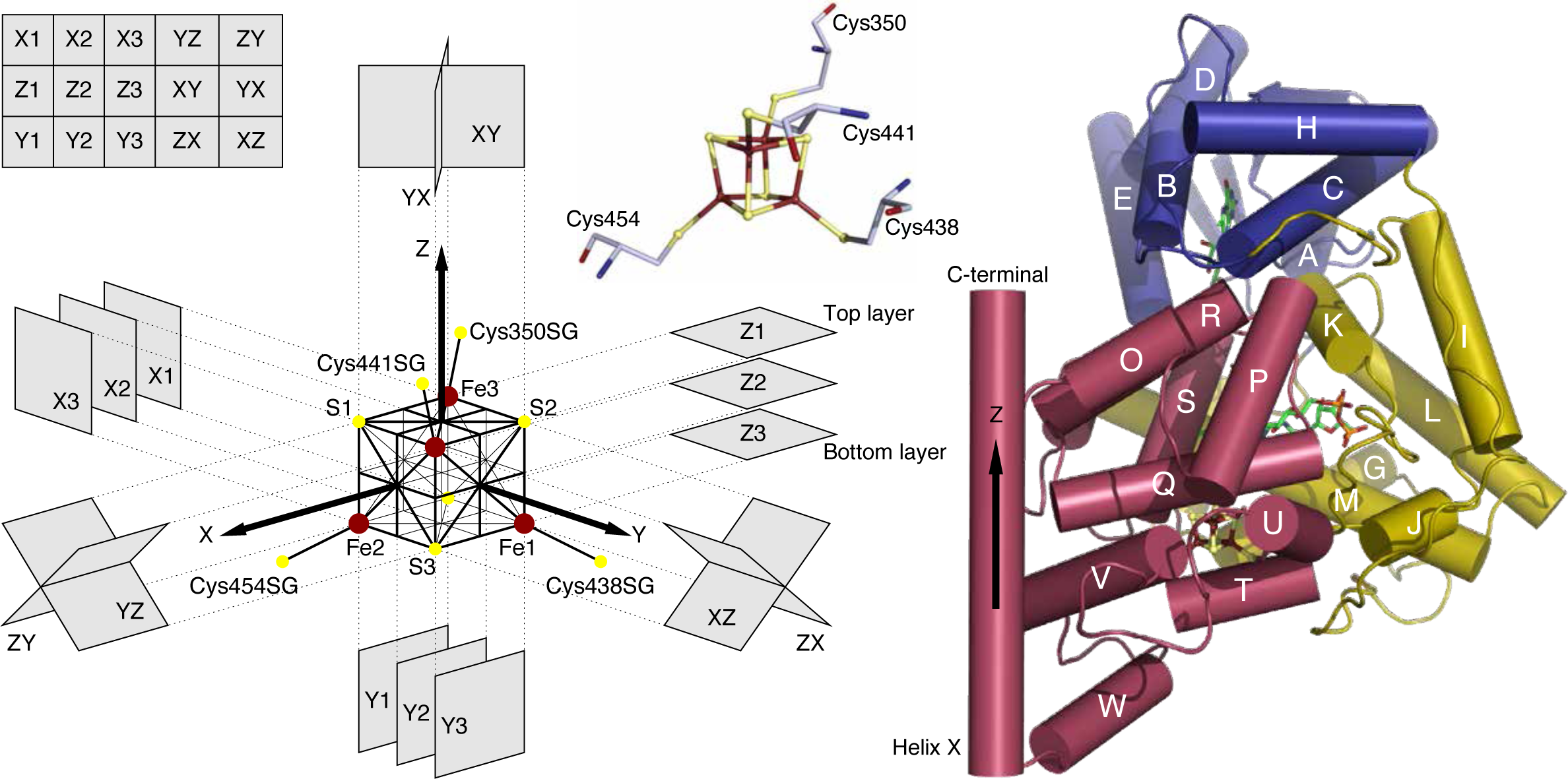
Orientations in PhrB structure. The iron-sulfur cluster [4Fe4S] is incorporated in the C-terminal domain of PhrB in red. The geometry of [4Fe4S] can be considered as two nested cubes that share the same center. Four irons are located on the corners of the smaller cube with an edge length of 1.90 Å. Four sulfurs are on the opposite corners of the larger cube with an edge length of 2.44 Å. Here the difference in the cubic size is omitted for clarity in the schematic drawing. The C-terminal direction of the last helix X is approximately perpendicular to one face of [4Fe4S], and therefore is selected as Z axis. The normal directions of the other two faces of [4Fe4S] are X and Y axes according to the righthand rule. Cross sections through [4Fe4S] are defined as the mean plane of the corresponding faces of the nested cubes. Images of electron density map on these cross sections are tiled in an array as shown in the upper left corner. Additional cross sections perpendicular to the body diagonals of [4Fe4S] are further defined in Fig. S13. Three-fold axes are located along the body diagonals, thus perpendicular to these cross sections.

**Figure S2.**
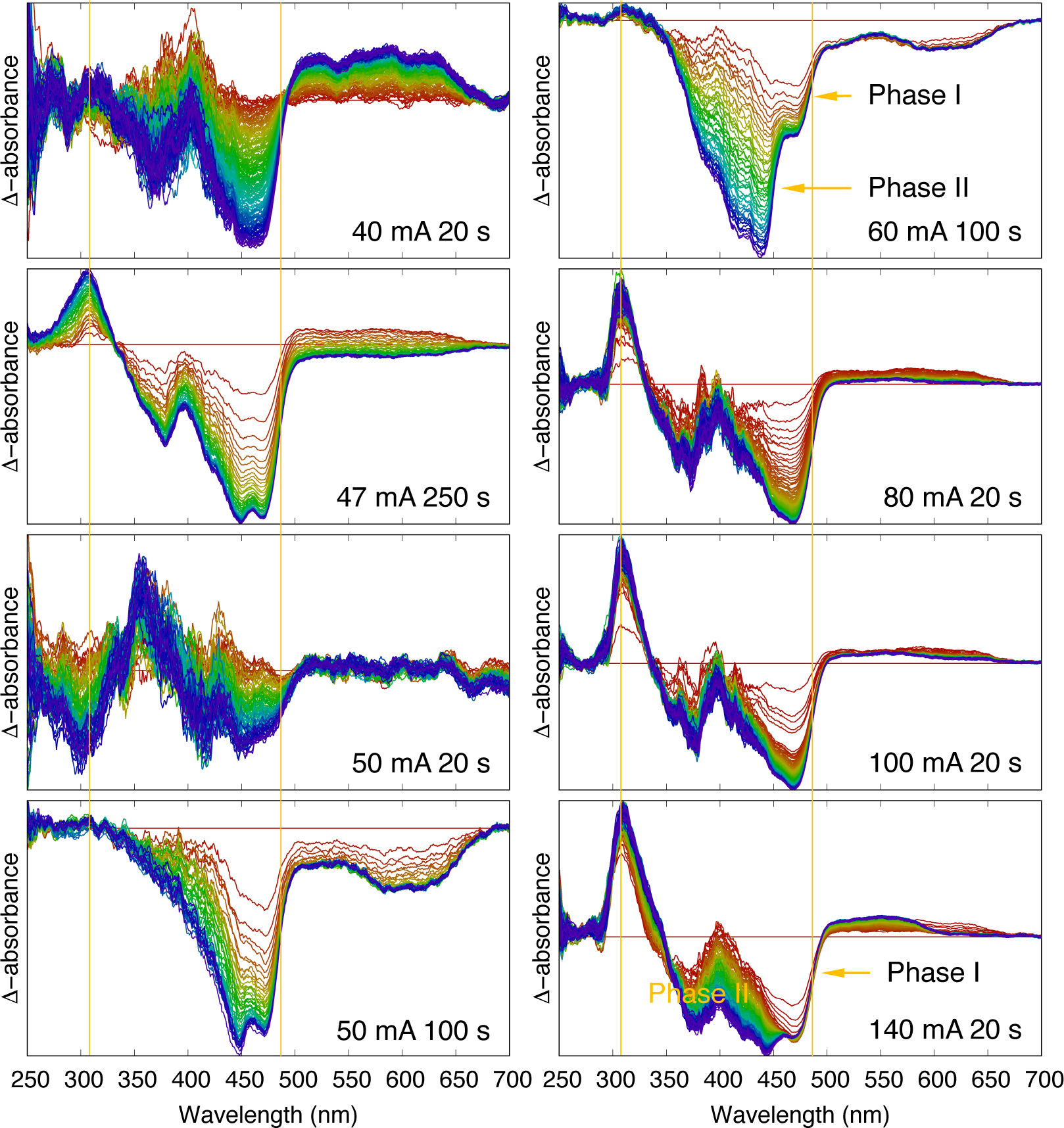
Difference absorption spectra of wildtype PhrB crystals. Wildtype PhrB crystals are illuminated by 405 nm light at different power densities (Fig. S4) and duration. Each spectrum is evenly spaced in a time series from red to blue during the period of illumination as marked (Fig. S3). The biphasic photoreductions of FAD and DMRL were previously studied (Ren et al., 2022). Some panels here clearly show two phases of the absorbance loss. Phase I present in all time series demonstrates the loss of the fully oxidized FAD that features the downward edge at 485 nm marked by vertical lines. During phase II, DMRL is being photoreduced, which is only obvious in some time series after a dimmer but prolonged illumination (top right) or under a more intense light condition (bottom right). The downward edge at 485 nm clearly stops developing in phase II indicating no further loss of FAD despite the continued illumination, which was interpreted as an active oxidation of the hydroquinone FADH^-^ back to the catalytically inactive form FAD by the photoreduction of DMRL (Ren et al., 2022). A sharp gain of absorbance at 308 nm with full width at half maximum (FWHM) of 25 nm as marked by the vertical lines tends to be stronger under higher power density. However, this is not consistent under lower light intensities, sometimes could have no change (bottom left) or even become an absorption loss (third left). This observation suggests that the [4Fe4S] cluster plays the role of a high-capacity electron buffer that absorbs excess electrons and supplies them back as needed (Ren et al., 2022).

**Figure S3.**
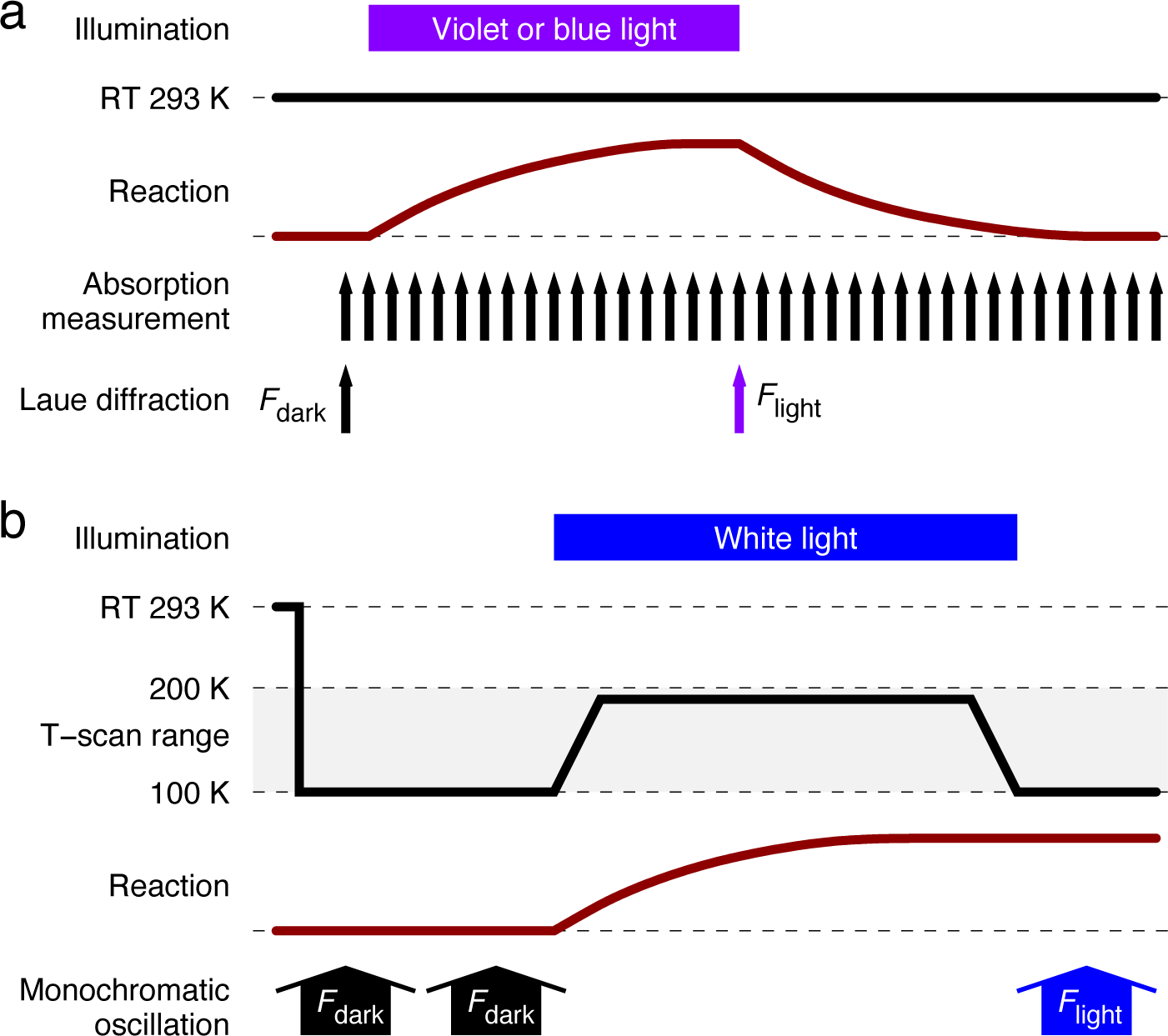
Experimental timing. (a) Absorption spectroscopy and Laue diffraction. Absorption spectra are repeatedly measured during illuminations of violet light at 405 nm (Fig. S2). Laue exposures are taken before an illumination starts for a dark dataset and at the end of an illumination of 2 s for light datasets. (b) Temperature scan cryo-trapping. A dark dataset is taken from one end of a crystal before any illumination at 100 K. The temperature of the crystal is elevated to a setting ranging from 100 to 200 K while the crystal is illuminated by flooded white light for 15 minutes at a high power density ranging from 2 to 3.5 mW/mm^2^ or a low power density of a few μW/mm^2^ (Fig. S4). The crystal is spinning during the illumination. A light dataset is taken from a fresh segment of the same crystal after the illumination and the temperature returns to 100 K. Additional dark datasets could be taken from the same crystal segment as controls to evaluate the effect of X-ray radiation damage. Additional light datasets could be taken if more fresh segments are available on the same crystal after more illumination cycles.

**Figure S4.**
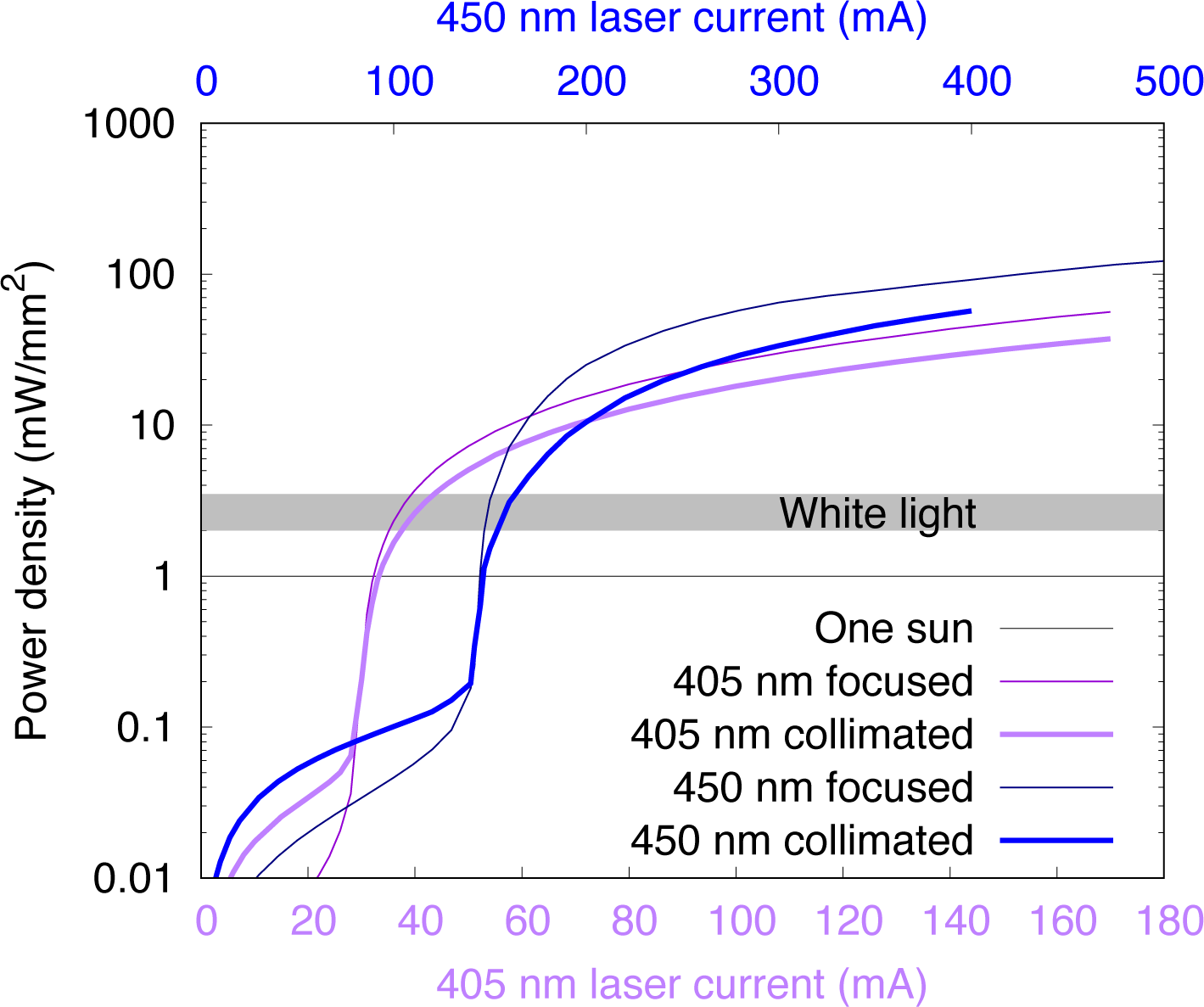
Power density of illumination. The power density as function of the current setting of the laser diodes are measured with an optical power meter (Thorlabs, Inc.). The power density of 1 mW/mm^2^ equals to one sun of 1 kW/m^2^, roughly the power density of the solar irradiation on the surface of the earth. Collimated beams are used for spectroscopic measurements. Focused beams are used to illuminate crystals grown on chip in serial Laue diffraction. The focused beams are only modestly more intense than the collimated beams, because most of the photons emitted from the laser diodes in a cone are trimmed to form a smaller focal spot of ∼300 μm FWHM in diameter. Flooded white light is used to illuminate crystals in temperature scan cryo-trapping experiments. Its power density ranges from 2 to 3.5 mW/mm^2^ depending on the setup. The low light illumination in temperature scan cryo-trapping experiments spans a few μW/mm^2^.

**Figure S5.**
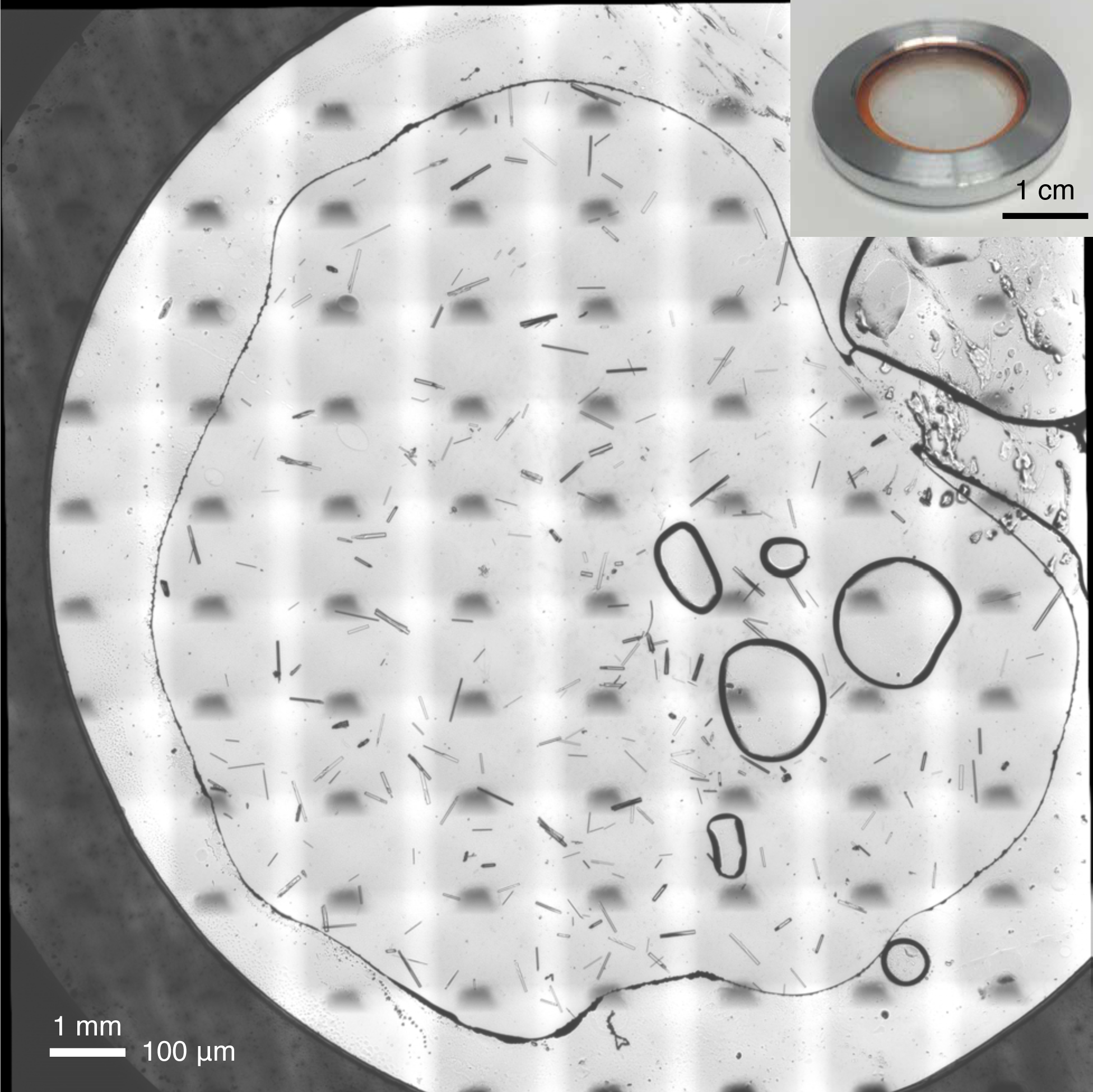
PhrB crystals grown on a chip for *in situ* diffraction. Many infrared micrographs are tiled together to provide a detailed image of crystals grown on chip at a resolution of 1.34 μm/pixel. The peripheral dark area is a shim that determines the thickness of the crystallization chamber. The circular area is the diffraction window. An assembled crystallization device is displayed in the inset (Biju et al., 2022; Ren et al., 2018, 2020). The shim in orange is visible.

**Figure S6.**
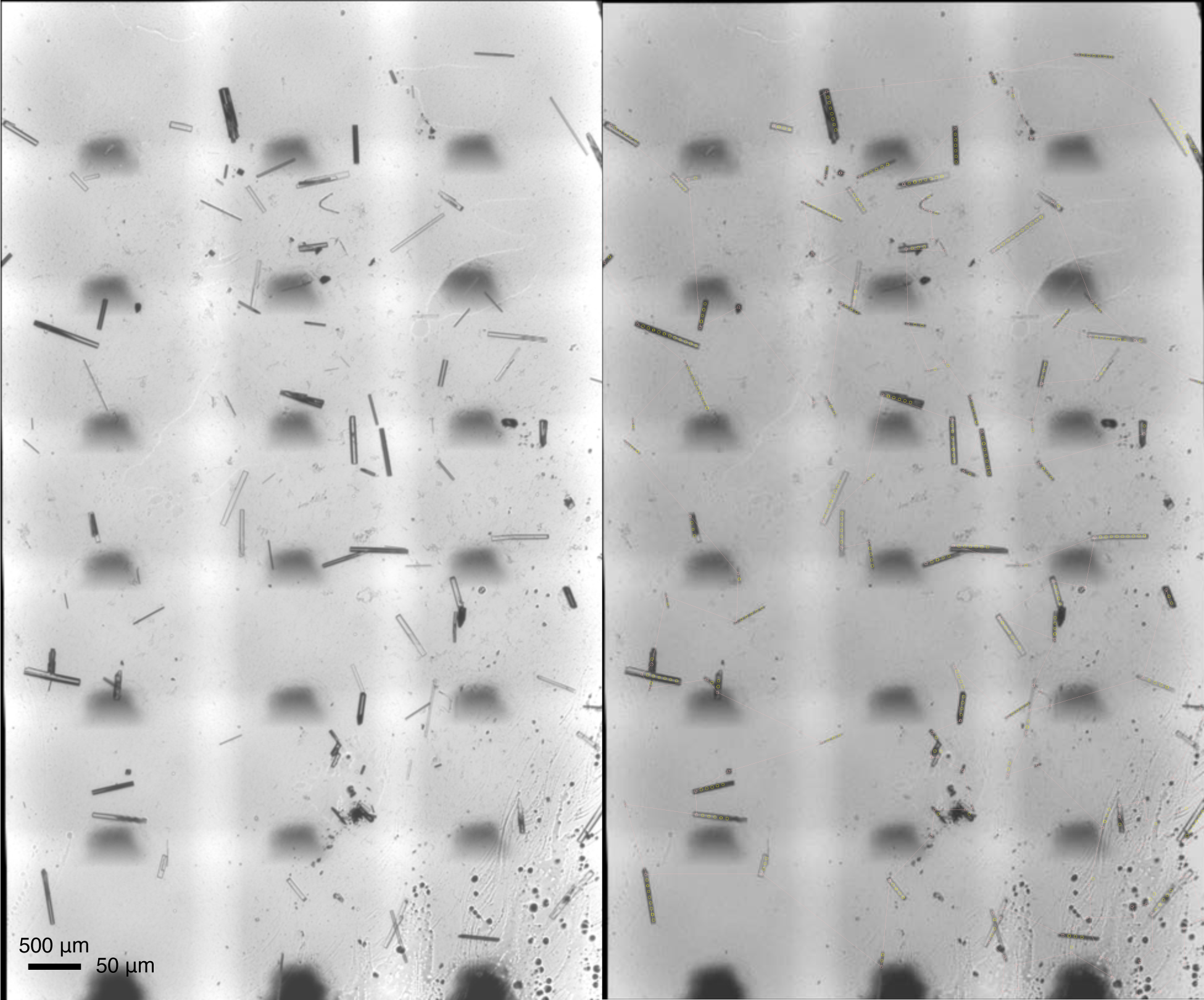
Crystal recognition and shot planning. The montage of infrared micrographs (left) is analyzed for crystal recognition. Shots are planned along the crystals in a rod shape depending on a user input of X-ray beam size as marked in small circles. The translocation route in pink is computed as the solution to the traveling salesman problem (Ren et al., 2020).

**Figure S7.**
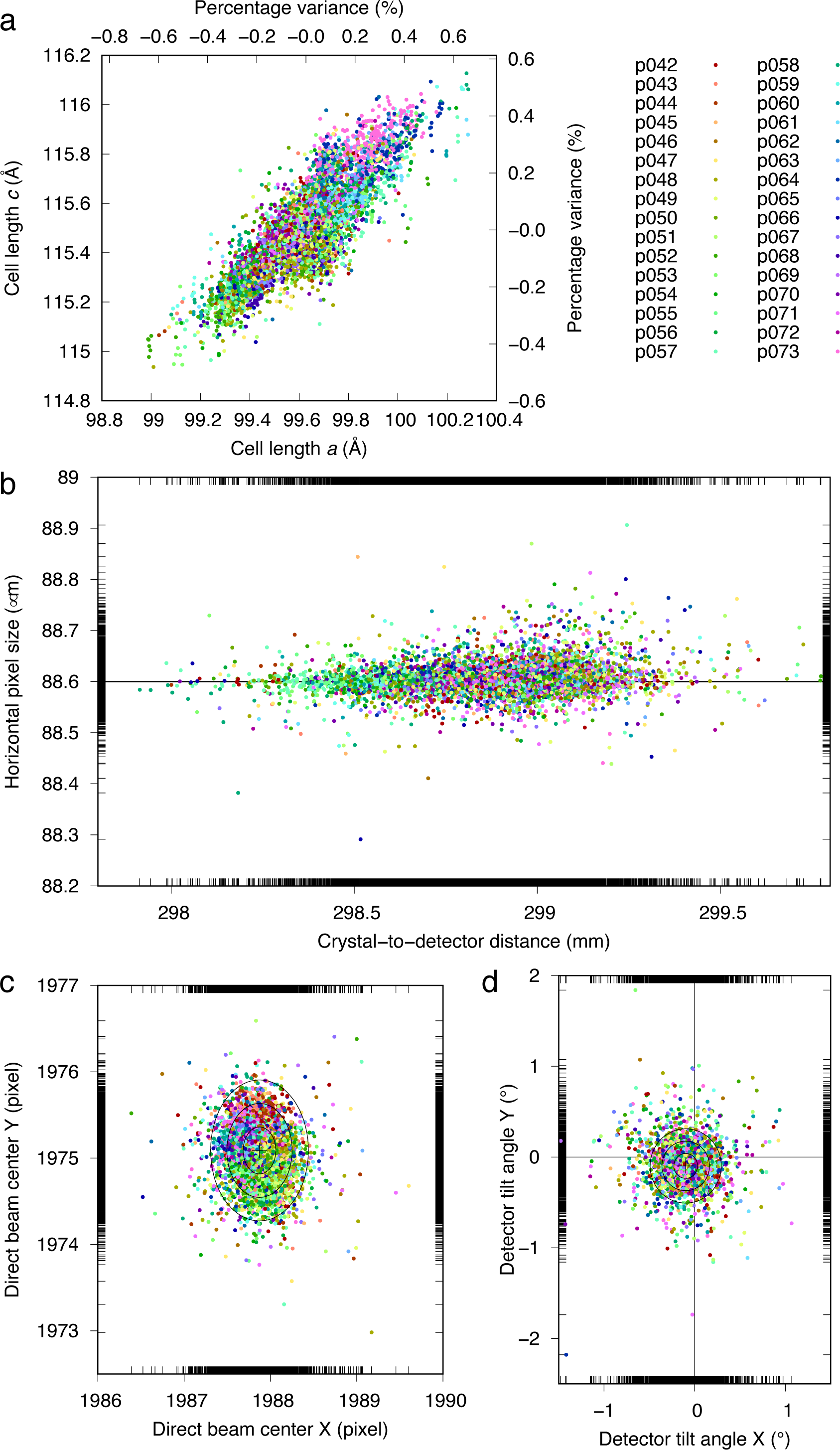
Geometric refinement of Laue diffraction patterns. Unit cell parameters and several geometric parameters of the diffractometer, such as crystal-to-detector distance, pixel size of the detector, direct beam center on the diffraction image, and detector tilt angles, are refined for each diffraction image. (a) The refined unit cell lengths *a* and *c* are plotted against each other. Each dot represents the refined values from one Laue diffraction pattern. Patterns derived from each crystallization device is distinguished by a different color. Unit cell length *b* is not refined due to the lack of a precise length calibration in polychromatic Laue diffraction. It is normal that the unit cell lengths show a correlation rather than an anti-correlation. An anti-correlation indicates a distortion of unit cell shape, thus more severe non-isomorphism. (b, c, and d) Several detector parameters are also refined. The ellipses mark 1, 2, and 3*σ* of the distributions.

**Figure S8.**
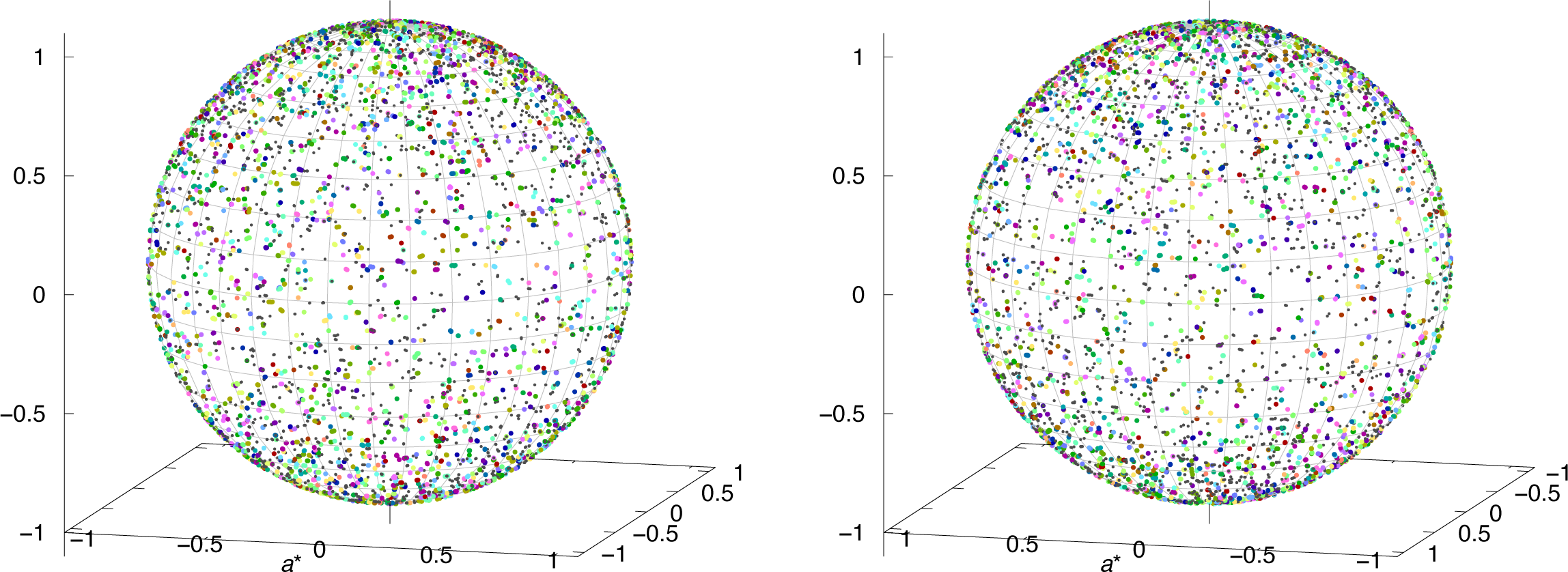
Orientation distribution of crystals. Each colored dot represents the orientation derived from the unit vector ***a***\* = (1, 0, 0) in the reciprocal space pre-multiplied by the refined mis-setting orientation matrix of a crystal on chip. Crystals from each crystallization device is distinguished by a different color. A black dot represents the opposite vector of a colored dot, the orientation from which the Friedel mates could be collected. This plot is used to evaluate whether a preference of orientation exists in all crystals on chip.

**Figure S9.**
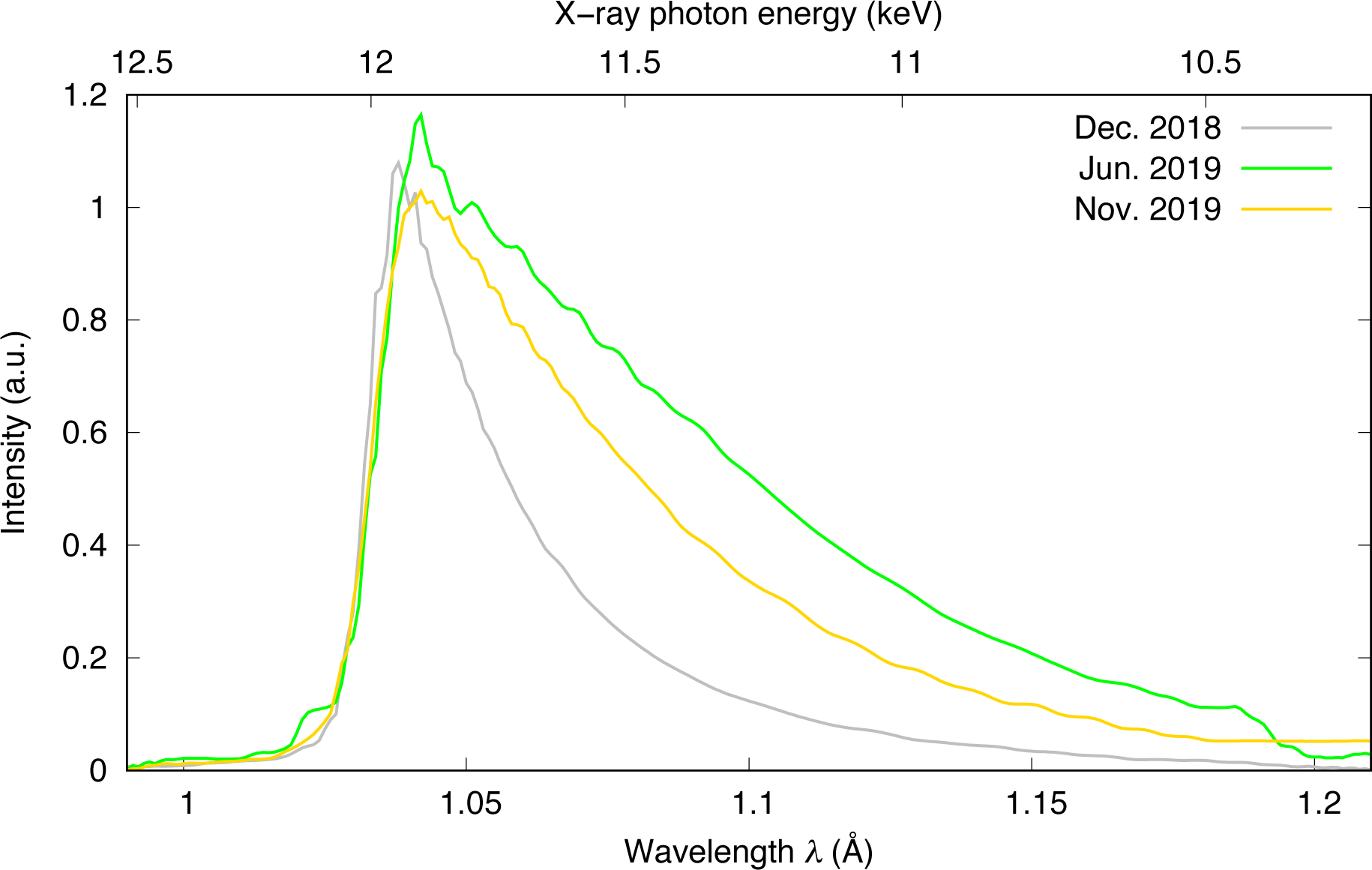
Wavelength normalization curves. These wavelength normalization curves are derived from the Laue data scaling software Epinorm™ (Ren, 2006, 2008). The quality of wavelength normalization is an important indication of the data quality. The incident spectra of the X-rays appear to vary significantly from one run to another.

**Figure S10.**
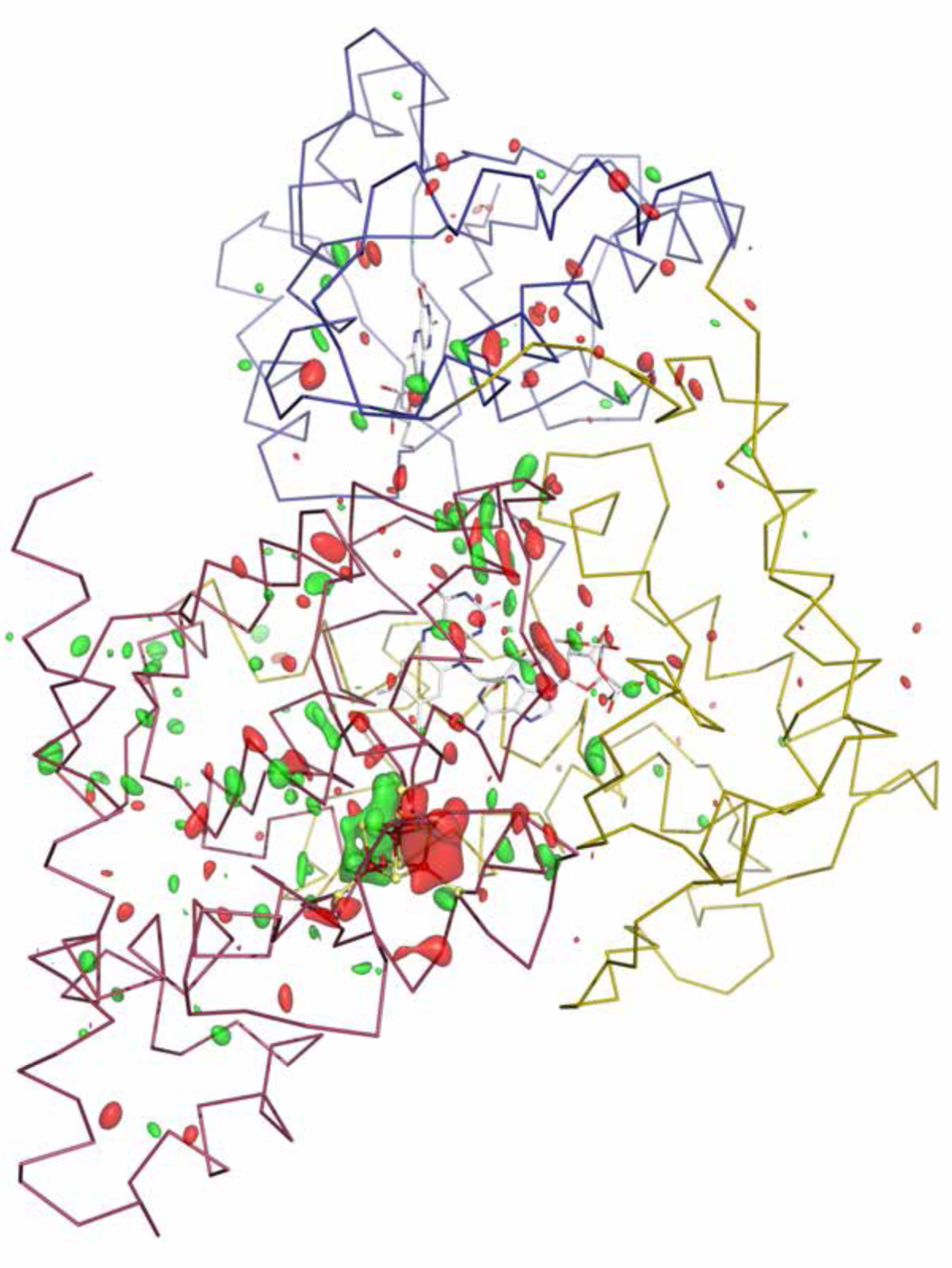
Overall signal from temperature scan cryo-trapping. The first component from SVD of light – dark difference maps (Methods) is contoured at ±4*σ* in green and red, respectively. The strongest difference signal at ±9*σ* is located at the [4Fe4S] cluster.

**Figure S11.**
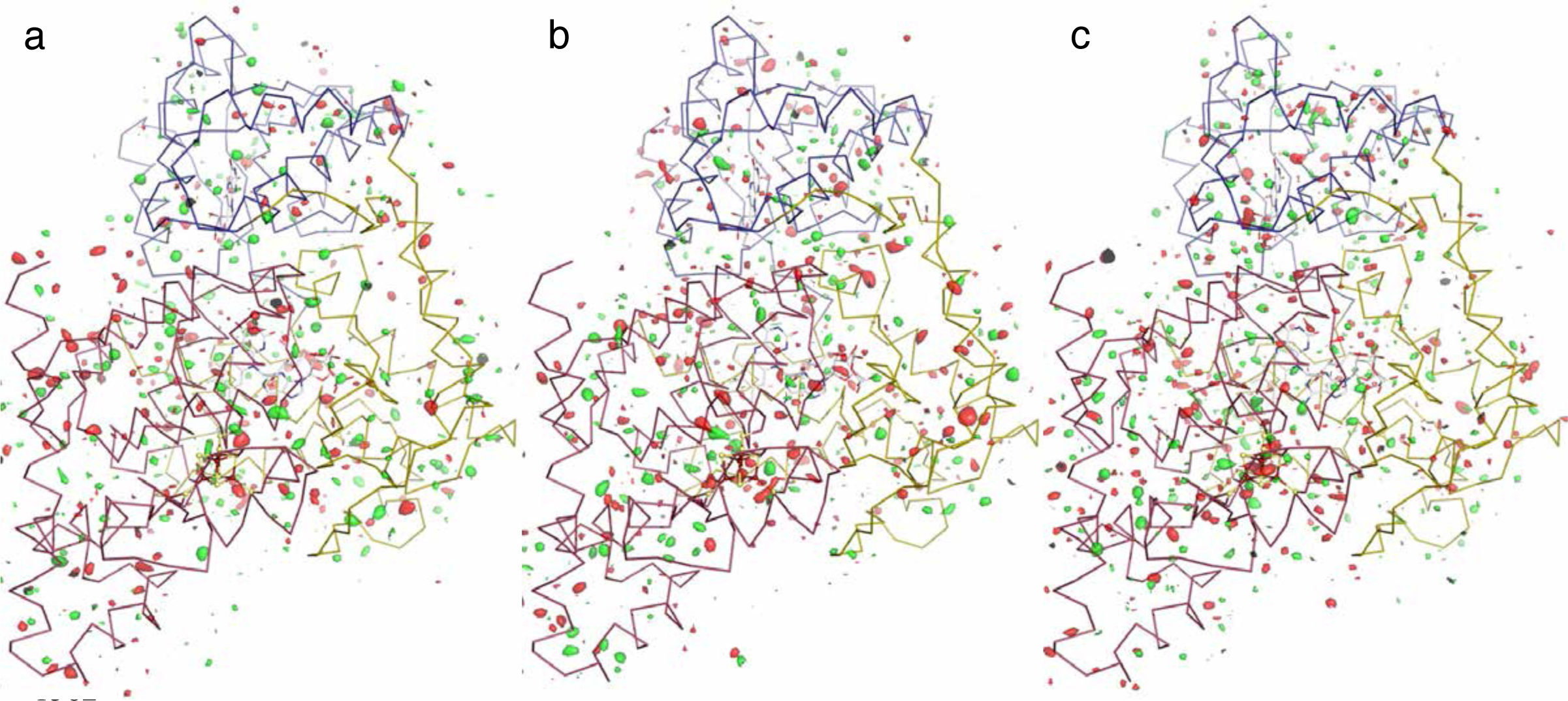
Dark-minus-dark difference Fourier maps at room temperature over the entire protein molecule. Each map is contoured at ±3*σ* in green and red, respectively. The map in (c), and arguably the map in (b) as well, show some clustered difference electron densities at the [4Fe4S] cluster. These redox signals could be induced by X-rays or less strict control of visible light. Nevertheless, the signal-to-noise ratio shown in light-minus-dark difference maps (Fig. 1) is outstanding compared to these dark-minus-dark maps.

**Figure S12.**
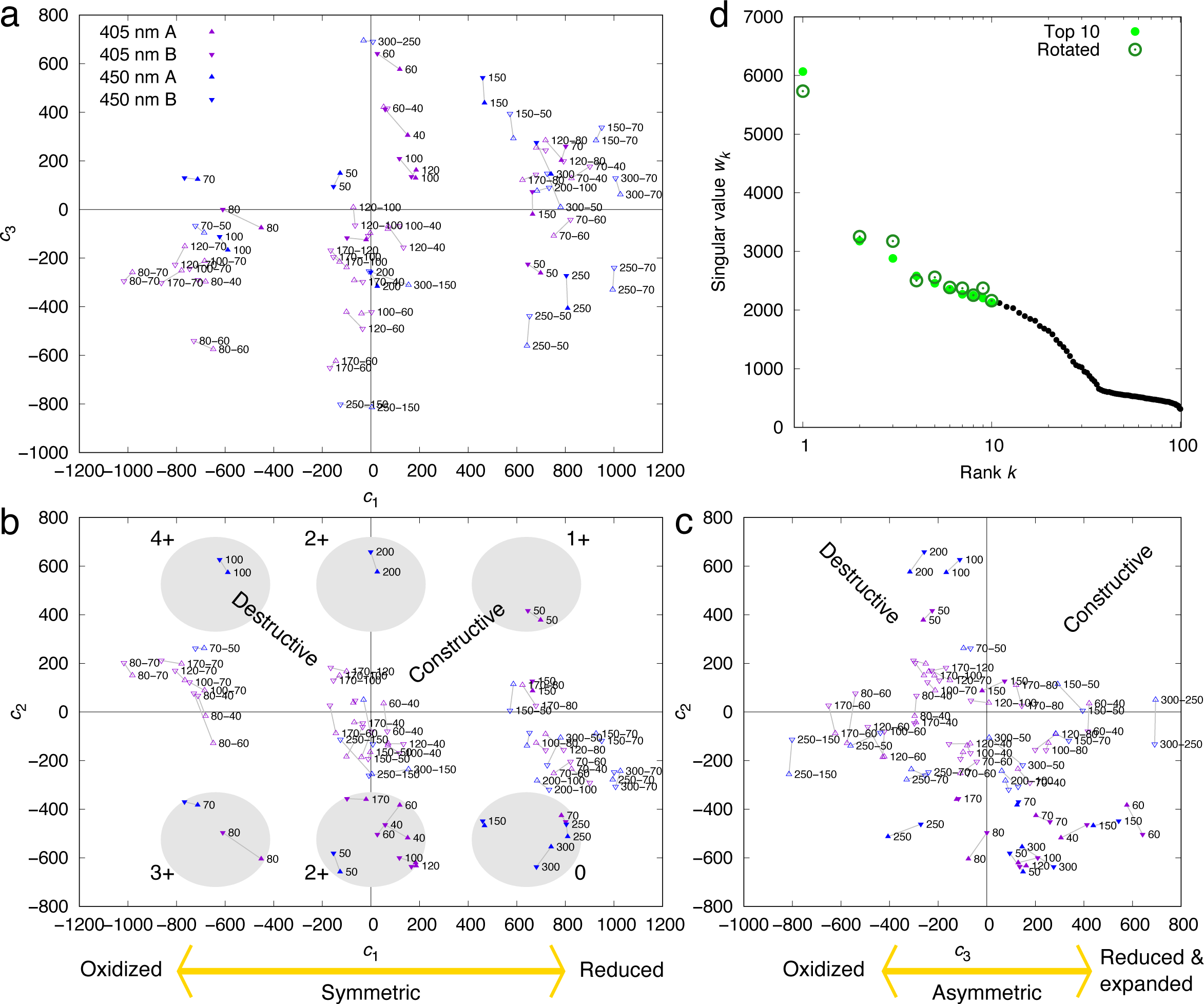
SVD analysis of the light-minus-dark and light-minus-light difference maps. Light datasets are collected after a 2-second illumination of violet or blue light from laser diodes at various power densities (Methods). Dark datasets are collected without illumination. As a result, each observed difference map is represented as a linear combination of several major components *c*_1_***U***_1_ + *c*_2_***U***_2_ + *c*_3_***U***_3_ + … (Methods). Each component map ***U****_k_* (*k* = 1, 2, 3, …) captures the common features among the raw difference maps (Fig. 1). (a-c) Three orthographical views of the Euclidian space of the first three dimensions. Light – dark and light – light difference maps are marked with solid and open triangles, respectively. Maps from two chains A and B related by the noncrystallographic symmetry are in up- and down-pointing triangles and linked with a gray line. The electric current setting of the laser diode is labeled wherever possible. See Fig. S4 for the corresponding power density of illumination. (d) Singular values before and after Ren rotation (Methods).

**Figure S13.**
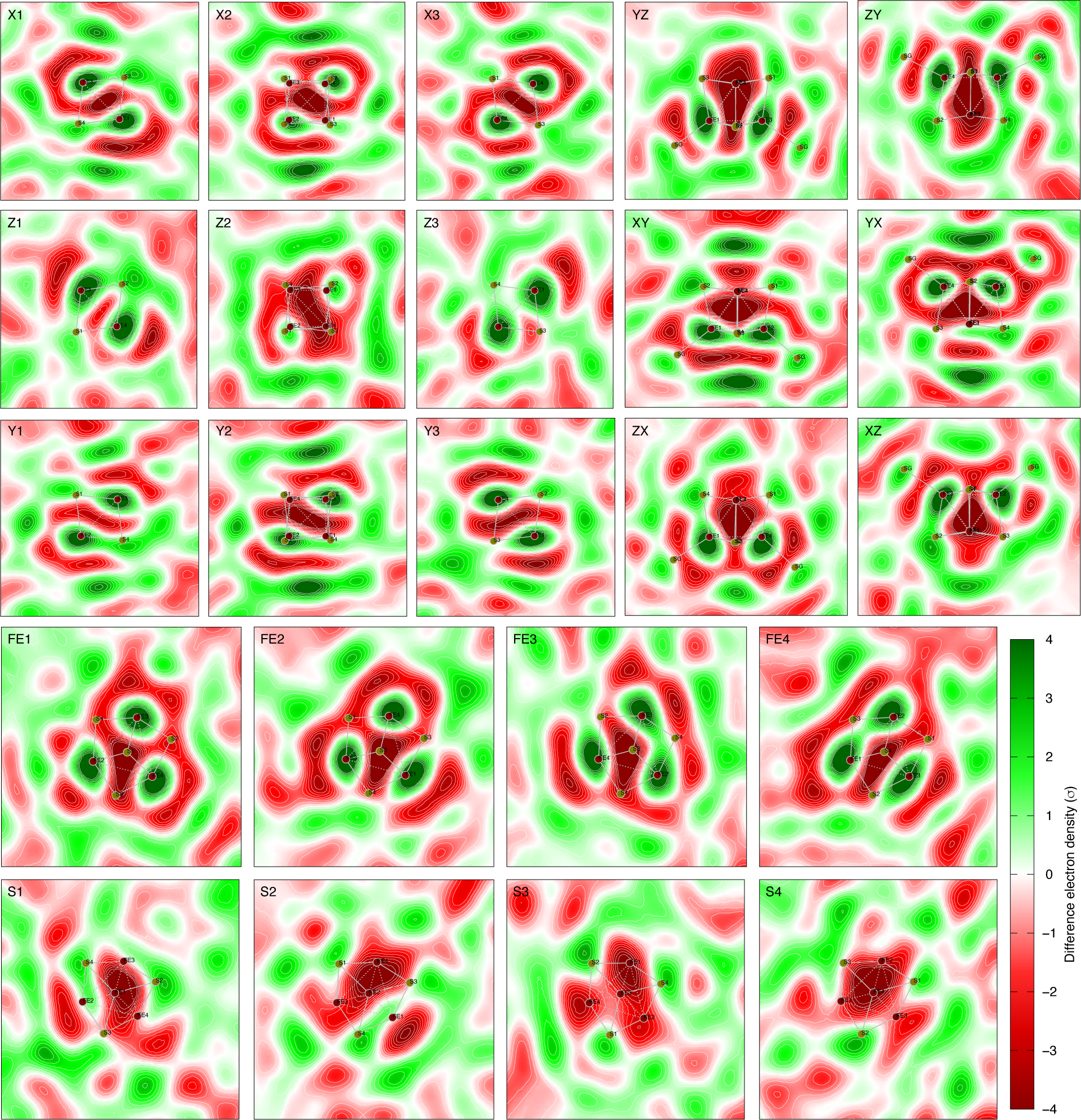
First component ***U***_1_ of difference maps at room temperature. The orthogonal cross sections are defined in Fig. S1. A cross section normal to a three-fold axis is marked by the atom number of an iron or a sulfur that is on the three-fold. Therefore, the other three atoms of the same type are related by the three-fold axis on the cross section. For example, the cross section marked FE1 is normal to the three-fold axis through FE1. Therefore, FE2, FE3, and FE4 are related by the three-fold axis on the cross section. Other atoms not on the cross section are projected to the cross section. Sγ atoms of the Cys residues are marked as SG, the atom identifier in the PDB file. Each panel displays an integration within a slab of 0.4 Å thick around the cross section, that is, a slab thick enough to include both irons and sulfurs around the same cubic face. ***U***_1_ largely follows a full tetrahedral symmetry, known as T_d_ in Schoenflies notation, that is, three two-fold axes normal to the cubic faces of [4Fe4S] and through their centers, four three-fold axes through the body diagonals, and six mirrors through opposite edges. A two-fold axis is perpendicular to each of the nine top left panels and located at the center. A mirror lies in each of the six top right panels. Another mirror is perpendicular to each of them and runs vertically through the center. A three-fold axis is perpendicular to each of the eight bottom panels and through the center. It is visible that the three-fold axes are not as strictly followed on the cross sections through the sulfurs. See Fig. 2b for a contour plot of this component.

**Figure S14.**
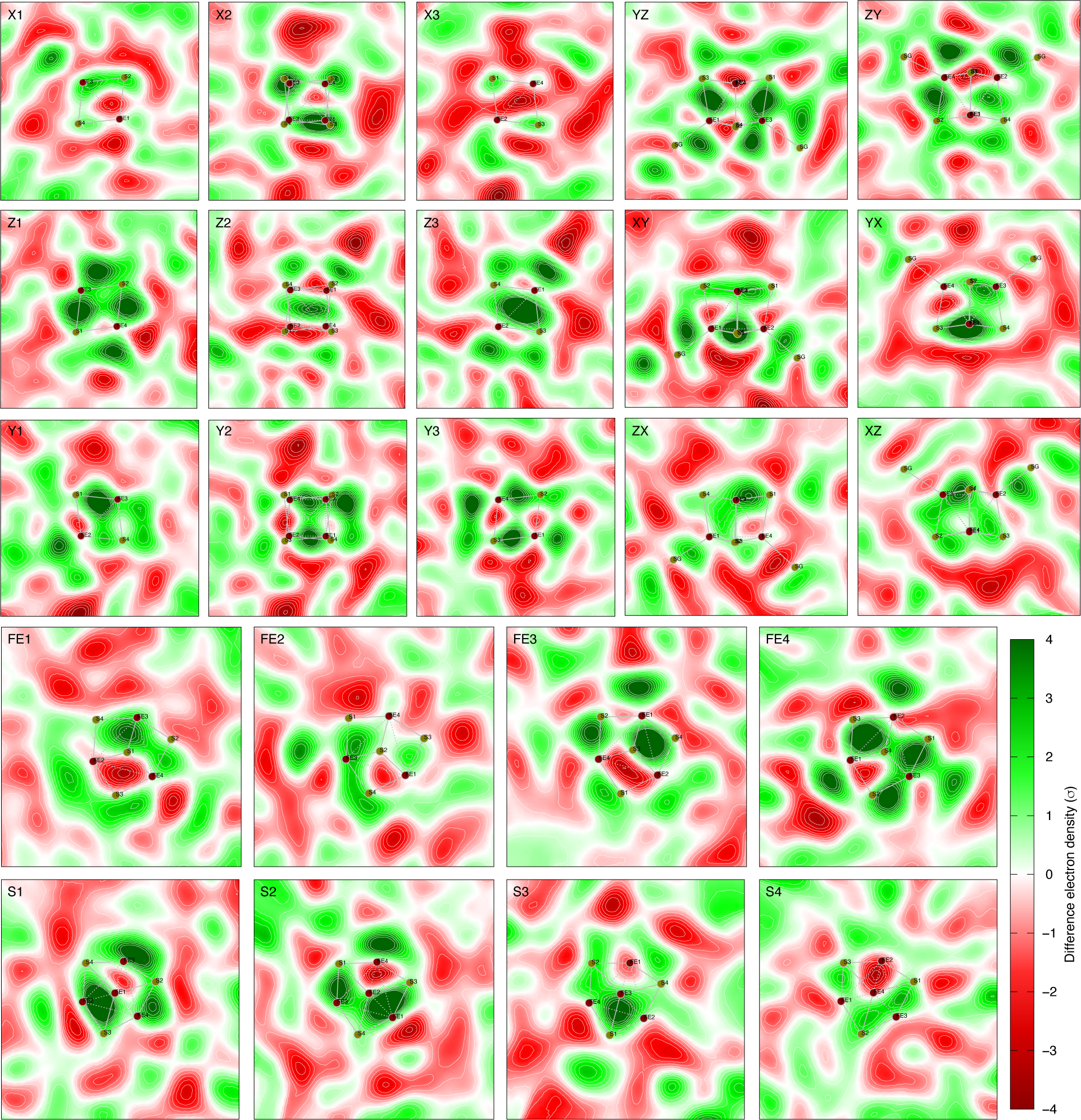
Second component ***U***_2_ of difference maps at room temperature. See Figs. S1 and S13 for the definitions of cross sections. Each panel displays an integration within a slab of 0.4 Å thick around the cross section. ***U***_2_ is approximately in C_2_ symmetry with the only two-fold axis along Z, that is, through the centers of Z1, Z2, and Z3 and normal to these cross sections. 12 positive peaks form a 3×2×2 array in X, Y, and Z directions, respectively. Two peaks on Z3 cross section are fused together while two peaks on Z1 are well separated. See Fig. 2b for a contour plot of this component.

**Figure S15.**
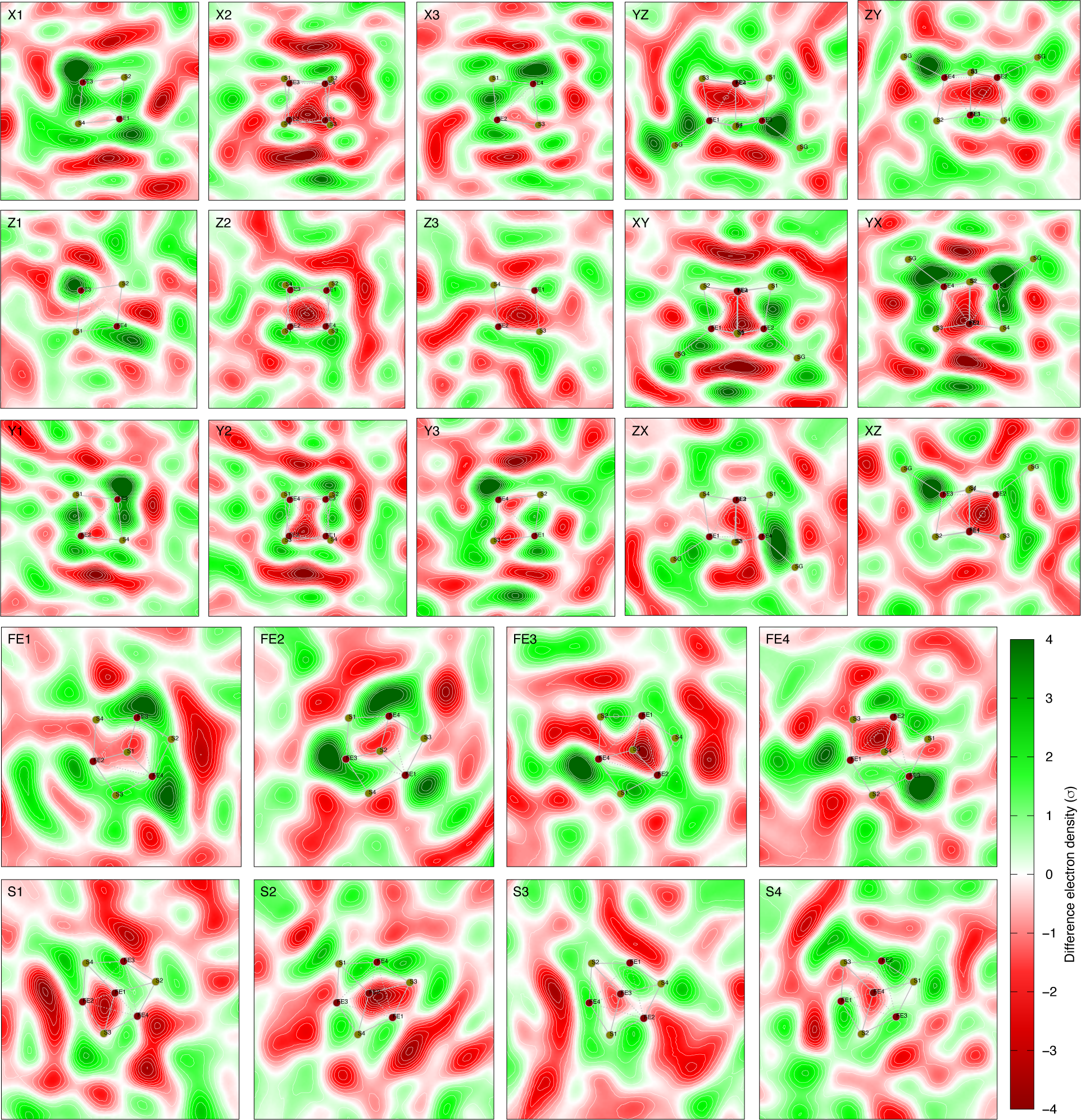
Third component ***U***_3_ of difference maps at room temperature. See Figs. S1 and S13 for the definitions of cross sections. Each panel displays an integration within a slab of 0.4 Å thick around the cross section. ***U***_3_ is approximately in C_2v_ symmetry with two perpendicular mirrors in XY and YX cross sections. Their intersection is a two-fold axis along Z, that is, through the centers of Z1, Z2, and Z3 and normal to these cross sections. 12 positive peaks form a 2×2×3 array in X, Y, and Z directions, respectively. This array is slightly larger than that in ***U***_2_. See Fig. 2b for a contour plot of this component. See Fig. 4b for integrations in thicker slabs of this component.

**Figure S16.**
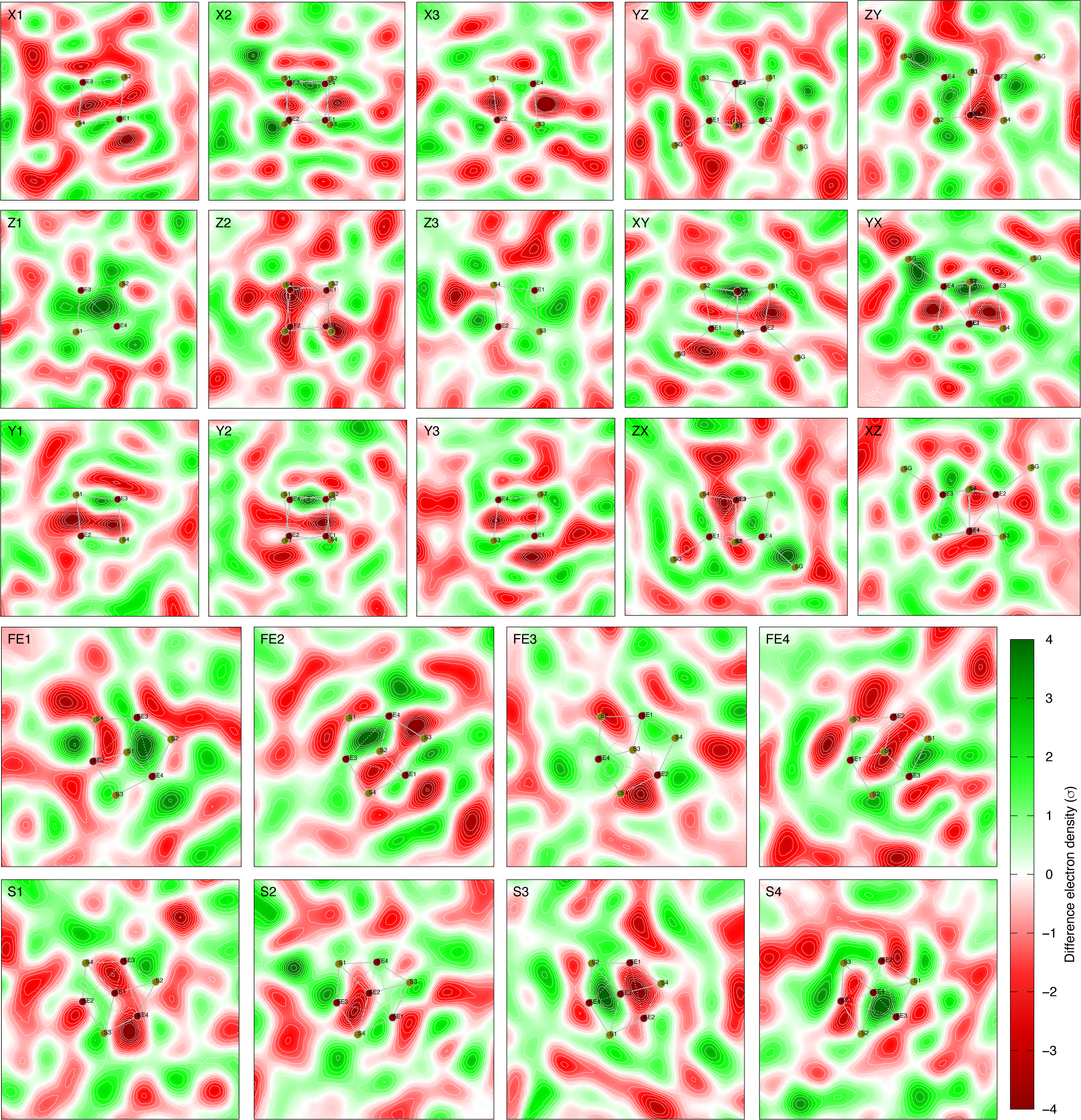
Seventh component ***U***_7_ of difference maps at room temperature. See Figs. S1 and S13 for the definitions of cross sections. Each panel displays an integration within a slab of 0.4 Å thick around the cross section. ***U***_7_ shows some additional structure of layering, but with no obvious symmetry.

**Figure S17.**
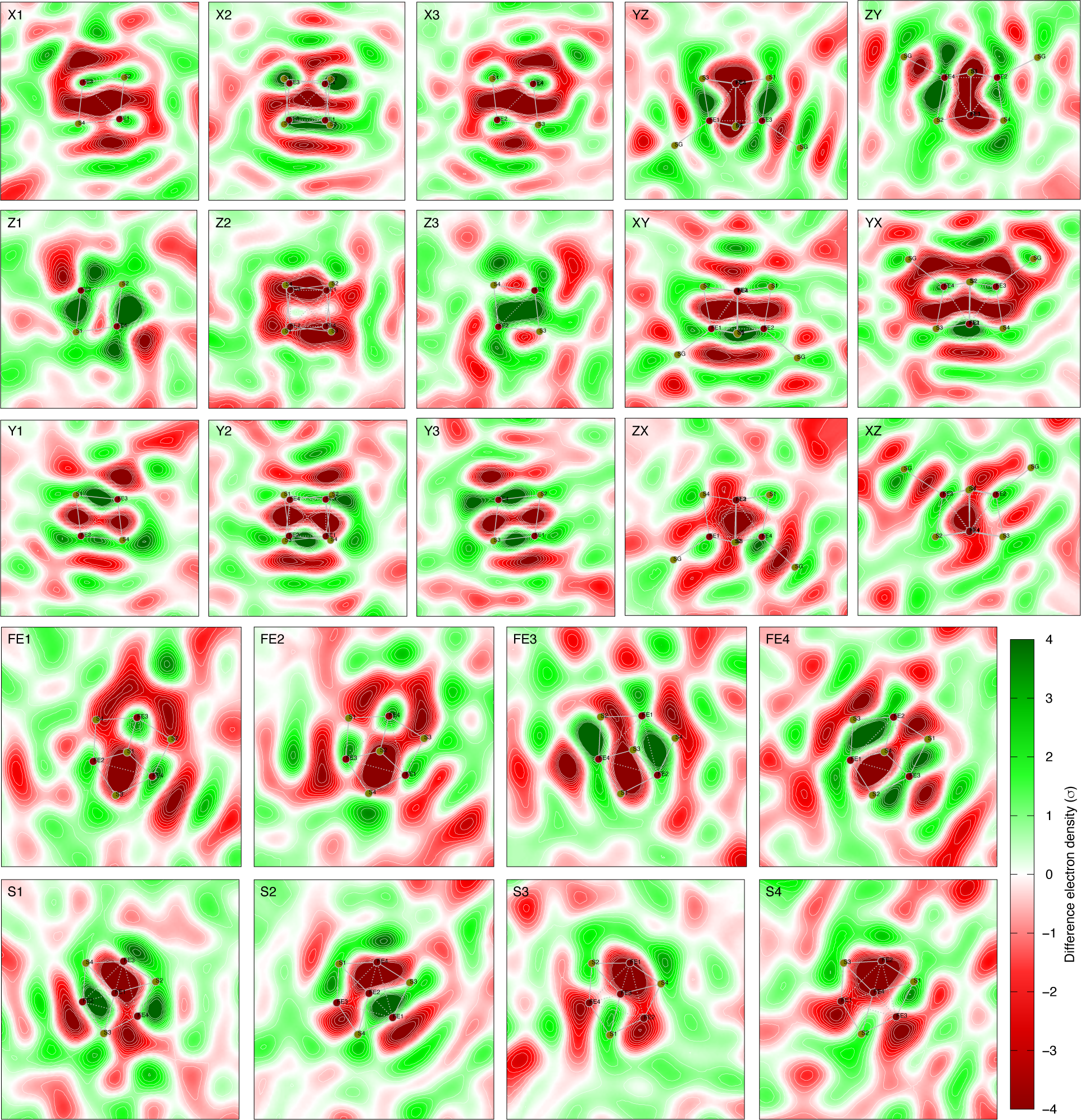
Raw difference map observed by illumination of 405 nm at 50 mA. See Figs. S1 and S13 for the definitions of cross sections. Each panel displays an integration within a slab of 0.4 Å thick around the cross section. This observed difference map demonstrates strong signal of valence layers and represents the reduced cluster [4Fe4S]^+^. See Fig. 3a for a three-dimensional contour of this map. See Fig. S18 for a reconstituted map that fits this observed map.

**Figure S18.**
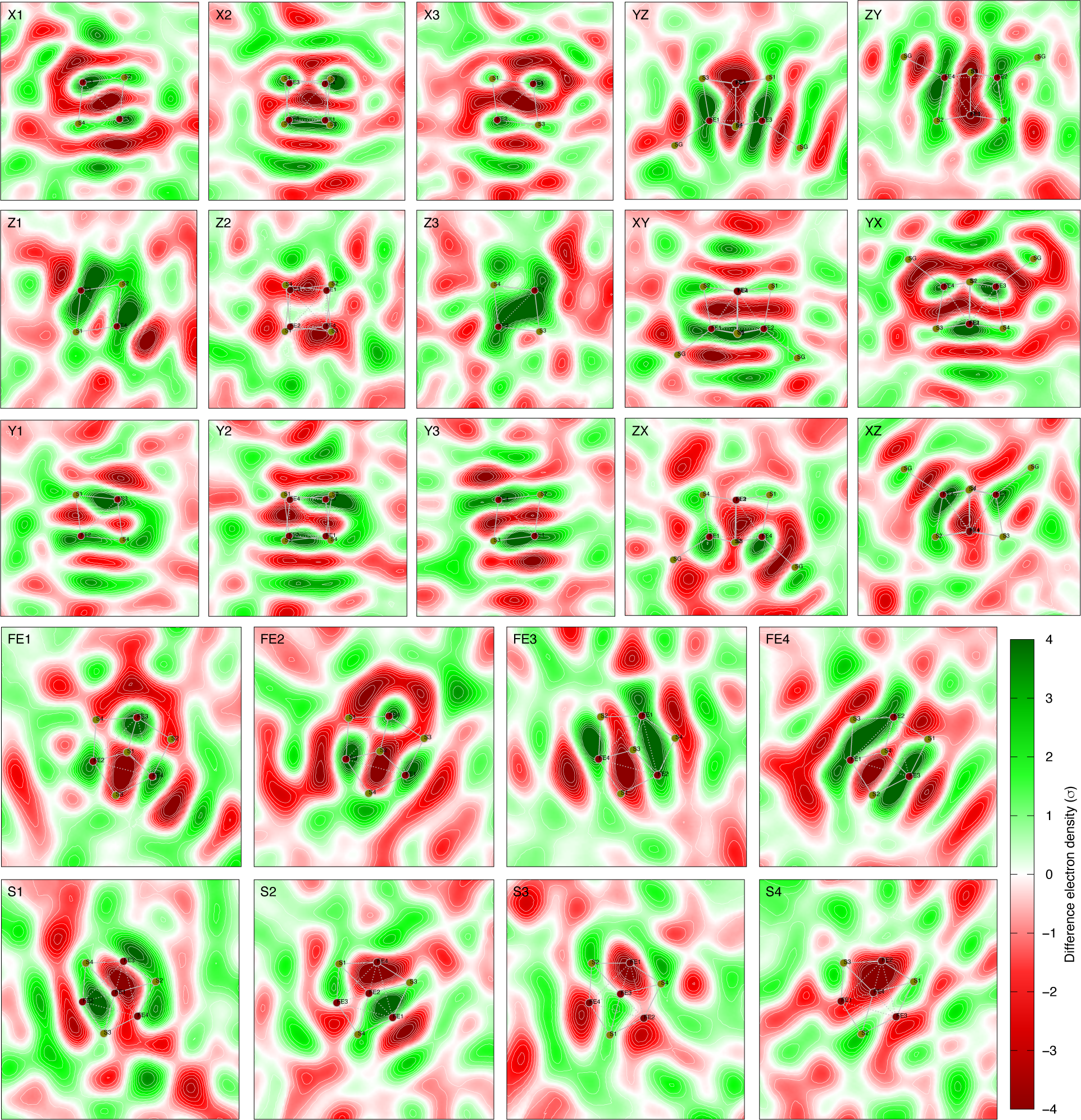
Reconstituted difference map of [4Fe4S]^+^ – [4Fe4S]^2+^ = 640***U***_1_ + 525***U***_2_ - 300***U***_3_. See Figs. S1 and S13 for the definitions of cross sections. Each panel displays an integration within a slab of 0.4 Å thick around the cross section. The raw difference map corresponding to this reconstitution is shown in Figs. 3ac and S17.

**Figure S19.**
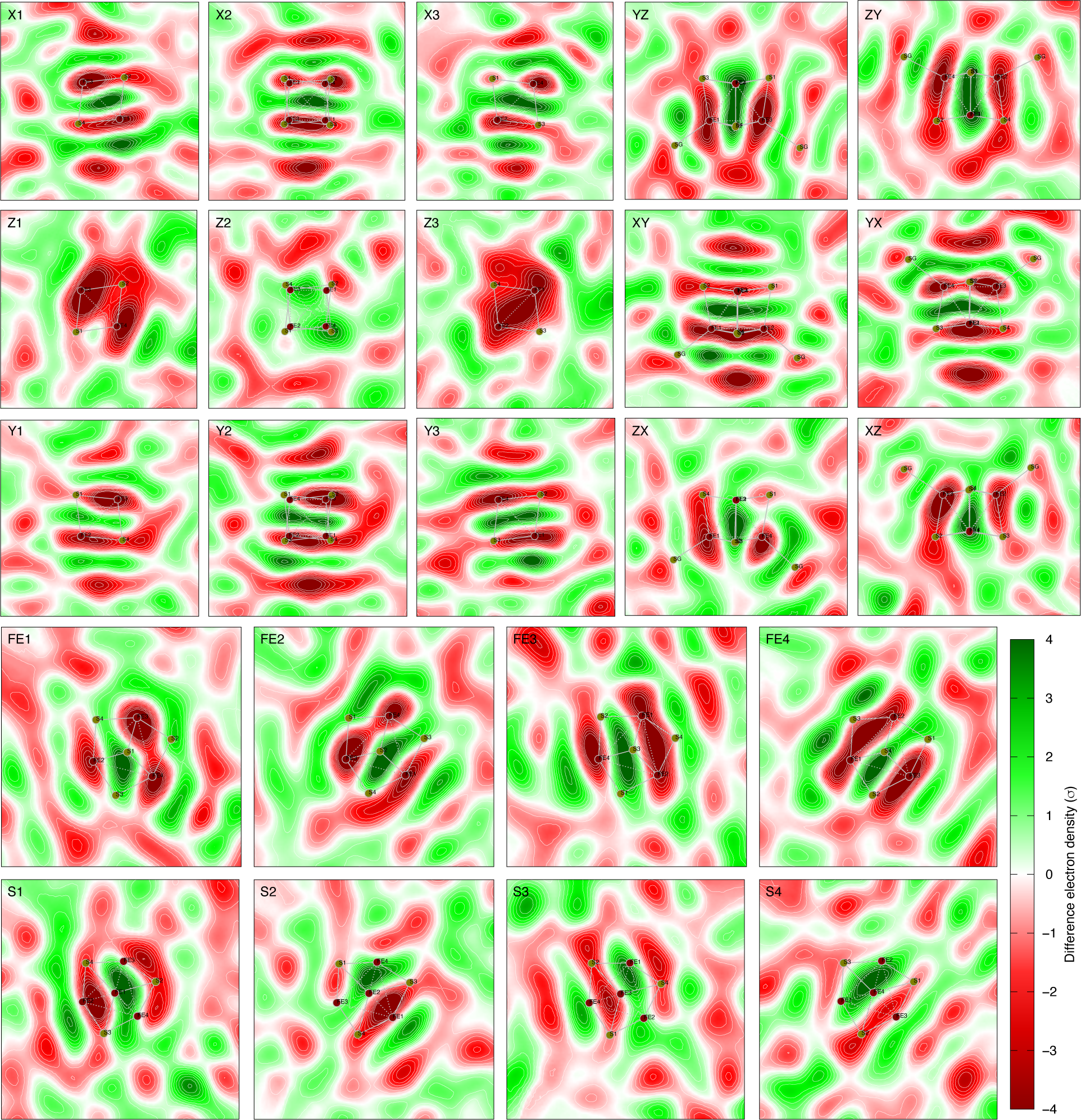
Raw difference map observed by illumination of 450 nm at 70 mA. See Figs. S1 and S13 for the definitions of cross sections. Each panel displays an integration within a slab of 0.4 Å thick around the cross section. This observed difference map also demonstrates strong signal of valence layers but largely opposite to that in Fig. S17. This observed map represents the oxidized cluster [4Fe4S]^3+^. See Fig. 3b for a three-dimensional contour of this map. See Fig. S20 for a reconstituted map that reproduces this observation.

**Figure S20.**
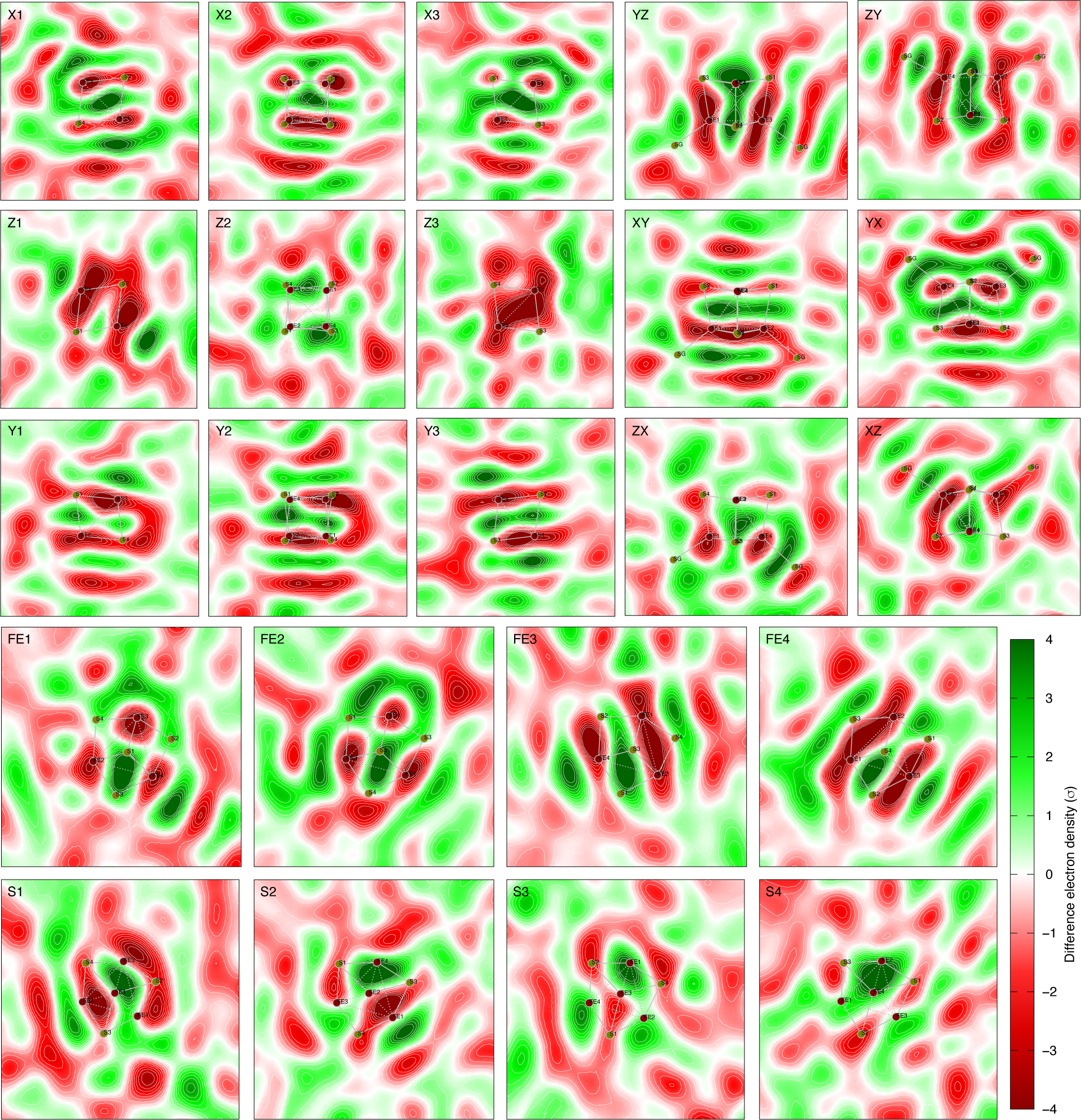
Reconstituted difference map of [4Fe4S]^3+^ – [4Fe4S]^2+^ = -640***U***_1_ - 525***U***_2_ + 300***U***_3_. See Figs. S1 and S13 for the definitions of cross sections. Each panel displays an integration within a slab of 0.4 Å thick around the cross section. The raw difference map corresponding to this reconstitution is shown in Figs. 3b and S19. This is a negation of Fig. S18.

**Figure S21.**
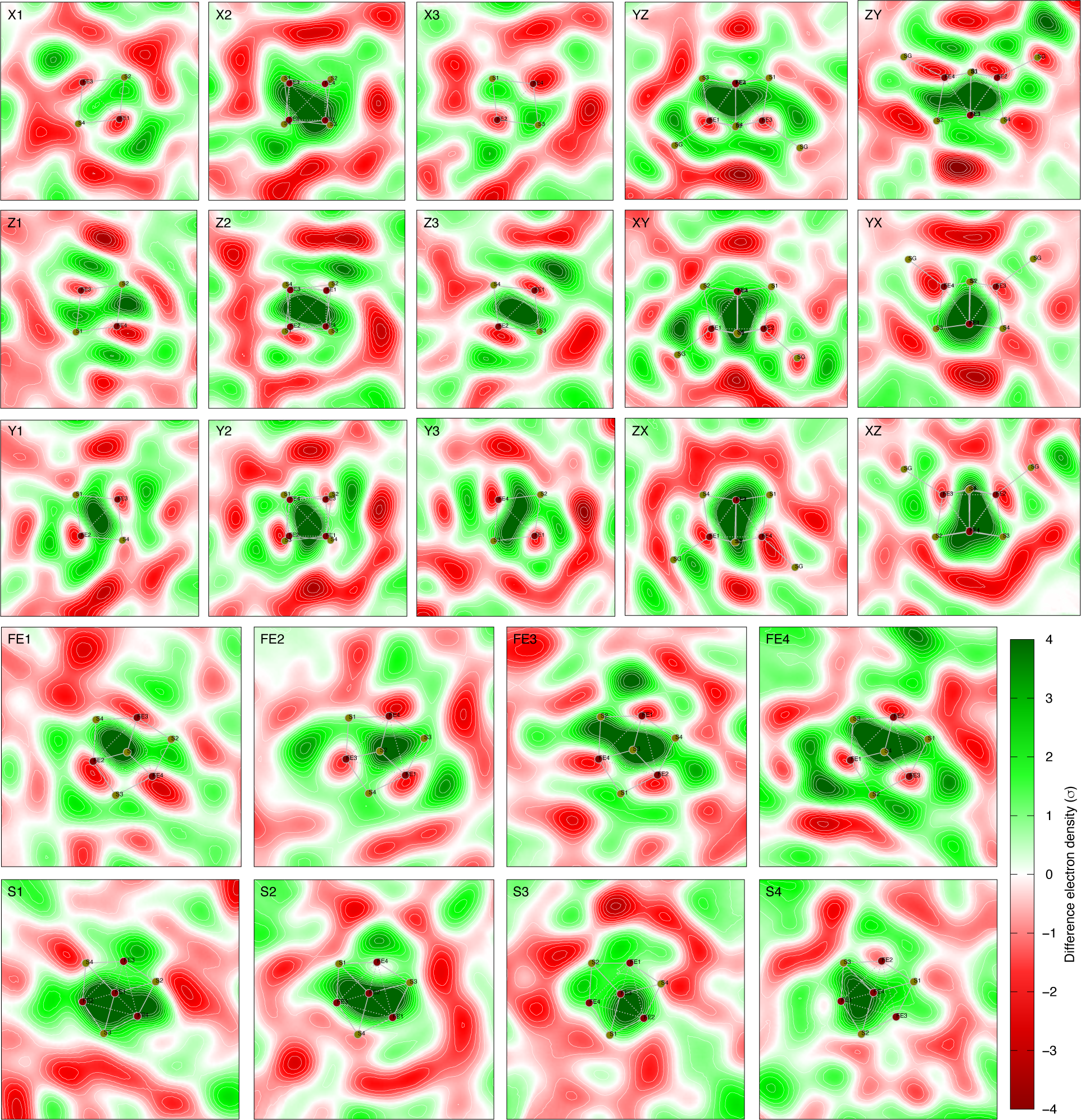
Raw difference map observed by illumination of 450 nm at 100 mA. See Figs. S1 and S13 for the definitions of cross sections. Each panel displays an integration within a slab of 0.4 Å thick around the cross section. This difference map is the only observation interpreted as [4Fe4S]^4+^ – [4Fe4S]^2+^. No layering of difference electron density is observed in this map. The corresponding reconstitution is shown in Figs. 4a and S22.

**Figure S22.**
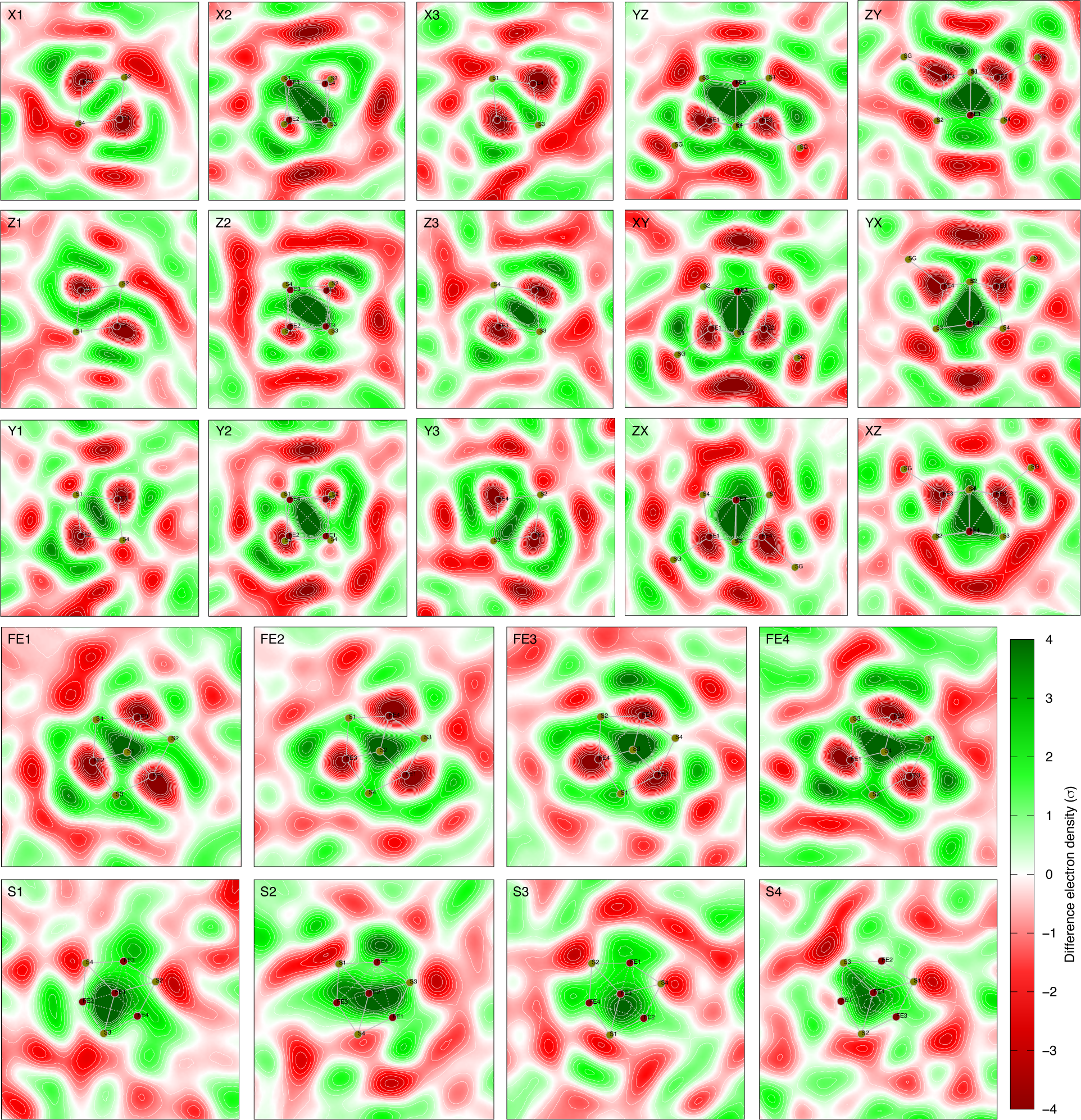
Reconstituted difference map of [4Fe4S]^4+^ – [4Fe4S]^2+^ = -640***U***_1_ + 525***U***_2_ - 300***U***_3_. See Figs. S1 and S13 for the definitions of cross sections. Each panel displays an integration within a slab of 0.4 Å thick around the cross section. The raw difference map corresponding to this reconstitution is shown in Fig. S21.

**Figure S23.**
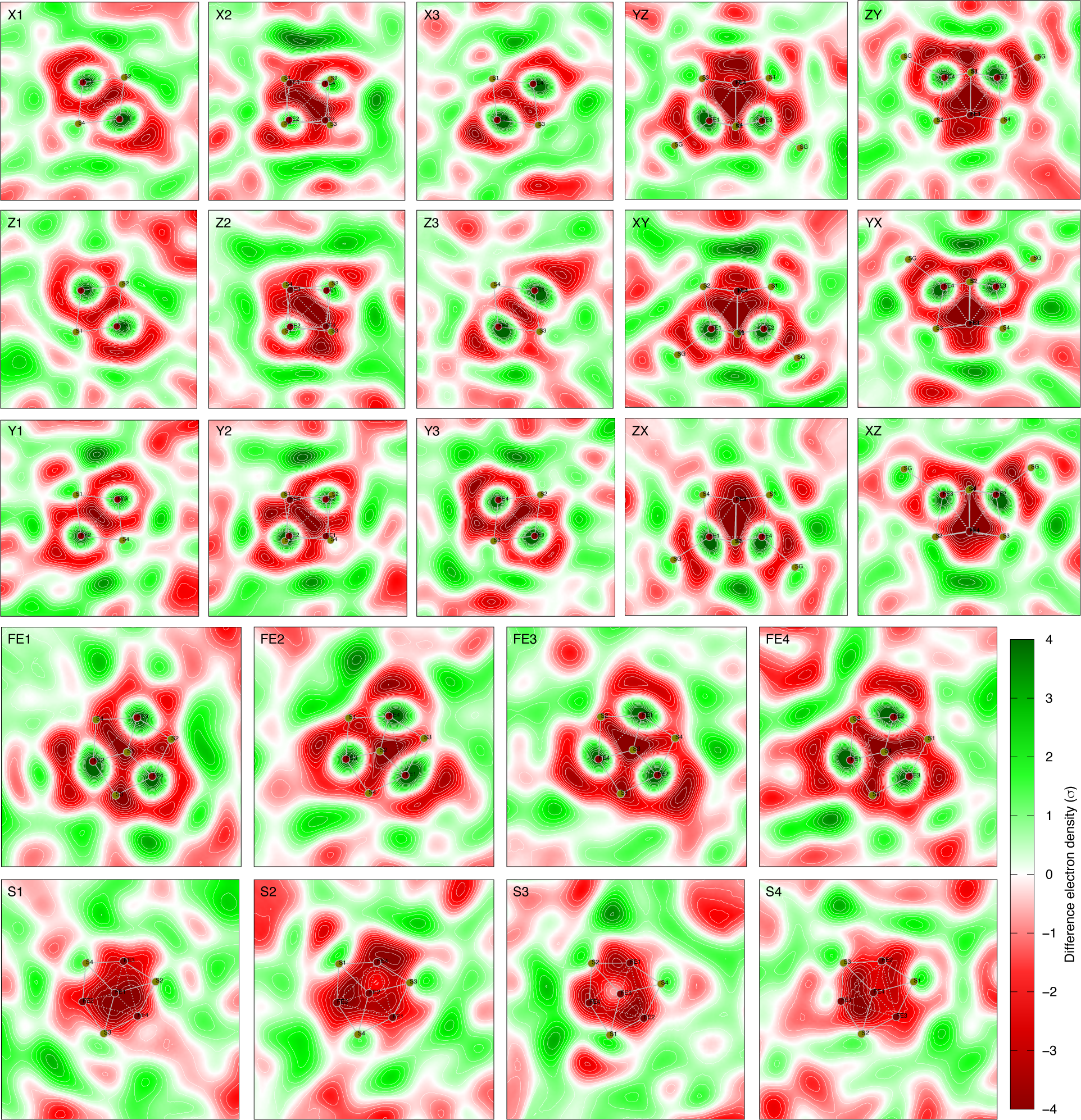
Raw difference map observed by illumination of 450 nm at 250 mA. See Figs. S1 and S13 for the definitions of cross sections. Each panel displays an integration within a slab of 0.4 Å thick around the cross section. This difference map in full tetrahedral symmetry T_d_ shows strong positive densities centered on all four irons and weaker positive densities associated with eight sulfurs. This map represents the observed [4Fe4S]^0^ – [4Fe4S]^2+^ without an expansion. The corresponding reconstitution is shown in Figs. 4c and S24.

**Figure S24.**
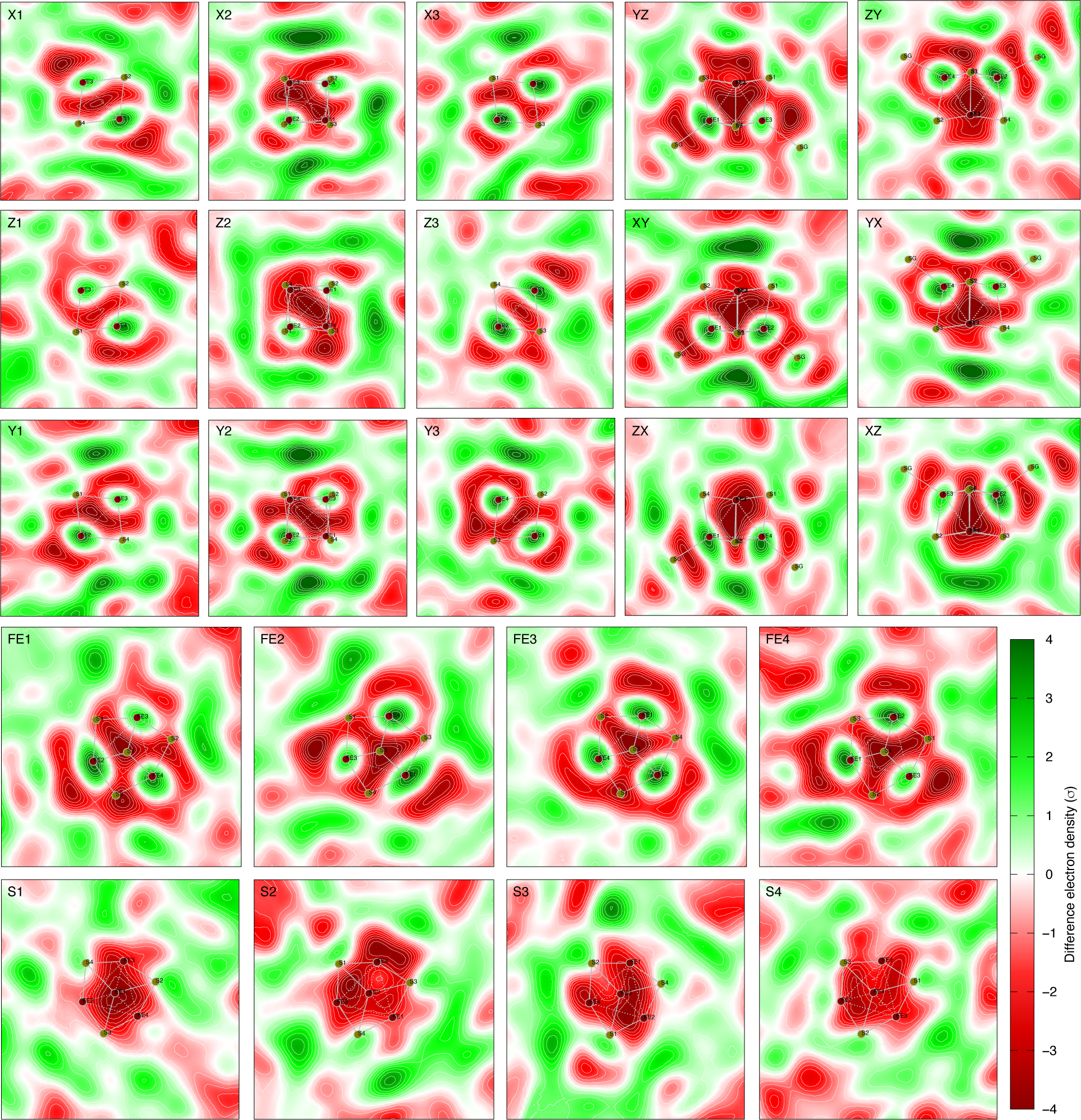
Reconstituted difference map of [4Fe4S]^0^ – [4Fe4S]^2+^ = 640***U***_1_ - 525***U***_2_ - 300***U***_3_ without an expansion. See Figs. S1 and S13 for the definitions of cross sections. Each panel displays an integration within a slab of 0.4 Å thick around the cross section. The linear combination 640***U***_1_ - 525***U***_2_ restores the tetrahedral symmetry of the neutral [4Fe4S]^0^. With a negative term of ***U***_3_, the positive densities are centered on the irons (Figs. 4c and S23) in contrast to a positive term of ***U***_3_ (Figs. 4d, S25, and S26). A raw difference map representing such neutral cluster [4Fe4S]^0^ with positive densities centered on the irons is observed by illumination of 450 nm light at 250 mA (Fig. S23).

**Figure S25.**
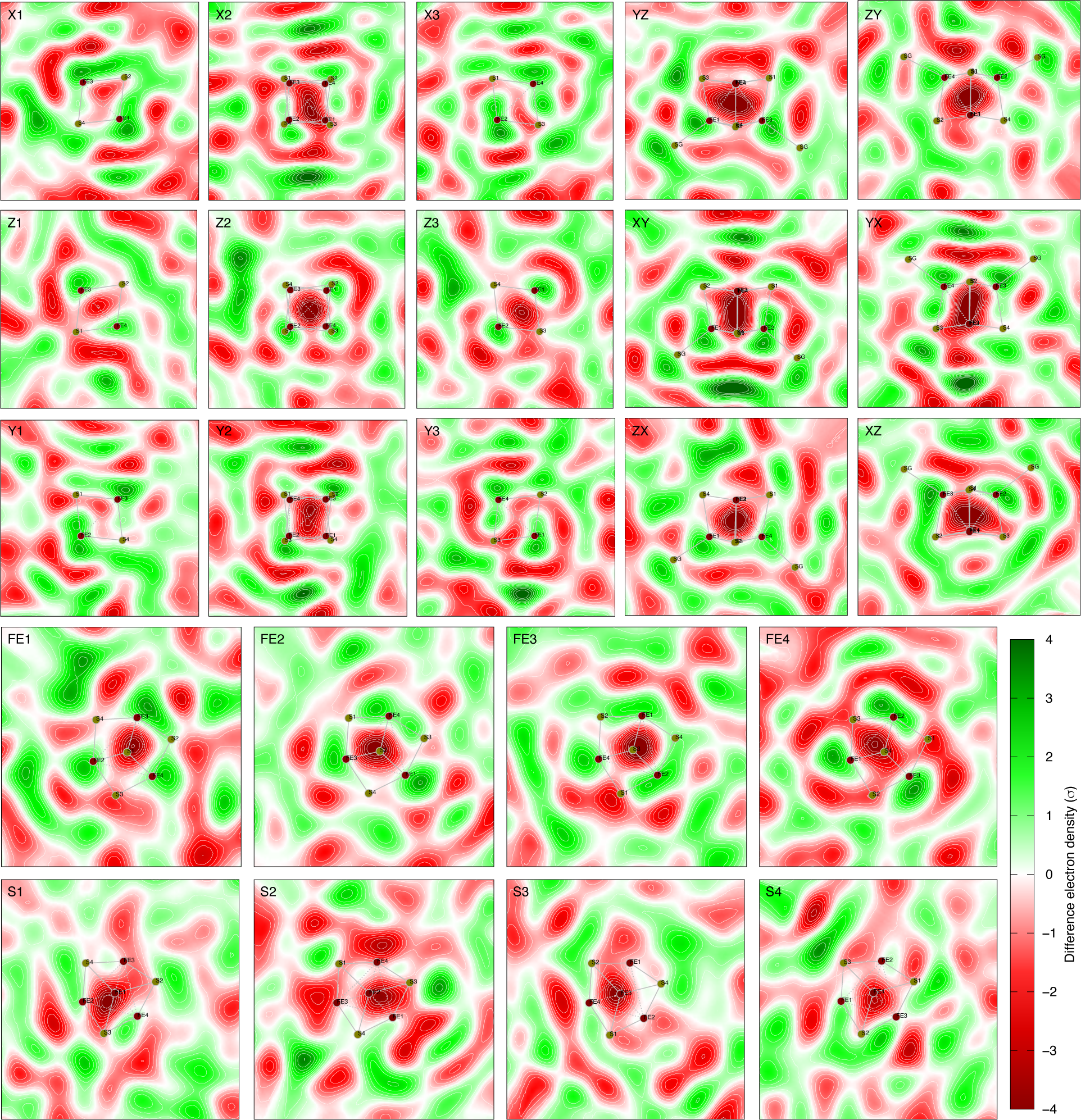
Raw difference map observed by illumination of 450 nm at 150 mA. See Figs. S1 and S13 for the definitions of cross sections. Each panel displays an integration within a slab of 0.4 Å thick around the cross section. This difference map in tetrahedral symmetry T_d_ shows positive densities near all four irons skewed outward. This map represents the observed [4Fe4S]^0^ – [4Fe4S]^2+^ with an expansion. The corresponding reconstitution is shown in Figs. 4d and S26.

**Figure S26.**
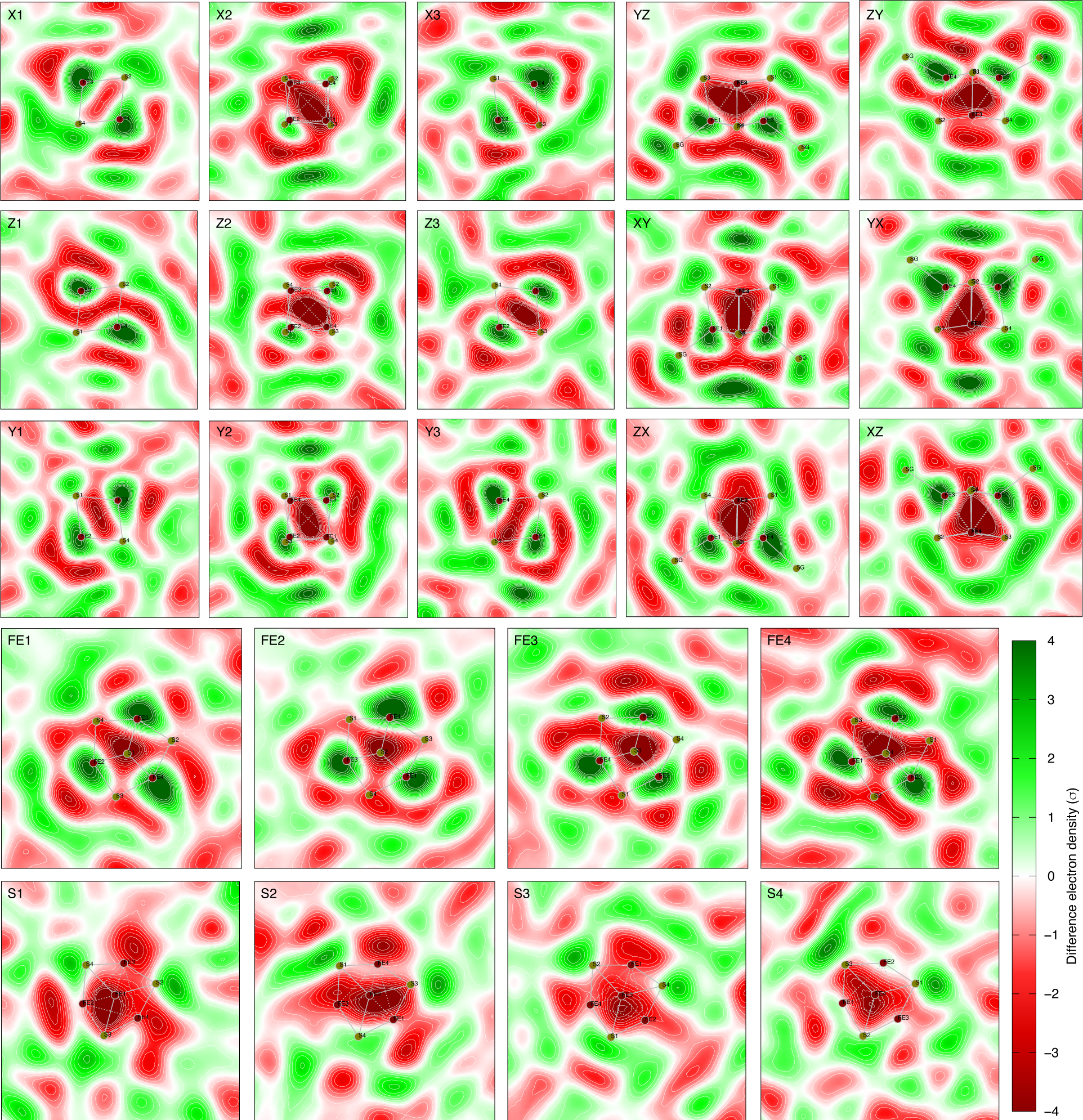
Reconstituted difference map [4Fe4S]^0^ – [4Fe4S]^2+^ = 640***U***_1_ - 525***U***_2_ + 600***U***_3_ with an expansion. See Figs. S1 and S13 for the definitions of cross sections. Each panel displays an integration within a slab of 0.4 Å thick around the cross section. The linear combination 640***U***_1_ - 525***U***_2_ restores the tetrahedral symmetry. With a large positive term of ***U***_3_, the positive densities on the irons skew outward away from the center of the [4Fe4S] cluster (Figs. 4d and S25), which seems to indicate an expansion of the all-ferrous cluster [4Fe4S]^0^. An observed all-ferrous cluster [4Fe4S]^0^ with an expansion is shown in Fig. S25.

**Figure S27.**
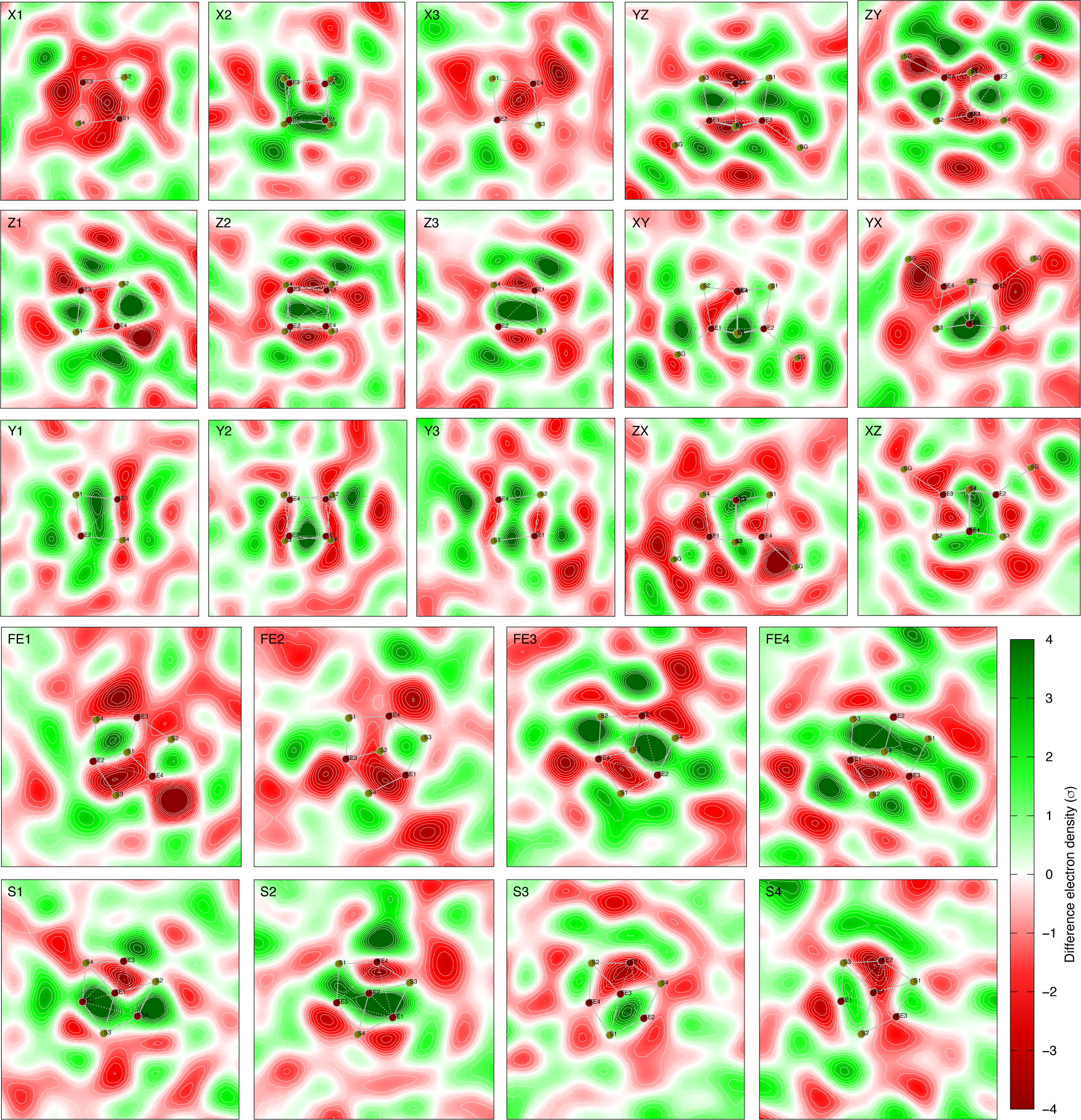
Raw difference map observed by illumination of 450 nm at 200 mA. See Figs. S1 and S13 for the definitions of cross sections. Each panel displays an integration within a slab of 0.4 Å thick around the cross section. An array of 3×2×2 positive peaks in the directions X, Y, and Z axes, respectively, are observed coexisting and interleaving with an array of 2×2×3 negative peaks. The most electron density gain can be seen in Z1 and Z3 cross sections while the most electron density loss is in X1 and X3 cross sections. This observation is nearly the negation of the other observed map shown in Fig. S29. The corresponding reconstitution is shown in Figs. 5a and S28.

**Figure S28.**
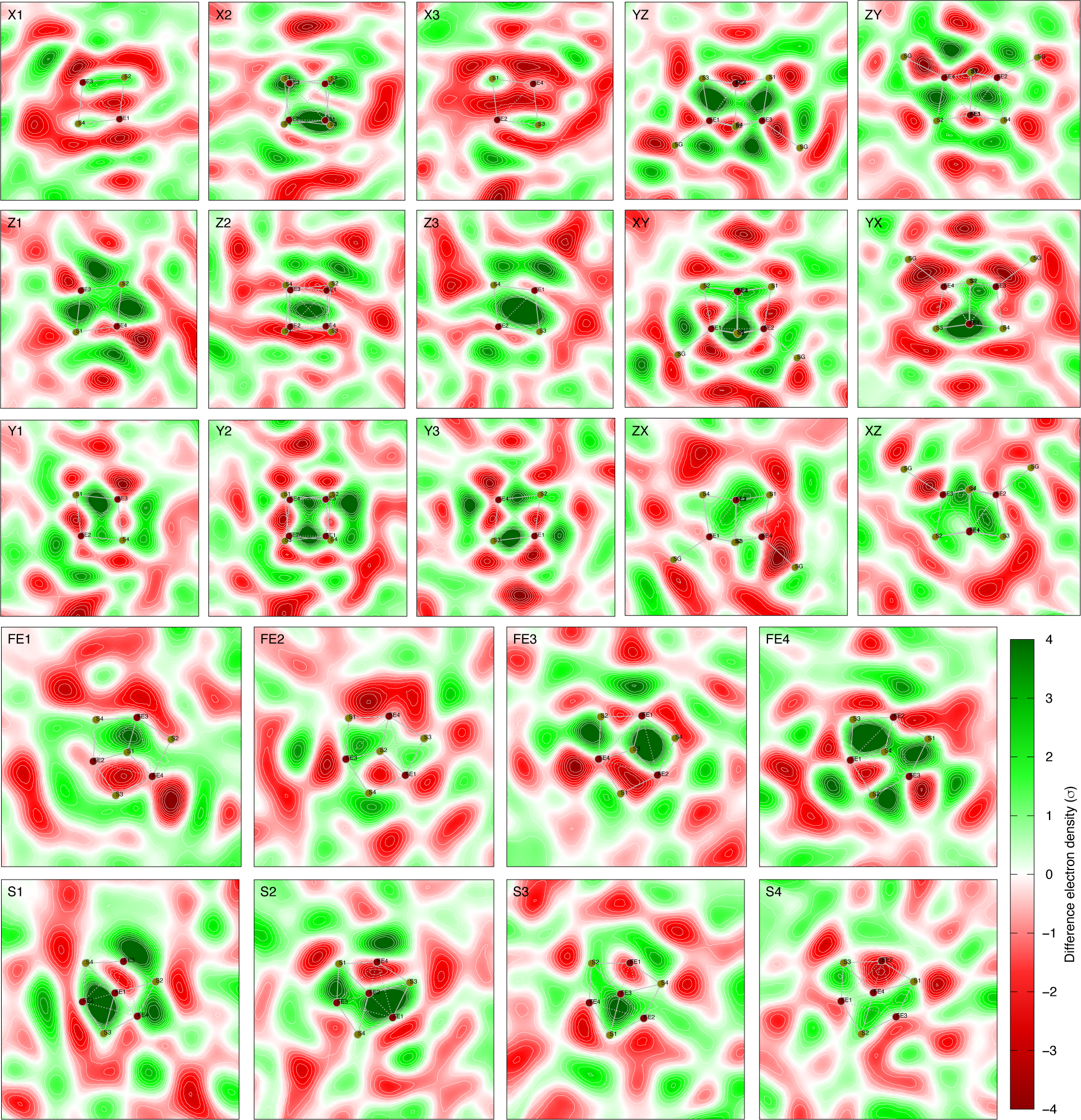
Reconstituted difference map 525***U***_2_ - 300***U***_3_. See Figs. S1 and S13 for the definitions of cross sections. Each panel displays an integration within a slab of 0.4 Å thick around the cross section. The corresponding observed difference map is shown in Fig. S27.

**Figure S29.**
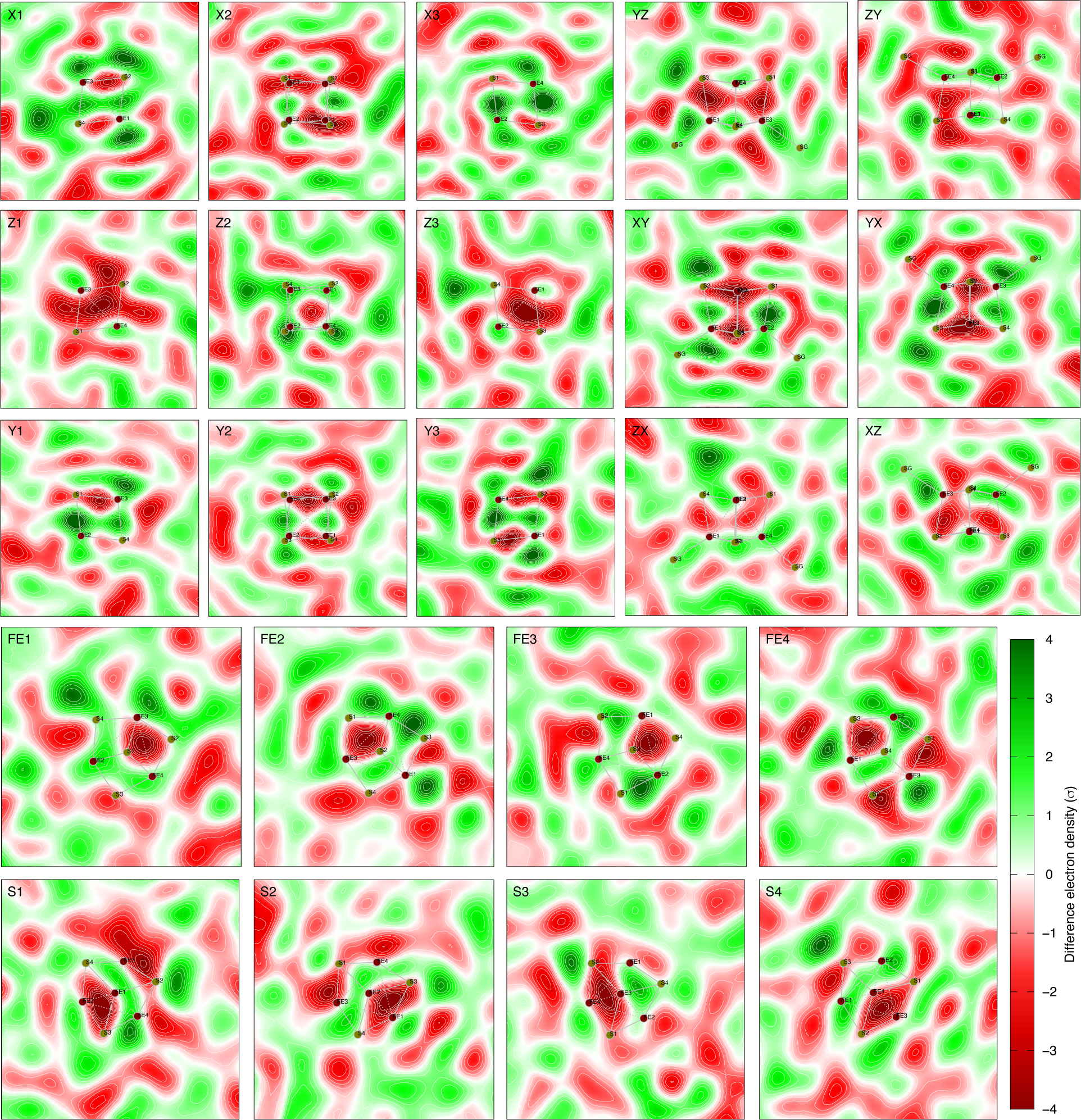
Raw difference map observed by illumination of 405 nm at 40 mA. See Figs. S1 and S13 for the definitions of cross sections. Each panel displays an integration within a slab of 0.4 Å thick around the cross section. An array of 3×2×2 negative peaks in the directions X, Y, and Z axes, respectively, are observed coexisting and interleaving with an array of 2×2×3 positive peaks. The most electron density gain can be seen in X1 and X3 cross sections while the most electron density loss is in Z1 and Z3 cross sections. This observation is nearly the negation of the other observed map shown in Fig. S27. The corresponding reconstitution is shown in Figs. 5bc and S30.

**Figure S30.**
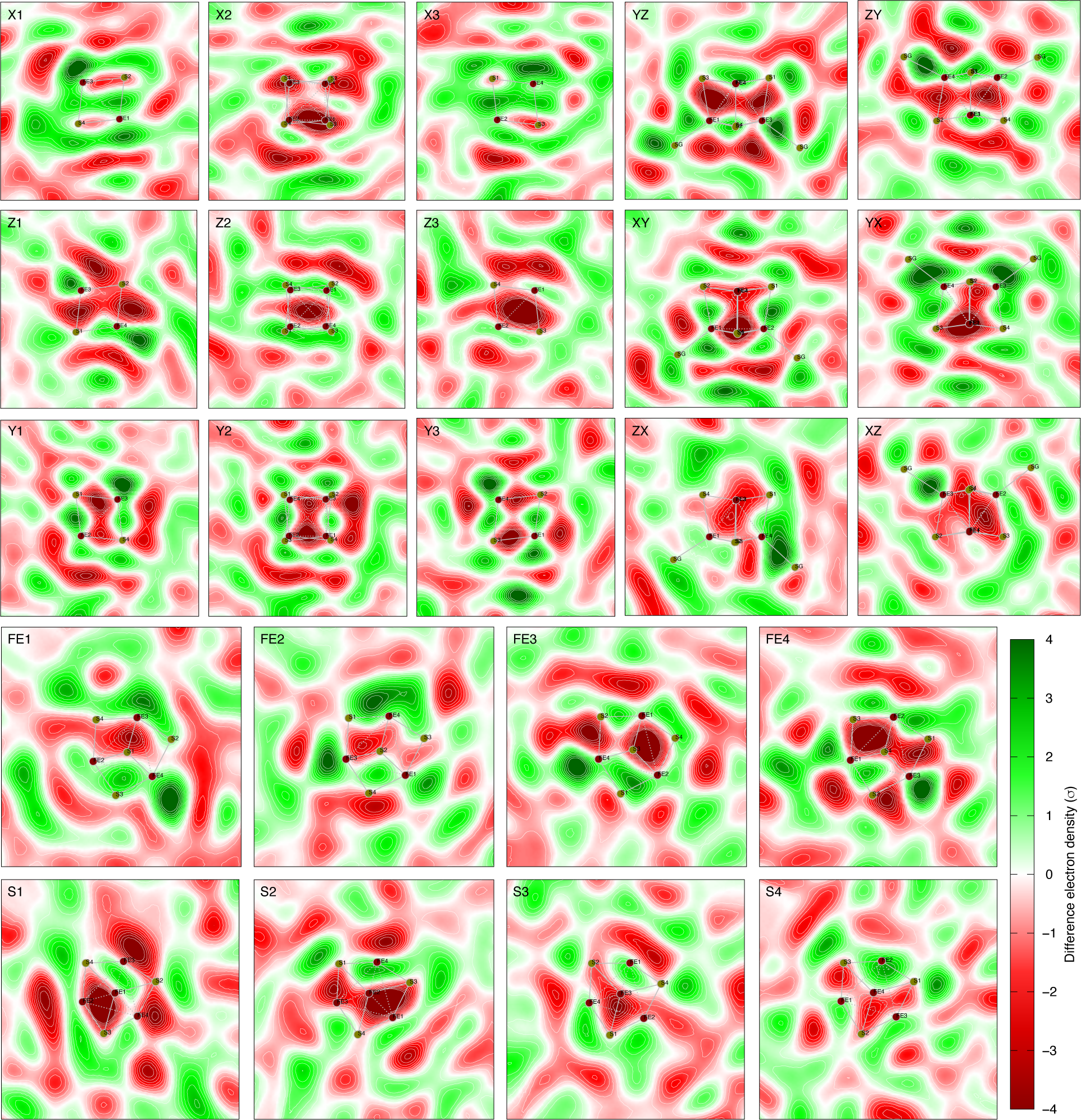
Reconstituted difference map -525***U***_2_ + 600***U***_3_. See Figs. S1 and S13 for the definitions of cross sections. Each panel displays an integration within a slab of 0.4 Å thick around the cross section. The corresponding observed difference map is shown in Fig. S29.

**Figure S31.**
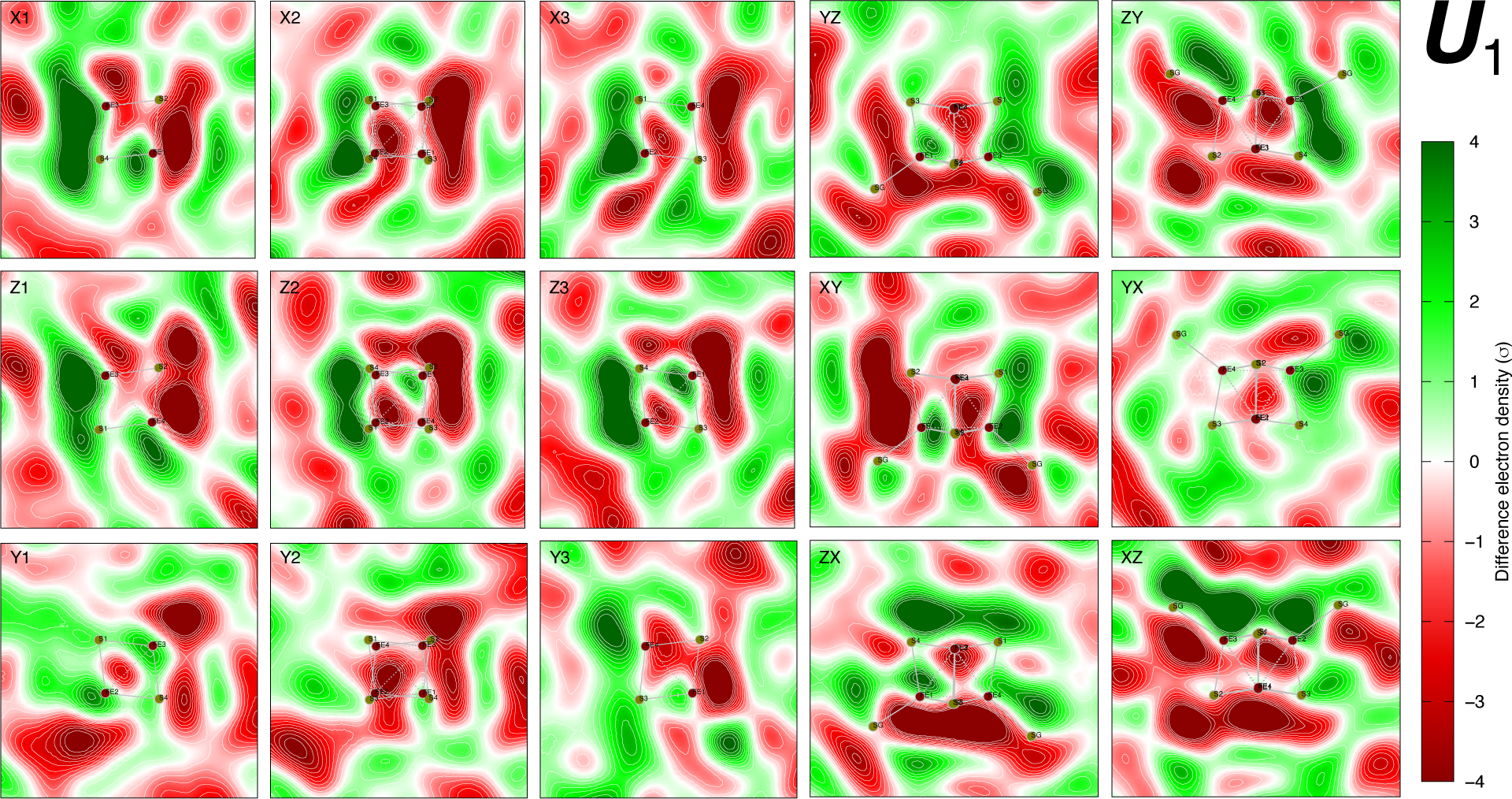
First component ***U***_1_ of the difference maps in temperature scan cryo-trapping. See Fig. S1 for the definitions of cross sections. Each panel displays an integration within a slab of 0.4 Å thick around the cross section. This component clearly shows a displacement of the [4Fe4S] cluster. See Figs. 6b and S10 for contour plots of this component.

**Figure S32.**
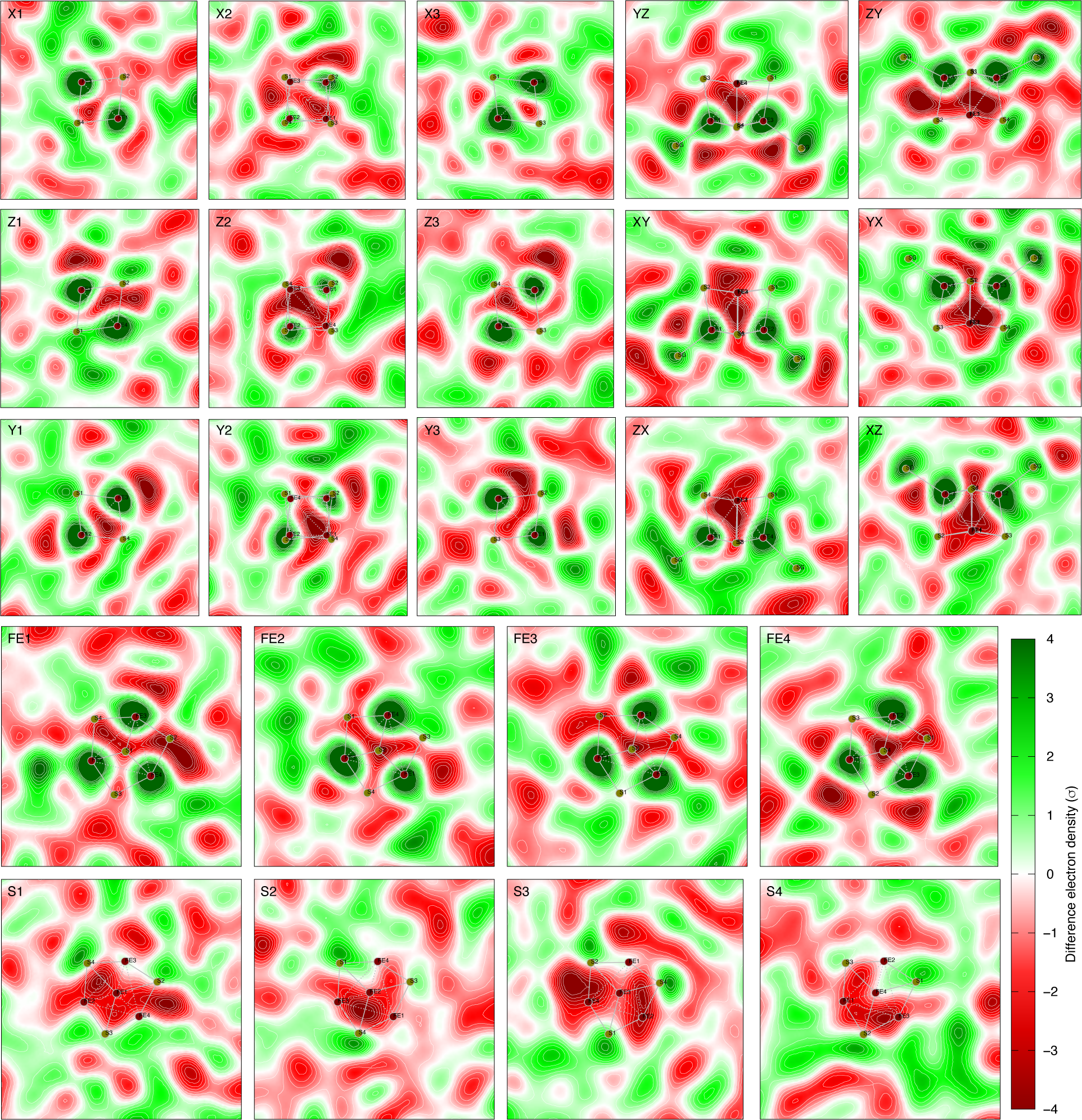
Third component ***U***_3_ of the difference maps in temperature scan cryo-trapping. See Figs. S1 and S13 for the definitions of cross sections. Each panel displays an integration within a slab of 0.4 Å thick around the cross section. This component follows the full tetrahedral symmetry T_d_. Strong spositive peaks are centered on the irons and other positive peaks are associated with eight sulfurs. See Fig. 6c for a contour plot of this component.

**Figure S33.**
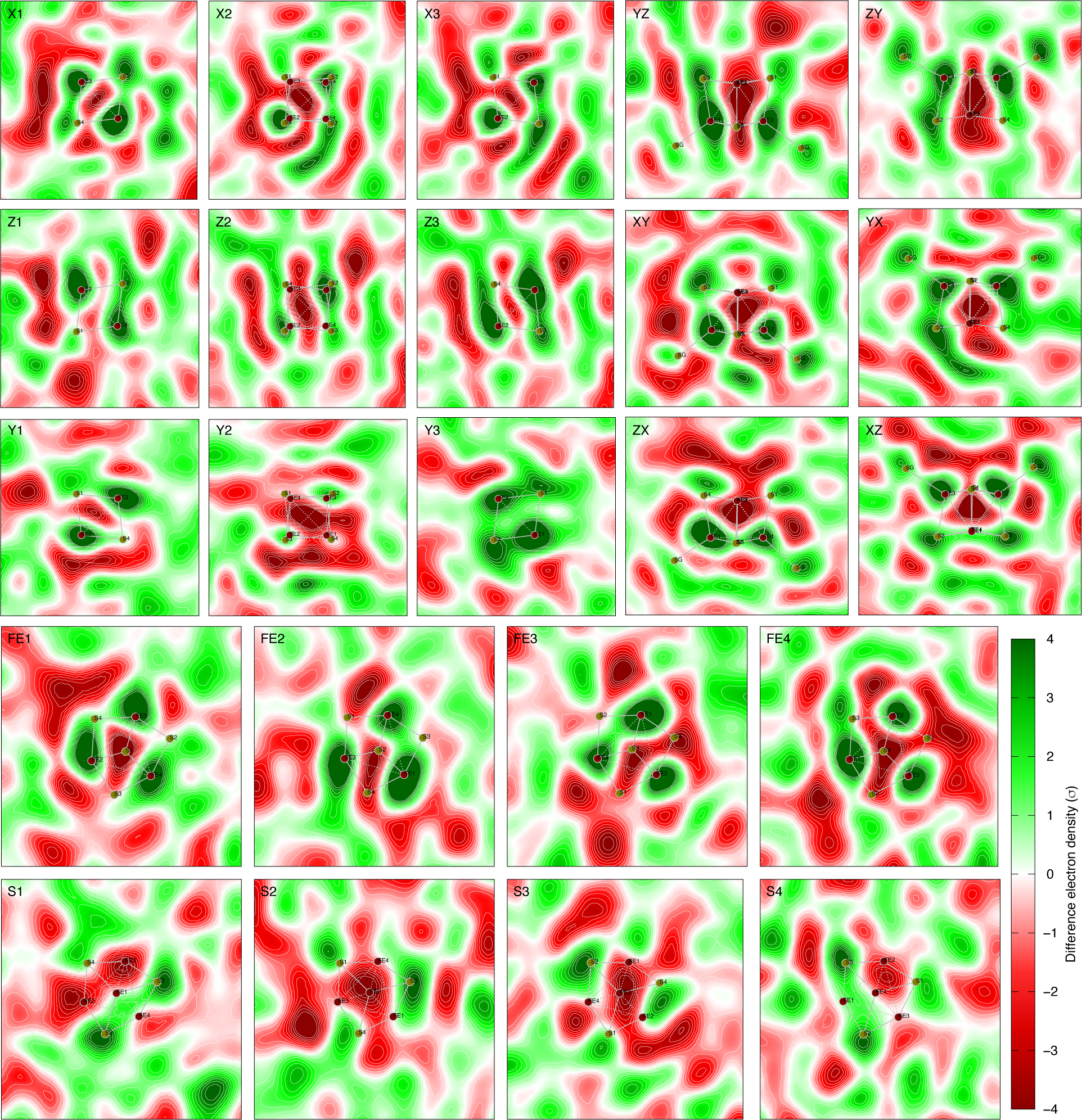
Sixth component ***U***_6_ of the difference maps in temperature scan cryo-trapping. See Figs. S1 and S13 for the definitions of cross sections. Each panel displays an integration within a slab of 0.4 Å thick around the cross section. Layering signal is along Y axis. Compare this component with the first component ***U***_1_ at room temperature (Fig. S13) and the reduced cluster [4Fe4S]^+^ (Figs. S17 and S18). See Fig. 6a for a contour plot of this component.

## References

Bandara S, Ren Z, Lu L, Xiaoli Z, Shin H, Zhao K-H, Yang X. 2017. Photoactivation mechanism of a carotenoid-based photoreceptor. Proc Natl Acad Sci U S A 114:6286–6291. doi:10.1073/pnas.1700956114

Baranovskiy AG, Siebler HM, Pavlov YI, Tahirov TH. 2018. Iron–sulfur clusters in DNA polymerases and primases of eukaryotesMethods in Enzymology. Elsevier. pp. 1–20. doi:10.1016/bs.mie.2017.09.003

Bauer TO, Graf D, Lamparter T, Schünemann V. 2014. Characterization of the photolyase-like iron sulfur protein PhrB from Agrobacterium tumefaciens by Mössbauer spectroscopy. Hyperfine Interact 226:445–449. doi:10.1007/s10751-013-0969-4

Beinert H. 2000. Iron-sulfur proteins: ancient structures, still full of surprises. JBIC J Biol Inorg Chem 5:2– 15. doi:10.1007/s007750050002

Beinert H. 1997. Iron-sulfur clusters: Nature’s modular, multipurpose structures. Science 277:653–659. doi:10.1126/science.277.5326.653

Biju LM, Wang C, Kang W, Tom IP, Kumarapperuma I, Yang X, Ren Z. 2022. On-chip crystallization and large-scale serial diffraction at room temperature. J Vis Exp 181:e63022. doi:10.3791/63022

Dikbas UM, Tardu M, Canturk A, Gul S, Ozcelik G, Baris I, Ozturk N, Kavakli IH. 2019. Identification and characterization of a new class of (6–4) photolyase from Vibrio cholerae. Biochemistry 58:4352– 4360. doi:10.1021/acs.biochem.9b00766

Emsley P, Cowtan K. 2004. Coot: model-building tools for molecular graphics. Acta Crystallogr D Biol Crystallogr 60:2126–2132. doi:10.1107/S0907444904019158

Fontecave M. 2006. Iron-sulfur clusters: ever-expanding roles. Nat Chem Biol 2:171–174. doi:10.1038/nchembio0406-171

Geisselbrecht Y, Frühwirth S, Schroeder C, Pierik AJ, Klug G, Essen L. 2012. CryB from Rhodobacter sphaeroides: a unique class of cryptochromes with new cofactors. EMBO Rep 13:223–229. doi:10.1038/embor.2012.2

Graber T, Anderson S, Brewer H, Chen Y-S, Cho HS, Dashdorj N, Henning RW, Kosheleva I, Macha G, Meron M, Pahl R, Ren Z, Ruan S, Schotte F, Šrajer V, Viccaro PJ, Westferro F, Anfinrud P, Moffat K. 2011. BioCARS: a synchrotron resource for time-resolved X-ray science. J Synchrotron Radiat 18:658–670. doi:10.1107/S0909049511009423

Graf D, Wesslowski J, Ma H, Scheerer P, Krauß N, Oberpichler I, Zhang F, Lamparter T. 2015. Key amino acids in the bacterial (6-4) photolyase PhrB from Agrobacterium fabrum. PLOS ONE 10:e0140955. doi:10.1371/journal.pone.0140955

Holm RH, Kennepohl P, Solomon EI. 1996. Structural and functional aspects of metal sites in biology. Chem Rev 96:2239–2314. doi:10.1021/cr9500390

Johnson DC, Dean DR, Smith AD, Johnson MK. 2005. Structure, function, and formation of biological iron-sulfur clusters. Annu Rev Biochem 74:247–281. doi:10.1146/annurev.biochem.74.082803.133518

Jozwiakowski SK, Kummer S, Gari K. 2019. Human DNA polymerase delta requires an iron–sulfur cluster for high-fidelity DNA synthesis. Life Sci Alliance 2:e201900321. doi:10.26508/lsa.201900321

Kavakli IH, Ozturk N, Gul S. 2019. DNA repair by photolyasesAdvances in Protein Chemistry and Structural Biology. Elsevier. pp. 1–19. doi:10.1016/bs.apcsb.2018.10.003

Khodour Y, Kaguni LS, Stiban J. 2019. Iron–sulfur clusters in nucleic acid metabolism: Varying roles of ancient cofactorsThe Enzymes. Elsevier. pp. 225–256. doi:10.1016/bs.enz.2019.08.003

Liu L, Huang M. 2015. Essential role of the iron-sulfur cluster binding domain of the primase regulatory subunit Pri2 in DNA replication initiation. Protein Cell 6:194–210. doi:10.1007/s13238-015-0134-8

Marizcurrena JJ, Lamparter T, Castro-Sowinski S. 2020. A (6-4)-photolyase from the Antarctic bacterium Sphingomonas sp. UV9: recombinant production and in silico features. Extremophiles. doi:10.1007/s00792-020-01202-z

Massa L, Huang L, Karle J. 1995. Quantum crystallography and the use of kernel projector matrices. Int J Quantum Chem 56:371–384. doi:10.1002/qua.560560841

Moula G, Matsumoto T, Miehlich ME, Meyer K, Tatsumi K. 2018. Synthesis of an all-ferric cuboidal iron– sulfur cluster [Fe^III^_4_S_4_(SAr)_4_]. Angew Chem Int Ed 57:11594–11597. doi:10.1002/anie.201803679

Musgrave KB, Angove HC, Burgess BK, Hedman B, Hodgson KO. 1998. All-ferrous titanium(III) citrate reduced Fe protein of nitrogenase: An XAS study of electronic and metrical structure. J Am Chem Soc 120:5325–5326. doi:10.1021/ja980598z

Netz DJA, Stith CM, Stümpfig M, Köpf G, Vogel D, Genau HM, Stodola JL, Lill R, Burgers PMJ, Pierik AJ. 2012. Eukaryotic DNA polymerases require an iron-sulfur cluster for the formation of active complexes. Nat Chem Biol 8:125–132. doi:10.1038/nchembio.721

Noodleman L, Case DA. 1992. Density-functional theory of spin polarization and spin coupling in iron— sulfur clusters In: Cammack R, editor. Advances in Inorganic Chemistry. Academic Press. pp. 423–470. doi:10.1016/S0898-8838(08)60070-7

Noodleman L, Lovell T, Liu T, Himo F, Torres RA. 2002. Insights into properties and energetics of iron– sulfur proteins from simple clusters to nitrogenase. Curr Opin Chem Biol 6:259–273. doi:10.1016/S1367-5931(02)00309-5

Noodleman L, Peng CY, Case DA, Mouesca J-M. 1995. Orbital interactions, electron delocalization and spin coupling in iron-sulfur clusters. Coord Chem Rev 144:199–244. doi:10.1016/0010-8545(95)07011-L

Oberpichler I, Pierik AJ, Wesslowski J, Pokorny R, Rosen R, Vugman M, Zhang F, Neubauer O, Ron EZ, Batschauer A, Lamparter T. 2011. A photolyase-like protein from Agrobacterium tumefaciens with an iron-sulfur cluster. PLoS ONE 6:e26775. doi:10.1371/journal.pone.0026775

O’Brien E, Holt ME, Thompson MK, Salay LE, Ehlinger AC, Chazin WJ, Barton JK. 2017. The [4Fe4S] cluster of human DNA primase functions as a redox switch using DNA charge transport. Science 355:eaag1789. doi:10.1126/science.aag1789

Otwinowski Z, Minor W. 1997. Processing of X-ray diffraction data collected in oscillation modeMethods in Enzymology. Elsevier. pp. 307–326. doi:10.1016/S0076-6879(97)76066-X

Polkosnik W, Matta CF, Huang L, Massa L. 2019. Fast quantum crystallography. Int J Quantum Chem 119. doi:10.1002/qua.25986

Ren Z. 2022. Photoinduced isomerization sampling of retinal in bacteriorhodopsin. PNAS Nexus 1. doi:10.1093/pnasnexus/pgac103

Ren Z. 2021. Directional proton conductance in bacteriorhodopsin is driven by concentration gradient, not affinity gradient. bioRxiv. doi:10.1101/2021.10.04.463074

Ren Z. 2019. Ultrafast structural changes decomposed from serial crystallographic data. J Phys Chem Lett 10:7148–7163. doi:10.1021/acs.jpclett.9b02375

Ren Z. 2017. Single crystal quartz chips for protein crystallization and X-ray diffraction data collection and related methods. US patent 9632042. 9,632,042.

Ren Z. 2016. Molecular events during translocation and proofreading extracted from 200 static structures of DNA polymerase. Nucleic Acids Res 6:1–13. doi:10.1093/nar/gkw555

Ren Z, Ayhan M, Bandara S, Bowatte K, Kumarapperuma I, Gunawardana S, Shin H, Wang C, Zeng X, Yang X. 2018. Crystal-on-crystal chips for in situ serial diffraction at room temperature. Lab Chip 18:2246–2256. doi:10.1039/C8LC00489G

Ren Z, Bourgeois D, Helliwell JR, Moffat K, Srajer V, Stoddard BL. 1999. Laue crystallography: coming of age. J Synchrotron Rad 6:891–917. doi:10.1107/S0909049599006366

Ren Z, Chan PWY, Moffat K, Pai EF, Royer WE, Šrajer V, Yang X. 2013. Resolution of structural heterogeneity in dynamic crystallography. Acta Cryst D69:946–959. doi:10.1107/S0907444913003454

Ren Z, Kang W, Gunawardana S, Bowatte K, Thoulass K, Kaeser G, Krauß N, Lamparter T, Yang X. 2022. Dynamic interplays between three redox cofactors in DNA photolyase PhrB. Cell Rep Phys Sci. doi:10.2139/ssrn.4194951

Ren Z, Wang C, Shin H, Bandara S, Kumarapperuma I, Ren MY, Kang W, Yang X. 2020. An automated platform for in situ serial crystallography at room temperature. IUCrJ 7:1009–1018. doi:10.1107/S2052252520011288

Sancar A. 2003. Structure and function of DNA photolyase and cryptochrome blue-light photoreceptors. Chem Rev 103:2203–2238. doi:10.1021/cr0204348

Shin H, Ren Z, Zeng X, Bandara S, Yang X. 2019. Structural basis of molecular logic OR in a dual-sensor histidine kinase. Proc Natl Acad Sci 116:19973–19982. doi:10.1073/pnas.1910855116

Vanoni MA. 2021. Iron-sulfur flavoenzymes: the added value of making the most ancient redox cofactors and the versatile flavins work together. Open Biol 11:210010. doi:10.1098/rsob.210010

Velazquez-Garcia J de J, Basuroy K, Storozhuk D, Wong J, Demeshko S, Meyer F, Henning R, Techert S. 2022. Metal-to-metal communication during the spin state transition of a [2×2] Fe(II) metallogrid at equilibrium and out-of-equilibrium conditions. Dalton Trans 51:6036–6045. doi:10.1039/D1DT04255F

Voityuk AA. 2010. Electron transfer between [4Fe–4S] clusters. Chem Phys Lett 495:131–134. doi:10.1016/j.cplett.2010.06.057

Wang X-B, Niu S, Yang X, Ibrahim SK, Pickett CJ, Ichiye T, Wang L-S. 2003. Probing the intrinsic electronic structure of the cubane [4Fe−4S] cluster: Nature’s favorite cluster for electron transfer and storage. J Am Chem Soc 125:14072–14081. doi:10.1021/ja036831x

Yang X, Ren Z, Kuk J, Moffat K. 2011. Temperature-scan cryocrystallography reveals reaction intermediates in bacteriophytochrome. Nature 479:428–432. doi:10.1038/nature10506

Yoo SJ, Angove HC, Burgess BK, Hendrich MP, Münck E. 1999. Mössbauer and integer-spin EPR studies and spin-coupling analysis of the [4Fe-4S]^0^ cluster of the Fe protein from Azotobacter vinelandii nitrogenase. J Am Chem Soc 121:2534–2545. doi:10.1021/ja9837405

Zeng X, Ren Z, Wu Q, Fan J, Peng P-P, Tang K, Zhang R, Zhao K-H, Yang X. 2015. Dynamic crystallography reveals early signaling events in ultraviolet photoreceptor UVR8. Nat Plants 1:14006. doi:10.1038/NPLANTS.2014.6

Zhang F, Scheerer P, Oberpichler I, Lamparter T, Krauss N. 2013. Crystal structure of a prokaryotic (6-4) photolyase with an Fe-S cluster and a 6,7-dimethyl-8-ribityllumazine antenna chromophore. Proc Natl Acad Sci 110:7217–7222. doi:10.1073/pnas.1302377110

Zhang M, Wang L, Zhong D. 2017. Photolyase: Dynamics and electron-transfer mechanisms of DNA repair. Arch Biochem Biophys 632:158–174. doi:10.1016/j.abb.2017.08.007

Zuo JM, Kim M, O’Keeffe M, Spence JCH. 1999. Direct observation of d-orbital holes and Cu–Cu bonding in Cu2O. Nature 401:49–52. doi:10.1038/43403

## Supplementary References

Biju, L.M., Wang, C., Kang, W., Tom, I.P., Kumarapperuma, I., Yang, X., and Ren, Z. (2022). On-chip crystallization and large-scale serial diffraction at room temperature. J Vis Exp 181, e63022. https://doi.org/10.3791/63022.

Graber, T., Anderson, S., Brewer, H., Chen, Y.-S., Cho, H.S., Dashdorj, N., Henning, R.W., Kosheleva, I., Macha, G., Meron, M., et al. (2011). BioCARS: a synchrotron resource for time-resolved X-ray science. J. Synchrotron Radiat. 18, 658–670. https://doi.org/10.1107/S0909049511009423.

Henry, E.R., and Hofrichter, J. (1992). Singular value decomposition: Application to analysis of experimental data. In Numerical Computer Methods, (Academic Press), pp. 129–192.

Oberpichler, I., Pierik, A.J., Wesslowski, J., Pokorny, R., Rosen, R., Vugman, M., Zhang, F., Neubauer, O., Ron, E.Z., Batschauer, A., et al. (2011). A photolyase-like protein from Agrobacterium tumefaciens with an iron-sulfur cluster. PLoS ONE 6, e26775. https://doi.org/10.1371/journal.pone.0026775.

Otwinowski, Z., and Minor, W. (1997). Processing of X-ray diffraction data collected in oscillation mode. In Methods in Enzymology, (Elsevier), pp. 307–326.

Ren, Z. (2006). Precognition user guide with reference and tutorials http://researchgate.net/publication/259441273_Precognition_User_Guide_with_Reference_and_Tutorials?ev=prf_pub.

Ren, Z. (2008). How to process Laue data using Precognition http://researchgate.net/publication/259441538_How_to_Process_Laue_Data_Using_Precognition?ev=prf_pub.

Ren, Z. (2016). Molecular events during translocation and proofreading extracted from 200 static structures of DNA polymerase. Nucleic Acids Res. 6, 1–13. https://doi.org/10.1093/nar/gkw555.

Ren, Z. (2019). Ultrafast structural changes decomposed from serial crystallographic data. J. Phys. Chem. Lett. 10, 7148–7163. https://doi.org/10.1021/acs.jpclett.9b02375.

Ren, Z., Bourgeois, D., Helliwell, J.R., Moffat, K., Srajer, V., and Stoddard, B.L. (1999). Laue crystallography: coming of age. J Synchrotron Rad 6, 891–917. https://doi.org/10.1107/S0909049599006366.

Ren, Z., Perman, B., Srajer, V., Teng, T.-Y., Pradervand, C., Bourgeois, D., Schotte, F., Ursby, T., Kort, R., Wulff, M., et al. (2001). A molecular movie at 1.8 Å resolution displays the photocycle of photoactive yellow protein, a eubacterial blue-light receptor, from nanoseconds to seconds. Biochemistry 40, 13788– 13801. https://doi.org/10.1021/bi0107142.

Ren, Z., Chan, P.W.Y., Moffat, K., Pai, E.F., Royer, W.E., Šrajer, V., and Yang, X. (2013). Resolution of structural heterogeneity in dynamic crystallography. Acta Cryst D69, 946–959. https://doi.org/10.1107/S0907444913003454.

Ren, Z., Ayhan, M., Bandara, S., Bowatte, K., Kumarapperuma, I., Gunawardana, S., Shin, H., Wang, C., Zeng, X., and Yang, X. (2018). Crystal-on-crystal chips for in situ serial diffraction at room temperature. Lab. Chip 18, 2246–2256. https://doi.org/10.1039/C8LC00489G.

Ren, Z., Wang, C., Shin, H., Bandara, S., Kumarapperuma, I., Ren, M.Y., Kang, W., and Yang, X. (2020). An automated platform for in situ serial crystallography at room temperature. IUCrJ 7, 1009–1018. https://doi.org/10.1107/S2052252520011288.

Ren, Z., Kang, W., Gunawardana, S., Bowatte, K., Thoulass, K., Kaeser, G., Krauß, N., Lamparter, T., and Yang, X. (2022). Dynamic interplays between three redox cofactors in DNA photolyase PhrB. Cell Rep. Phys. Sci. https://doi.org/10.2139/ssrn.4194951.

Schmidt, M., Rajagopal, S., Ren, Z., and Moffat, K. (2003). Application of singular value decomposition to the analysis of time-resolved macromolecular X-ray data. Biophys. J. 84, 2112–2129. https://doi.org/10.1016/S0006-3495(03)75018-8.

Šrajer, V., Ren, Z., Teng, T.-Y., Schmidt, M., Ursby, T., Bourgeois, D., Pradervand, C., Schildkamp, W., Wulff, M., and Moffat, K. (2001). Protein conformational relaxation and ligand migration in myoglobin: A nanosecond to millisecond molecular movie from time-resolved Laue X-ray diffraction. Biochemistry 40, 13802–13815. https://doi.org/10.1021/bi010715u.

Ursby, T., and Bourgeois, D. (1997). Improved estimation of structure-factor difference amplitudes from poorly accurate data. Acta Crystallogr. A 53, 564–575. https://doi.org/10.1107/S0108767397004522.

Zhang, F., Scheerer, P., Oberpichler, I., Lamparter, T., and Krauss, N. (2013). Crystal structure of a prokaryotic (6-4) photolyase with an Fe-S cluster and a 6,7-dimethyl-8-ribityllumazine antenna chromophore. Proc. Natl. Acad. Sci. 110, 7217–7222. https://doi.org/10.1073/pnas.1302377110.

